# Neurofibromin Regulates Sleep Consolidation via Cholinergic Neurons in a *Drosophila* Model of Neurofibromatosis 1

**DOI:** 10.64898/2026.10.20.760518

**Authors:** Kalliopi Atsoniou, Eirini-Maria Georganta, Kathrin Pfeifer, Ruth Palmer, Martina Samiotaki, Efthimios MC Skoulakis

## Abstract

Neurofibromatosis type 1 (NF1) patients frequently present with fragmented, poor-quality sleep, but the neural circuits and molecular mechanisms underlying this deficit remain undefined. Using Drosophila, we show that loss of the NF1 ortholog dNf1 recapitulates the human phenotype: null mutants exhibit markedly reduced total sleep with blunted nighttime consolidation, reflected in fewer and shorter sleep bouts and elevated inter-bout activity, a recessive phenotype reproduced by independent null alleles and by pan-neuronal RNAi knockdown. Despite dNf1’s broad expression across the adult CNS, including the mushroom bodies, ellipsoid body, dorsal fan-shaped body, and GABAergic neurons, cell-type-restricted knockdown showed the protein is dispensable for sleep regulation in each of these populations. Instead, dNf1 loss specifically in cholinergic neurons, the most abundant neuronal class in the fly brain, was sufficient to reproduce the deficient, fragmented sleep of null mutants, and restoring wild-type dNf1 to cholinergic neurons of null animals was both necessary and sufficient to rescue sleep. Mechanistically, cholinergic expression of the GAP-related domain (GRD) alone, which normalizes elevated Ras/MAPK signaling, fully restored sleep in nulls, whereas a catalytically dead GRD variant did not, establishing Ras/MAPK dysregulation in cholinergic neurons as the proximate cause. Two disease-associated missense mutations, *dNf1^C1045Y^*and *dNf1^R1861C^*, produced distinct, dose-graded sleep phenotypes. *dNf1^C1045Y^* showed severe deficits resembling the null and was resistant to pan-neuronal, but responded to cholinergic neuron-restricted rescue, consistent with a dominant-negative effect on dNf1 dimerization. In contrast, *dNf1^R1861C^*presented a milder phenotype that was largely reversible by wild-type dNf1 but only partially by the GRD domain, implicating an additional, GRD-independent mechanism. Collectively, these findings define cholinergic neurons as the key circuit through which Neurofibromin regulates sleep consolidation via Ras/MAPK signaling and reveal mutation-specific mechanisms that mirror the phenotypic heterogeneity seen in NF1 patients, with implications for genotype-informed understanding of sleep disturbance in this disorder.

## INTRODUCTION

Neurofibromatosis type 1 (NF1) (OMIM#162200) is a multi-symptomatic disorder caused by inherited or *de novo* mutations in the *NF1* tumor suppressor gene (Bergqvist, Servy et al. 2020, Moodley and Lopez 2024). The gene encodes neurofibromin (Nf1), a ∼2800 aa protein that regulates Ras/MAPK, Raf/MEK/ERK, PI3K/AKT/mTOR, and cAMP/PKA signaling (Martin, Viskochil et al. 1990, Tong, Hannan et al. 2002, Harrisingh and Lloyd 2004, Johannessen, Reczek et al. 2005, Bergoug, Doudeau et al. 2020). NF1 presents with a diverse clinical spectrum, including cutaneous neurofibromas, café-au-lait macules, Lisch nodules, elevated neoplasia risk, and cognitive-behavioral deficits such as impaired learning and memory. Disrupted sleep quality and patterning are also common neurological morbidities in pediatric NF1 (Marana Perez, Duat Rodriguez et al. 2015), manifesting as reduced total sleep time, increased awakenings, and parasomnia that together produce sleep fragmentation (Johnson, Wiggs et al. 2005, Licis, Vallorani et al. 2013, Marana Perez, Duat Rodriguez et al. 2015, Carotenuto, Messina et al. 2023).

In a cross-sectional study of 129 children with NF1 and 89 unaffected siblings, NF1 patients showed significantly elevated rates of difficulty initiating and maintaining sleep, arousal disturbances, and sleep-wake transition problems, deficits that persisted even after accounting for ADHD diagnosis, cognitive impairment, and stimulant medication use, indicating that fragmented, poor-quality sleep is a core feature of NF1 rather than a secondary consequence of its neurobehavioral comorbidities (Licis et al., 2013). Objective polysomnographic recordings confirm and extend this observation: relative to typically developing controls, NF1 children show a nearly four-fold increase in hourly arousals (7.0 vs. 1.9), reduced sleep efficiency (80% vs. 91%), and markedly shortened total sleep time (405 vs. 534 minutes), demonstrating that NF1 sleep is not merely subjectively poor but objectively fragmented (Carotenuto et al., 2023). This disruption is not confined to childhood: adults with NF1 report global sleep quality scores more than three-fold worse than population norms, with frequent nighttime awakenings and disrupted sleep continuity persisting across the lifespan (Leschziner, Golding et al. 2013). In adults, sleep disturbance is linked to pain, anxiety and depression (Leschziner, Golding et al. 2013). Regardless of trigger, fragmentation disrupts consolidated sleep, the continuous, typically nocturnal block thought ideal for physical and mental restoration (Bonnet and Arand 2003). Sleep fragmentation likely underlies the exhaustion and low energy patients report (Vassallo, Mughal et al. 2020). However, how Nf1 loss in the CNS affects sleep fragmentation regarding the neuronal populations and molecular mechanisms involved, is not well-explored. Moreover, although missense mutations comprise about a third of the known patient mutant alleles (Fahsold, Hoffmeyer et al. 2000, Zhang, Tong et al. 2015, Chen, Li et al. 2024),whether and how such variants affect all or particular attributes of sleep is unclear.

Although the function of sleep is not fully resolved, its neuronal and functional basis is remarkably conserved from mammals to invertebrates, including *Drosophila melanogaster* (Joiner 2016, Keene and Duboue 2018, Shafer and Keene 2021). Owing to its unmatched genetic tractability, *Drosophila* has proven invaluable for dissecting the genetic, cellular, and evolutionary basis of sleep, including its molecular mechanisms and behavioral consequences (Allada, Cirelli et al. 2017, Ly, Pack et al. 2018, Beckwith and French 2019, Artiushin and Sehgal 2020, Dissel 2020, Weiss and Donlea 2021). Flies show ∼24 hr circadian rhythmicity comprising cycles of locomotion, rest and sleep (Hendricks, Finn et al. 2000, Rosato and Kyriacou 2006, Donlea, Pimentel et al. 2018). Sleep in Drosophila is defined as behavioral quiescence lasting at least 5 minutes, occurring repeatedly across the subjective day and night (Hendricks, Finn et al. 2000, Joiner, Crocker et al. 2006), with bout frequency and duration peaking at night and around subjective midday (Hendricks, Finn et al. 2000, Joiner, Crocker et al. 2006). As in vertebrates, longer uninterrupted nighttime bouts are thought to reflect consolidated sleep, which is modulated by hunger, with starved flies showing shorter bouts (Hasegawa, Tomita et al. 2017). *Drosophila* sleep also parallels mammalian sleep in showing distinct CNS activity patterns, reduced metabolic rate, and decreased arousal threshold during quiescence (Shafer and Keene 2021), suggesting that flies display all core hallmarks of vertebrate sleep and possibly even distinct stages (Shafer and Keene 2021). Sleep is governed by multiple conserved, interacting genetic loci (Blum, Bell et al. 2018) and by distinct neuronal circuits that regulate sleep pressure, sleep and wakefulness (Shaw, Cirelli et al. 2000, Yurgel and Keene 2018, Shafer and Keene 2021), making *Drosophila* well suited to investigating the genetic basis of sleep deficits.

The *Drosophila* genome contains a single *Nf1* gene (*dNf1*), potentially encoding five nearly identical isoforms differing only at their carboxy-termini, with 68% similarity to the longest human isoform (Atsoniou, Giannopoulou et al. 2024). The fly Nf1 gene encodes a large GTPase-activating protein (GAP) that negatively regulates Ras signaling and appears to independently modulate cAMP/PKA signaling {Guo, 1997 #287;Guo, 2000 #277;The, 1997 #253;Tong, 2002 #292}. Despite its size, few *dNf1* mutations have been described, in contrast to the many missense mutations in the human gene that produce highly variable NF1 phenotypes with poor genotype-phenotype correlation (Koczkowska, Chen et al. 2018, Koczkowska, Callens et al. 2019, Koczkowska, Callens et al. 2020). Null *dNf1* mutants recapitulate many hallmarks of the human disorder, including reduced body size, impaired associative learning, altered circadian activity, behavioral inflexibility (e.g., excessive grooming), impaired social behavior, and aberrant sleep (Atsoniou, Giannopoulou et al. 2024) making the fly a valuable model to investigate molecular pathways and neuronal populations underlying each phenotype. Although several studies have examined sleep phenotypes in *dNf1* mutants (Bai and Sehgal 2015, Machado Almeida, Lago Solis et al. 2021, Brown, Zhang et al. 2023, Durkin, Poe et al. 2023), the specific neurons requiring dNf1—and the molecular mechanisms involved—for normal sleep remain undefined. Complicating matters, the commonly used null allele arises from a large deletion that also removes neighboring loci of the neurogenesis-regulating E(spl) complex (The, Hannigan et al. 1997, Delidakis, Monastirioti et al. 2014), potentially confounding neuronal development and contributing to the reported sleep deficits (Durkin, Poe et al. 2023).

To resolve these issues, we used existing and novel null and missense *dNf1*mutants to characterize their sleep patterns and potential fragmentation. In addition, we employed a spatially restricted RNAi knockdown strategy to identify and characterize neurons in the fly CNS requiring Neurofibromin for normal sleep. Furthermore, we investigate whether dNf1 disruption therein results in sleep fragmentation, whereas reintroduction of the protein in these neurons of mutants suffices to eliminate the sleep deficit and explore the requisite molecular mechanisms engaged in the process.

## RESULTS

### Reduced and fragmented sleep upon Nf1 loss

Loss of dNf1 in homozygous null *dNf1^E2^*mutants was first confirmed by quantitative proteomics of mutant versus control head lysates, which verified the absence of the protein (Fig 1A). To determine whether loss of dNf1 results in reduced or aberrant sleep, null mutants and controls were monitored for day-and night-time sleep using DAM5 monitors (Trikinetics). Unlike controls, *dNf1^E2^*homozygotes showed markedly reduced sleep with rudimentary periodicity (Fig 1B), amounting to a >60% reduction in total sleep over the assessed period (Fig 1C). Whereas controls showed the expected significant (p<0.0001, Suppl Table 1) increase in nighttime sleep (Hendricks, Finn et al. 2000, Shaw, Cirelli et al. 2000), null homozygotes did not (p=0.1267, Suppl Table 1) (Fig 1D). Correspondingly, the nighttime rise in bout number (Fig 1E) and bout duration (Fig 1F) seen in controls was largely absent in mutants, whose sleep bouts and duration remained low and stable across day and night. This blunted nighttime sleep consolidation was mirrored by elevated activity both during the night (Fig 1G) and between bouts (Fig 1H), in contrast to the marked activity reduction seen at night in controls.

**Figure 1.**
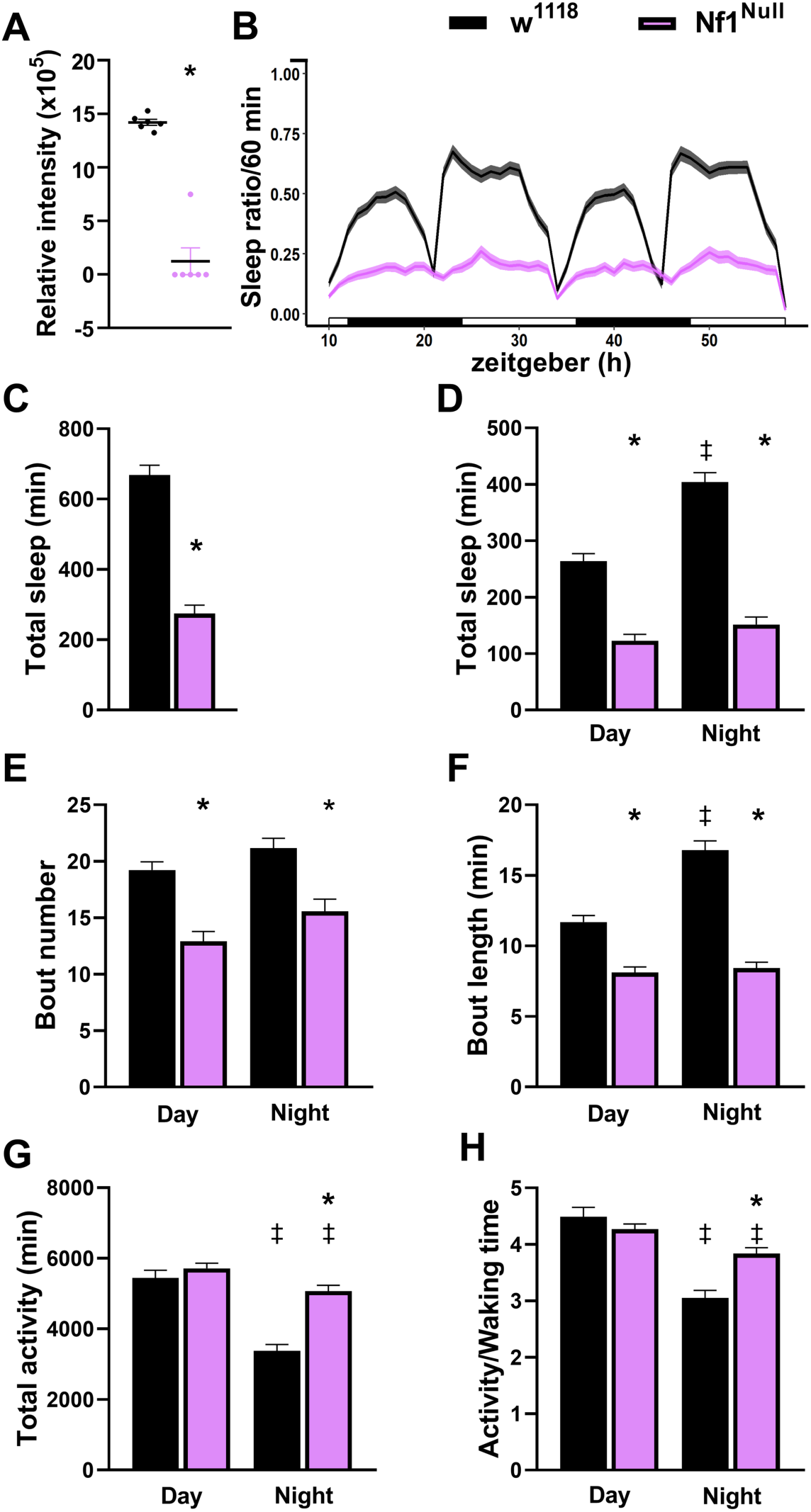
Sleep deficits and hyperactivity of null *dNf1* homozygotes. **A.** The abundance of Neurofibromin-derived tryptic peptides in head lysates of *w^1118^* control animals (black dots) and *dNf1^Null^* homozygotes (magenta dots) was calculated based on the normalized spectral intensity. The difference is highly significant (p<0.0022, n=6) and is indicated by the asterisk. Statistical details in Suppl. Table 1. **B.** An actogram quantifying the amount of time spent asleep per hour over a 48 hr period as indicated for *w^1118^* controls (black line, n=156) and *dNf1^Null^* homozygotes (magenta line, n=127). The darker lines indicate the mean, and the lighter width of the lines indicates the standard error of the mean (SEM). Daytime is denoted by the open bars and nighttime by the filled bars on the x axis. **C-H.** Quantification of sleep and its attributes in *w^1118^* control animals (black bars) and *dNf1^Null^* homozygotes (magenta bars). As all data were not normally distributed (as determined by Shapiro-Wilk, a<0.05), all statistical comparisons were performed with Mann-Whitney tests, with statistical details presented in Suppl. Table 1. Stars indicate that the mutant score is significantly different than the control, while the double dagger signifies differences in controls or mutants in the day relative to night-time scores. **C.** The total time spent asleep over the monitored period was significantly less (p<0.0001) for *dNf1^Null^* homozygotes than *w^1118^* controls, as indicated by the asterisk. **D.** The total time spent asleep during the day and at night revealed that while *w^1118^* control animals slept significantly more at night than during the day (p<0.0001, double dagger), *dNf1^Null^* homozygotes, slept significantly less than controls both during the day and at night (p<0.0001 for both) denoted by the asterisks, and their sleep at night was not increased relative to daytime (p=0.1267). **E.** The number of sleep episodes was significantly reduced in *dNf1^Null^* homozygotes compared to those of *w^1118^* controls both during the day and at night (p<0.0001 for both) denoted by the asterisks. **F.** Sleep bout length was increased at night for control animals (p<0.0001, double dagger), but not for the nulls (p=0.0724), which also presented with significantly reduced bout length (p<0.0001) relative to controls (asterisks). **G.** Total activity was decreased at night relative to daytime for controls (double dagger, p<0.0001) and for the nulls (double dagger, p=0.0053). However, nighttime activity remained significantly elevated in the nulls (p<0.0001) compared to controls (asterisk). **H.** Activity between sleep episodes decreased at night relative to daytime for controls and for the nulls (p<0.0001 for both) indicated by the double daggers. However, nighttime activity remained significantly elevated in the nulls (p<0.0001) compared to controls (asterisk), although their nighttime activity was reduced compared to that during the day (p=0.0014).

Taken together, the fewer, shorter sleep bouts and heightened inter-bout activity at night indicate shortened and fragmented sleep in *dNf1* null mutants, resembling sleep quality reported by NF1 patients (Marana Perez, Duat Rodriguez et al. 2015, Carotenuto, Messina et al. 2023). The same deficits appeared in the independently generated *dNf1^Δ216^*homozygotes and in heteroallelic combinations of the two null alleles, but not in *dNf1^E2^* heterozygotes, confirming the phenotype in the nulls is recessive (Suppl Fig 1). Thus, dNf1 loss both reduces and fragments sleep, consistent with impaired sleep consolidation.

Because sleep in *Drosophila* involves multiple neuronal circuits (Shafer and Keene 2021), mapping the adult CNS neurons expressing dNf1 could clarify its role in sleep regulation and the deficits arising from its loss. A novel CRIMIC reporter line (Kanca, Zirin et al. 2022) revealed the neuronal circuits with preferential dNf1 expression. dNf1 was found broadly distributed in the brain (Fig 2A), including the mushroom bodies, ellipsoid body, antennal lobes, and neurons near the GABAergic population previously implicated in the learning deficits of the *dNf1^E2^* null mutants (Georganta, Moressis et al. 2021).

**Figure 2.**
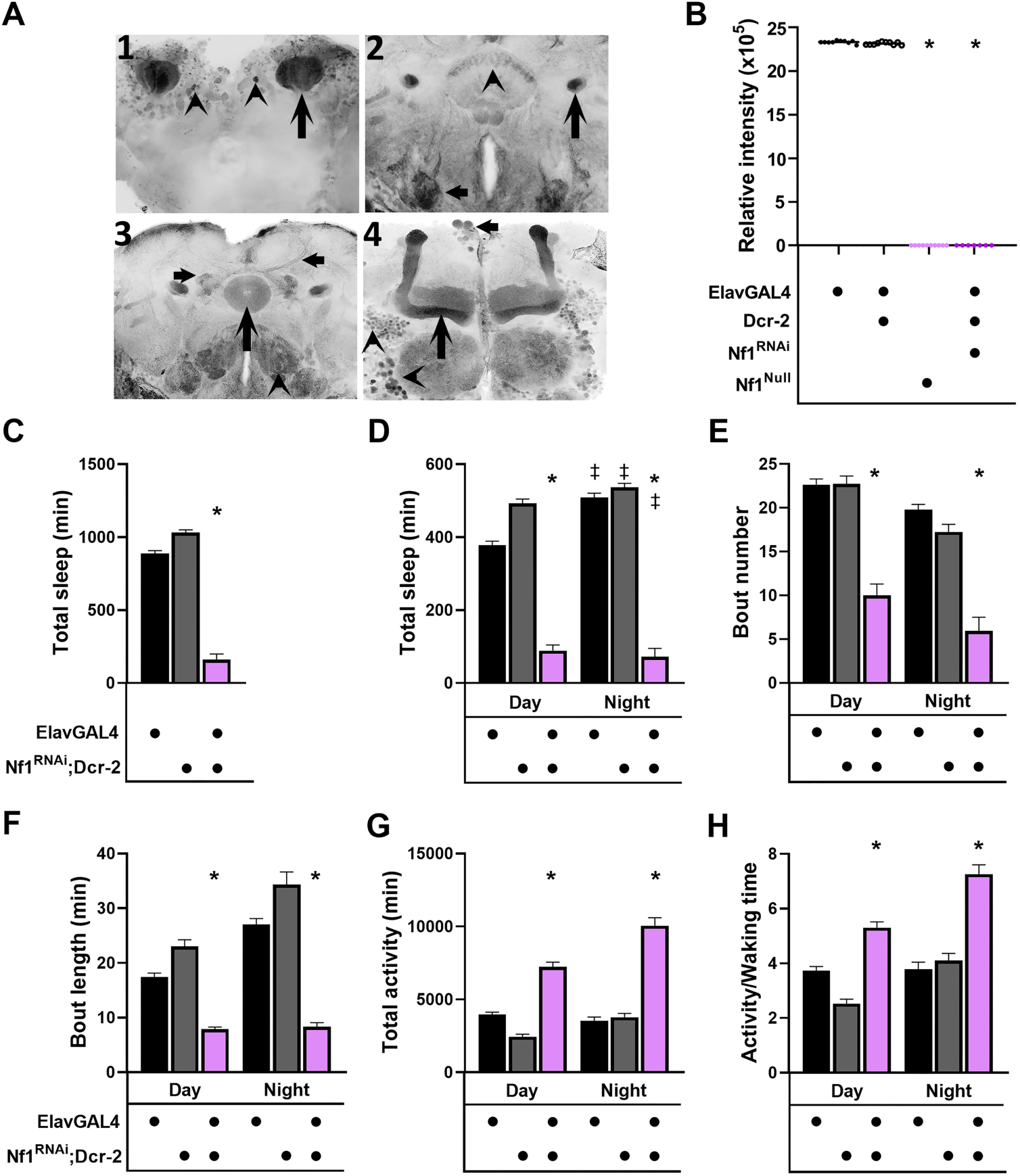
Pan-neuronal attenuation of dNf1 results in sleep deficits and hyperactivity. **A.** The distribution of dNf1 in the adult Drosophila brain revealed by the *dNf1^CRIMIC^* line driving GFP expression. The single-plane confocal images are arranged from posterior to anterior (1-4). **1**. Intense accumulation of dNf1 in the dendrites of the mushroom bodies (arrow) and scattered neurons in the dorsal posterior of the brain (arrowheads). **2**. dNf1 abounds in the axonal projections of the mushroom body neurons (arrow), neuronal projections innervating the dorsal fan-shaped body (arrowhead), neurons in the suboesophageal area (small arrow), among other neuronal tracts projecting throughout the brain. **3**. dNf1 accumulation in the ellipsoid body (arrow) and neuronal tracts in its vicinity (small arrows), possibly innervating it and neurons in the suboesophageal area (arrow heads). **4**. Robust accumulation of dNf1 in the αβ-type axonal projections and weaker in the α΄β΄of the mushroom body (arrow). dNf1 accumulates in the dorsally located insulin-like-peptide-producing neurons (small arrow) and multiple cell bodies ventrolateral to the mushroom body lobes (arrowheads) in the vicinity of the antennal lobes (star). **B.** The abundance of Neurofibromin-derived tryptic peptides in head lysates of ElavGal4/+ control animals, animals pan-neuronally expressing Dcr-2 and a dNf1 RNAi-encoding transgene along with Dcr-2, and *dNf1^Null^* homozygotes. Relative abundance was calculated based on the normalized spectral intensity. Although no significant difference was revealed upon pan-neuronal Dcr-2 expression, dNf1 abundance was highly reduced in all other genotypes (p<0.0001), as indicated by the asterisks. Statistical details in Suppl. Table 1. **C-H.** Quantification of sleep and its attributes in ElavGal4/+ control animals (black bars), flies expressing pan-neuronally Dcr-2 alone (gray bars) and those co-expressing a dNf1 RNAi-encoding transgene and Dcr-2 (magenta bars). Multiple non-parametric (as determined by Shapiro-Wilk, a<0.05) comparisons were performed with Kruskal-Wallis tests, while two-way comparisons were performed with Mann-Whitney. Multiple comparisons of normally distributed data were performed with Dunnett’s test following initial ANOVA. All statistical comparisons are detailed in Suppl. Table 1. Stars indicate that the mutant score is significantly different than that of the control animals, while the double dagger signifies differences in controls or mutants in the day relative to night-time scores.**C.** The total time spent asleep over the monitored period was significantly less (p<0.0001, asterisk) for ElavGal4>*dNf1^RNAi^*; Dcr-2-expressing animals, relative to that of ElavGal4/+ and *dNf1RNAi*; Dcr-2 control flies. **D.** The total time spent asleep during the day and at night revealed that ElavGal4/+ and *dNf1RNAi*; Dcr-2 control flies slept significantly more at night than during the day (p<0.0001, double daggers). In contrast, ElavGal4>*dNf1^RNAi^*; Dcr-2-expressing animals slept significantly less than controls both during the day and at night (p<0.0001 for both, asterisks), although their sleep at night (89.02 ± 15.65) appeared slightly increased relative to daytime (72.34 ± 22.74, p=0. 0003).

Given the broad dNf1 distribution in the adult CNS, we first sought to phenocopy the reduced, fragmented sleep of null mutants by abrogating dNf1 pan-neuronally via RNAi, driven by ElavGal4 in the presence of the RNAi enhancer, Dcr-2 (ElavGal4; UASDcr2). Quantitative proteomics showed that pan-neuronal dNf1 attenuation under ElavGal4; UASDcr2 reduced its levels in head lysates to those in null mutants (Fig 2B). Consequently, a robust reduction in total sleep emerged in these dNf1 knockdown flies (Fig 2C), which was distributed evenly across day and night (Fig 2D). The sleep deficit was driven at least in part, by fewer and significantly shorter sleep bouts relative to controls both during the day and night-time (Fig 2E, F). This sleep loss was mirrored by correspondingly large increases in day and night-time total activity (Fig 2G), or while awake between bouts (Fig 2H). Therefore, pan-neuronal dNf1 abrogation phenocopied the sleep fragmentation of null mutants (Fig 1)., though the effects on total sleep and hyperactivity appeared more severe than in the null mutants themselves (Fig 2 vs. Fig 1).

This raised the question of whether the enhanced sleep defects and excess activity might be driven by Dcr-2 expression itself. Pan-neuronal Dcr-2 expression alone resulted in a modest but significant reduction in total sleep (Suppl Fig 2A, B), ostensibly due to the surprising increase in day-and night-time activity (Suppl Fig 2E, F), rather than any significant change in bout number or duration (Suppl Fig 2C, D). Thus, Dcr-2 expression has a minor effect on total sleep but affects activity more substantially, which could partly explain the exaggerated phenotype of dNf1 knockdown under ElavG4; Dcr2 relative to *dNf1^E2^* homozygotes (Fig 1). This conclusion was verified regarding sleep by attenuating dNf1 pan-neuronally under Elav in the absence of Dcr2, which yielded substantially deficient day and night-time sleep (Suppl Fig 3A-C), though not as severe as under ElavG4; Dcr2. Therefore, with this caveat noted, Dcr-2 was retained in all subsequent experiments to optimize RNAi efficacy for tissue-specific emulation of the dNf1 null state.

Collectively, pan-neuronal dNf1 abrogation phenocopies the sleep consolidation deficit of *dNf1^E2^* homozygotes, consistent with independent reports of sleep deficits in other null and hypomorphic *Nf1* mutants (Bai and Sehgal 2015, Brown, Zhang et al. 2023, Durkin, Poe et al. 2023). Therefore, dNf1 activity is required to regulate both day-and night-time sleep by promoting sleep episodes of appropriate length.

### dNf1 loss from multiple circuits where it is preferentially expressed does not reduce sleep

Given that pan-neuronal dNf1 abrogation phenocopied the deficits of the null mutants, we aimed to define neurons where the protein is required to support physiological sleep. Because sleep is regulated by multiple neurons and circuits in the adult fly brain (Shaw, Cirelli et al. 2000, Blum, Bell et al. 2018, Yurgel and Keene 2018, Shafer and Keene 2021), we focused on circuits where dNf1 accumulates preferentially (Fig 2A).

A prominent such circuit is the mushroom body (MB). MB neurons are bilateral clusters of roughly 2500 Kenyon cells (KCs) in the dorsal posterior brain, with distinct tripartite axonal morphology (Crittenden, Skoulakis et al. 1998), and are essential for associative and non-associative learning and memory (Roussou, Papanikolopoulou et al. 2019, Foka, Georganta et al. 2022, Fiala and Kaun 2024). Sleep is reduced when KCs are silenced (Joiner, Crocker et al. 2006, Pitman, McGill et al. 2006, Bai and Sehgal 2015) and circadian fluctuation of dNf1 levels in these neurons contributes to daytime wakefulness (Machado Almeida, Lago Solis et al. 2021). This predicts that dNf1 loss in the MBs would promote, rather than impair, consolidated sleep by reducing daytime wake drive. We therefore tested this hypothesis by attenuating dNf1 in the MBs using the spatially restricted RNAi approach described above.

Knockdown of dNf1 specifically in the MBs (MB247-Gal4 (Aso, Grübel et al. 2009)) did not significantly affect total sleep, bout number, or bout length during the day or at night relative to driver controls (Fig 3A-C). Similar results were obtained with a different MB-specific driver, Leo-Gal4 (Messaritou, Leptourgidou et al. 2009) (Suppl Fig 3D-F). However, attenuating dNf1 in all CNS neurons except the MBs (Elav-Gal4;247-Gal80) precipitated significant deficits (Suppl Fig 3G), especially in night-time sleep (Suppl Fig 3H) and highly reduced sleep night-time episode number (Suppl Fig 3I) and their length during the day and at night (Suppl Fig 3). Collectively, these data suggest that despite its significant enrichment in the MBs (Fig 2A), neurons that have been implicated in sleep regulation (Wu and Liu 2023), dNf1 is not required therein to mediate this function.

**Figure 3.**
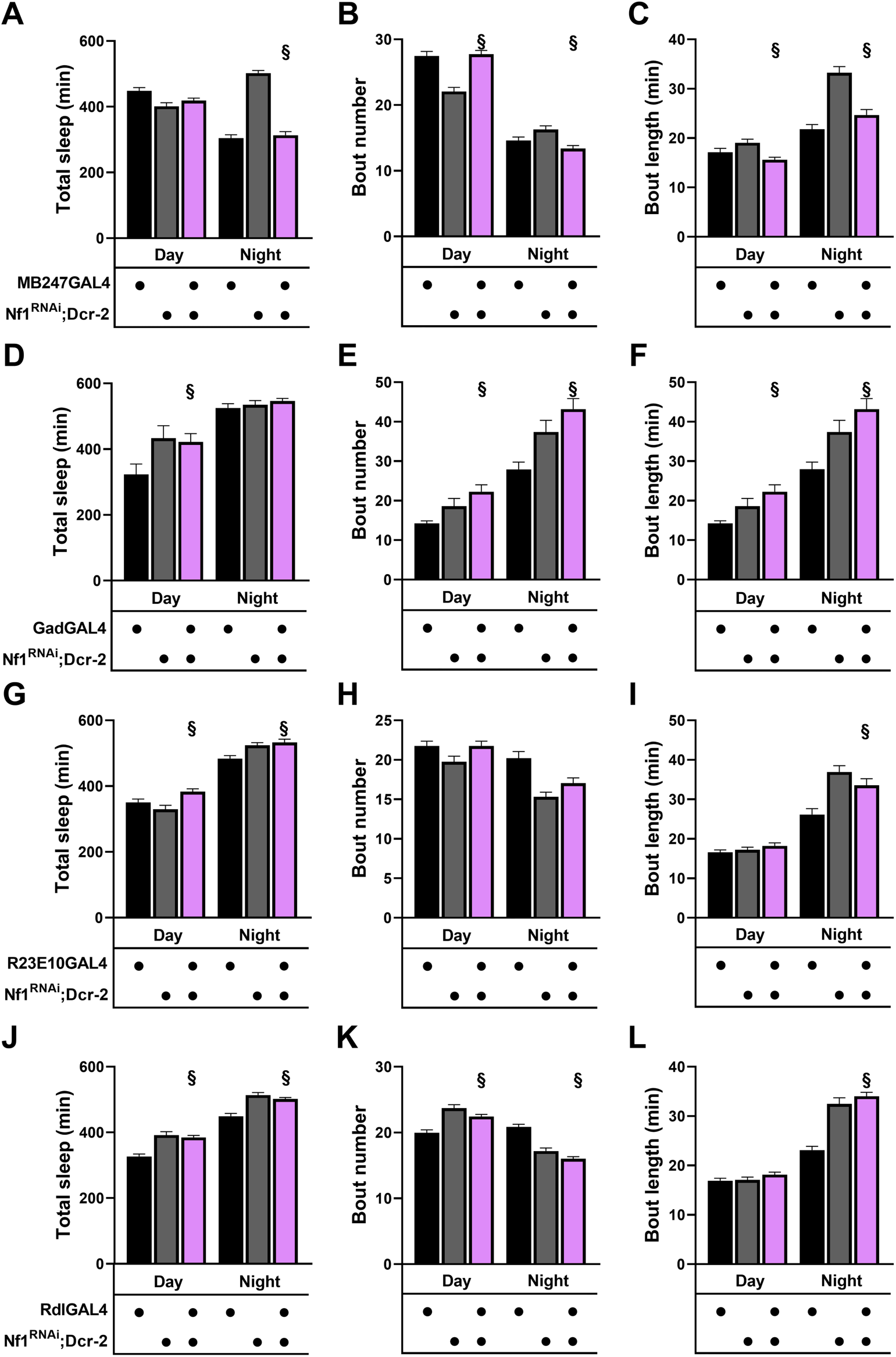
Abrogation of dNf1 in many neurons where the protein is preferentially distributed does not affect sleep significantly. Quantification of sleep and its attributes in animals where dNf1 is abrogated, as indicated below by the tissue-specific drivers (magenta bars) and the respective controls driver/+ (black bars) and *dNf1^RNAi^*/+;Dcr-2/+ (grey bars). Multiple non-parametric (as determined by Shapiro-Wilk, a<0.05) comparisons were performed with Kruskal-Wallis tests, while two-way comparisons were performed with Mann-Whitney. All statistical comparisons are detailed in Suppl. Table 1. The § symbol denotes significant differences of the experimental animal scores from one, but not both controls. **A, B, C.** Attenuation of dNf1 in MBns under MB247Gal4. **A**. No significant difference in daytime sleep was revealed, while nighttime sleep of MB247Gal4>*dNf1^RNAi^*; Dcr-2 was significantly different than that of *dNf1^RNAi^*/+; Dcr-2/+ (p<0.0001), but not from MB247Gal4/+ (p=0.9999). **B.** The number of sleep episodes of MB247Gal4>*dNf1^RNAi^*; Dcr-2 flies was significantly different only from one of the controls (*dNf1^RNAi^*/+; Dcr-2/+, p<0.0001, section sign) during the day and at nighttime (p= 0.0005, section sign). **C.** In accord with this, sleep bout length of MB247Gal4>*dNf1^RNAi^*; Dcr-2 flies was also significantly different only from *dNf1^RNAi^*/+; Dcr-2/+ flies both during the day and at nighttime (p=0.0011 and p<0.0001, respectively). **D, E, F.** Attenuation of dNf1 in GABAergic neurons under GadGal4. **D**. Daytime sleep of GadGal4>*dNf1^RNAi^*; Dcr-2 animals was significantly different only from GadGal4/+ flies (p=0.0056, section sign), but no significant difference was revealed at nighttime. **Ε**. Similarly, sleep bout number of GadGal4>*dNf1^RNAi^*; Dcr-2 flies was significantly different only from GadGal4/+ animals both during the day (p=0.0009, section sign) and at night (p<0.0001, section sign). **F**. Likewise, sleep bout length of GadGal4>*dNf1^RNAi^;* Dcr-2 flies was significantly different only from that of GadGal4/+ during the day (p=0.0009) and at night (p<0.0001). **G, H, I.** Attenuation of dNf1 in neurons of the dorsal Fan Shaped Body under R2310Gal4. **G**. Total daytime sleep of R2310Gal4>*dNf1^RNAi^*; Dcr-2 flies was increased relative to dNf1^RNAi^/+; Dcr-2/+ (p=0.0017, section sign), but at night it was different only from that of R2310Gal4/+ (p=0.0004 section sign). **H**. No significant differences in sleep bout number were revealed. **I**. Sleep bout length of GadGal4>*dNf1^RNAi^*; Dcr-2 flies was significantly different only from that of R2310Gal4/+ and during the nighttime (p=0.0025, section sign). **J, K, L.** Attenuation of dNf1 in neurons expressing the Rdl receptor under RdlGal4. **J**. Sleep of RdlGal4>*dNf1^RNAi^*; Dcr-2 flies was significantly different only from that of Rdl/+ animals both during the day (p<0.0001, section sign) and at night (p<0.0001). **K**. Sleep bout length of RdlGal4>*dNf1^RNAi^;* Dcr-2 flies was significantly different from that of Rdl/+ during the day and night (p<0.0001, section sign). **L**. Sleep bout length of RdlGal4>*dNf1^RNAi^*; Dcr-2 animals was significantly different only from that of Rdl/+ controls during the night (p<0.0001).

Neurofibromin is required within at least some GABAergic neurons, as its loss there elevates GABA near the MBs and inhibits associative learning (Georganta, Moressis et al. 2021). GABA also inhibits circadian arousal neurons and modulates sleep homeostasis, such that excess GABA increases sleep (Chaturvedi and Emery 2024). If dNf1 is required in GABAergic neurons to restrain the levels or release of this neurotransmitter as is for associative learning (Georganta, Moressis et al. 2021), then its loss therein should elevate sleep rather than impair it. Therefore, we investigated whether dNf1 is required in GABAergic neurons to promote or suppress sleep and its consolidation. However, pan-GABAergic dNf1 abrogation (under Gad-Gal4) did not compromise total sleep. Instead, it increased daytime sleep (Fig 3D) and increased the number and length of sleep episodes, mostly at night (Fig 3E, F), though these effects were significant versus driver controls for bout number and length, but not versus dNf1^RNAi^;Dcr2 heterozygotes (p=0.056 Fig 3D; p=0.0009 Fig 3E day and p<0.0001 at night). Thus, dNf1 loss from GABAergic neurons does not result in reduced and fragmented sleep. Therefore, the fragmented sleep of the null mutants does not result from dNf1 loss in GABAergic neurons.

Neurons of the dorsal fan-shaped body (dFSB) are also implicated in sleep regulation (Donlea, Pimentel et al. 2014, Donlea, Pimentel et al. 2018). As dNf1 appears to be distributed in this area (Fig 2A2), we investigated whether the protein is required therein to regulate sleep. Thus, we abrogated dNf1 in neurons marked with the R23E10Gal4 driver (Donlea, Pimentel et al. 2014), because of its similar expression in the dFSB with that of dNf1 and the role of these neurons in promoting sleep (Donlea, Pimentel et al. 2014, Donlea, Pimentel et al. 2018). However, abrogation of the protein in these neurons did not reduce but rather modestly increased (p=0.0017) daytime and a significantly increased (p=0.0004) night-time sleep, relative to driver heterozygotes, but not relative to Nf1^RNAi^; Dcr2 heterozygotes (Fig 3G). This increase appeared to be largely due to the modest increase (p=0.0025) in bout length compared to driver heterozygotes (Fig 3I), which is likely due to Dcr2 expression (Suppl Fig 2D). Therefore, loss of dNf1 from these dFSB neurons, does not appear to contribute to the sleep deficits in *dNf1* null animals.

Furthermore, because of its expression in the ellipsoid body (Fig2A3), we also knocked down dNf1 in these neurons, which also resulted in significant (p<0.0001) increase in day and night-time sleep (Fig 3J), largely due to significantly increased (p<0.0001) bout length (Fig 3L), but again only relative to driver controls, also most likely due to Dcr2 expression (Suppl Fig 2D). Therefore, a conservative interpretation of these results is that dNf1 activity is also not necessary in the ellipsoid body to regulate sleep.

### dNf1 is required in cholinergic neurons to regulate sleep consolidation

Given the lack of significant sleep deficits and fragmentation following dNf1 loss from the neuronal assemblies assayed above and Neurofibromin’s broad distribution, we examined the effects of its elimination from the most abundant neuronal type in the adult CNS, the cholinergic neurons (Eckstein, Bates et al. 2024). This resulted in significantly reduced total sleep (Fig 4A), with the deficit parsed evenly between day and night-time sleep (Fig 4B). Daytime sleep-episode number did not differ from controls, but night-time episodes increased (p<0.0001), at least relative to driver heterozygotes (Fig 4C). Despite this rise in night-time episode number, episode length was sharply shortened (p<0.0001) relative to both controls, especially at night (Fig 4D). These results demonstrate sleep fragmentation similar to that in dNf1 nulls (Fig 1, Suppl Fig 1). However, unlike in null homozygotes, dNf1 loss specifically from cholinergic neurons did not elevate activity, as seen in mutants and with pan-neuronal dNf1 reduction (Fig 1G, H; Fig 2G, H). Instead, daytime activity declined relative to driver controls (p<0.0001 Fig 4E; p<0.0001 Fig 4F). Together, these results demonstrate that dNf1 loss in cholinergic neurons recapitulates the deficient, fragmented sleep of null mutants without significantly affecting activity. Therefore, in conjunction with the outcomes of pan-neuronal attenuation (Fig 2G, H), the apparent dNf1 contribution to locomotor activity regulation, does not involve cholinergic neurons.

**Figure 4.**
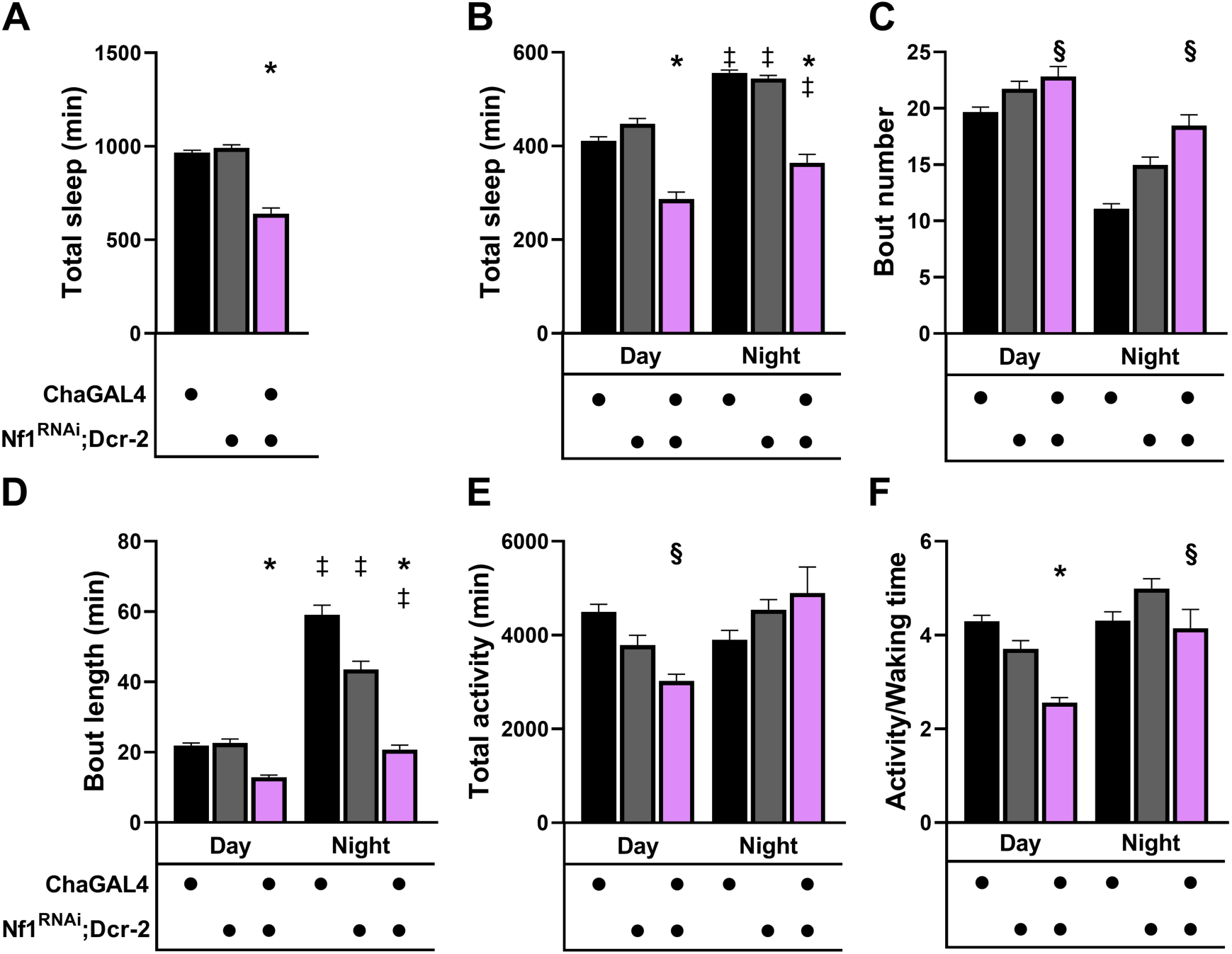
Loss of dNf1 from cholinergic neurons precipitates sleep deficits. Quantification of sleep and its attributes in cholinergic neurons under ChaGal4 driving dNf1^RNAi^; Dcr-2 (magenta bars), relative to the ChaGal4/+ (black bars) and dNf1^RNAi^/+; Dcr-2/+ (gray bars) controls. Multiple non-parametric (as determined by Shapiro-Wilk, a<0.05) comparisons were performed with Kruskal-Wallis tests, while two-way comparisons were performed with Mann-Whitney. Multiple comparisons of normally distributed data were performed with Dunnett’s test following initial ANOVA. All statistical comparisons are detailed in Suppl. Table 1. Stars indicate that the mutant score is significantly different than that of the control animals, the double dagger signifies differences in controls or mutants in the day relative to night-time scores, and § denotes significant differences from one, but not both controls. **A.** The total time spent asleep for ChaGal4>dNf1^RNAi^; Dcr-2 flies was significantly different than that of both controls (ANOVA, p<0.0001, Dunnett’s p<0.0001 for both). **B.** The total time spent asleep during the day and at night revealed that for both periods, ChaGal4>dNf1^RNAi^; Dcr-2 animals slept significantly less than both controls (p<0.0001 from both controls during the day and at night). Both controls slept more during the night (p<0.0001 for both, double daggers), and a similar increase was observed in the ChaGal4>dNf1^RNAi^; Dcr-2 flies (p=0.0007, double dagger). **C.** The number of daytime sleep episodes of ChaGal4>dNf1^RNAi^; Dcr-2 flies was significantly increased relative to those of the ChaGal4/+ controls (p=0.0044, section sign), but not from those of dNf1^RNAi^/+; Dcr-2/+ (p=0.8990). Similarly, at nighttime, sleep bouts of ChaGal4>dNf1^RNAi^; Dcr-2 animals were significantly elevated relative to those of ChaGal4/+ controls (p=0.0001, section sign), but not relative to those of dNf1^RNAi^/+; Dcr-2/+ (p=0.0170). **D.** Daytime sleep bout length analysis revealed the apparent cause of reduced sleep in ChaGal4>dNf1^RNAi^; Dcr-2 flies, as the time spent asleep per episode was significantly reduced relative to that of both controls both during the day (p<0.0001 from both, asterisk) and at night (p<0.0001 from both, asterisk). All three strains, however, slept longer during the night than during the day (p<0.0001 for all comparisons, double dagger). **E.** Total activity during the day was depressed for ChaGal4>dNf1^RNAi^; Dcr-2 flies, but only relative to that of ChaGal4/+ controls (p<0.0001), not from dNf1^RNAi^/+; Dcr-2/+ (p=0.0158). No significant differences were observed at night. **F.** Activity when awake was significantly reduced for ChaGal4>dNf1^RNAi^; Dcr-2 animals relative to both controls (p<0.0001 from both, asterisk), but only from dNf1^RNAi^/+; Dcr-2/+ at nighttime (p=0.0002).

Is Neurofibromin necessary and sufficient in cholinergic neurons for normal sleep? To address this question, we expressed a wild-type dNf1 transgene in cholinergic neurons of null mutants, aiming to reverse their deficient, fragmented sleep. Unlike null mutants, transgene-expressing animals showed largely normal sleep patterns (Fig 5A). Quantitation confirmed that dNf1 expression restored day and nighttime sleep in nulls, though not to control levels (Fig 5B, C). Pan-neuronal transgene expression in null homozygotes yielded similar results (Suppl Fig 4A-C), suggesting that the modest difference from control levels reflects inadequate transgene expression rather than a requirement for dNf1 in other than cholinergic neurons. Sleep periodicity and episode number during day and night were also largely restored, as was episode duration (Fig 5D, E; Suppl Fig 4D, E), consistent with the actograms (Fig 5A, Suppl Fig 4A). Since cholinergic dNf1 loss did not affect activity, we did not assess this parameter here or in subsequent experiments. Taken together, these results show that dNf1 activity in cholinergic neurons is necessary and sufficient to mediate consolidated sleep during both day and night.

**Figure 5.**
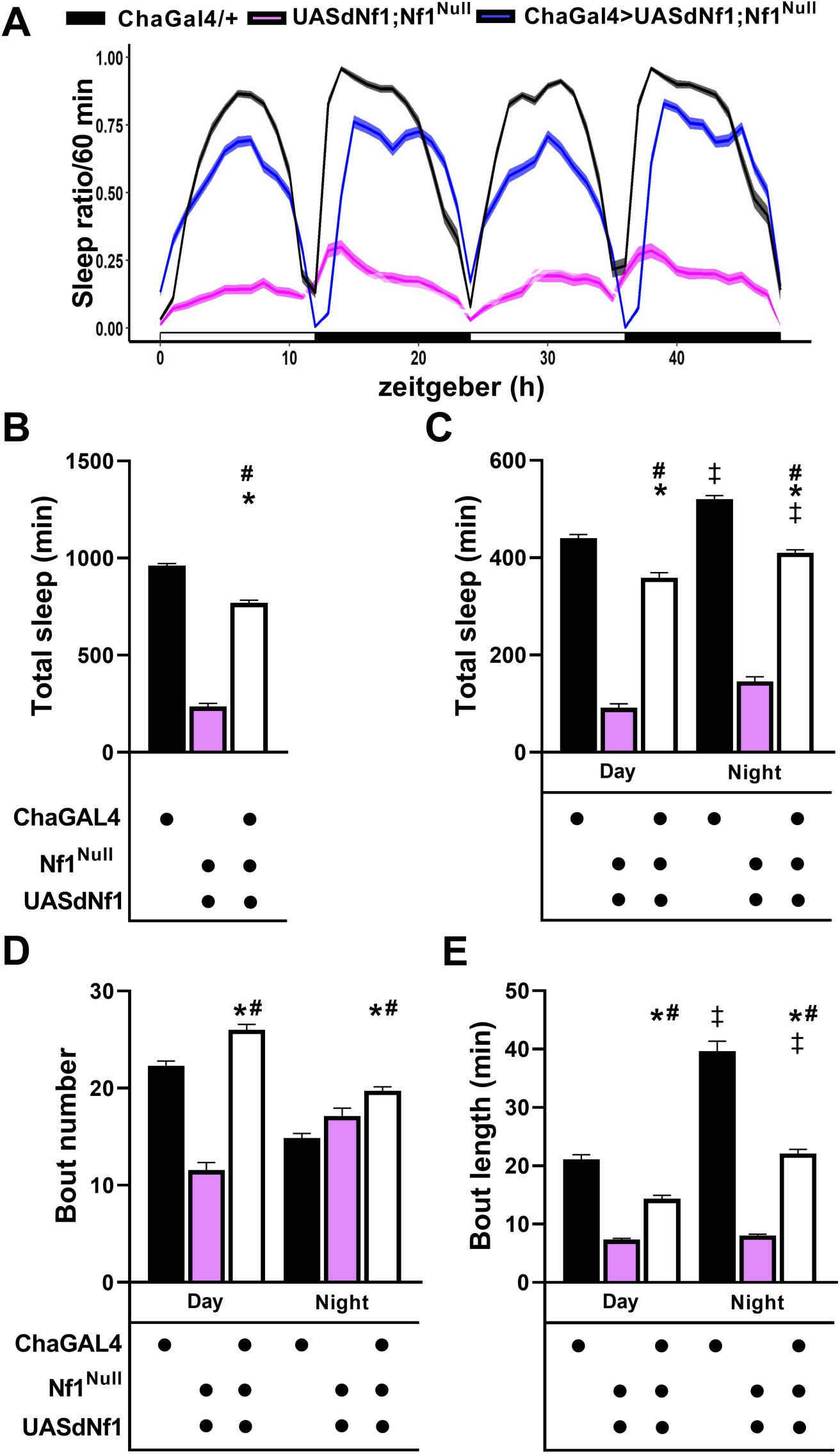
Transgenic restoration of dNf1 expression in cholinergic neurons ameliorates the sleep deficit of dNf1^Null^ homozygotes. **B.** An actogram quantifying the amount of time spent asleep per hour over a 48 hr period as indicated for ChaGal4/+ controls (black line) and UASdNf1/+; *dNf1^Null^* homozygotes (magenta line) and ChaGal4>UASdNf1/+; *dNf1^Null^* (blue line). The mean is indicated by the darker lines, and the standard error of the mean (SEM) is indicated by the lighter width of the lines. Daytime is denoted by the open bars and nighttime by the filled bars on the x axis.**B-E.** Quantification of sleep and its attributes in ChaGal4/+ controls (black bars) and UASdNf1/+; *dNf1^Null^* homozygotes (magenta bars) and ChaGal4>UASdNf1/+; *dNf1^Null^* animals (open bars). As not all data were normally distributed (as determined by Shapiro-Wilk, a<0.05), statistical comparisons were performed with Mann-Whitney tests, with statistical details presented in Suppl. Table 1. Stars indicate that the mutant score is significantly different than that of the experimental animals, the double dagger signifies differences in controls or mutants in the day relative to night-time scores, and # denotes significant differences of the experimental score compared to that of the controls. **B.** The total time spent asleep for ChaGal4>UASdNf1/+; *dNf1^Null^* flies was significantly different than that of UASdNf1/+; *dNf1^Null^* (p<0.0001, asterisk), but also from ChaGal4/+ controls (p<0.0001, hashtag). **C.** Daytime sleep of ChaGal4>UASdNf1/+; *dNf1^Null^* flies was significantly different from that of UASdNf1/+; *dNf1^Null^* (p<0.0001, asterisk), but also from that of ChaGal4/+ controls (p<0.0001, hashtag). The total time spent asleep during the day and at night revealed that ElavGal4/+ control flies and ElavGal4>*dNf1RNAi*; Dcr-2 flies slept significantly more at night than during the day (p<0.0001, double daggers). **D.** Daytime sleep bout number of ChaGal4>UASdNf1/+; *dNf1^Null^* animals was significantly different from that of UASdNf1/+; *dNf1^Null^* (p<0.0001, asterisk), but also from ChaGal4/+ controls (p<0.0001, hashtag). The number of sleep episodes of ChaGal4>UASdNf1/+; *dNf1^Null^*animals during the night was again significantly different from that of UASdNf1/+; *dNf1^Null^* (p<0.0026, asterisk), but also from that of ChaGal4/+ (p<0.0001, hashtag). **E.** Daytime sleep bout length of ChaGal4>UASdNf1/+; *dNf1^Null^* flies was significantly different from that of UASdNf1/+; *dNf1^Null^* (p<0.0001, asterisk), but also from ChaGal4/+ controls (p<0.0001, hashtag). Similarly, at night, sleep lasted longer for ChaGal4>UASdNf1/+; *dNf1^Null^* flies than that of UASdNf1/+; *dNf1^Null^* (p<0.0001, asterisk), but also than that of ChaGal4/+ (p<0.0001, hashtag). Moreover, as for ChaGal4/+ (p<0.0001, double dagger), sleep bout length was also significantly longer during the night (p<0.0001, double dagger) for ChaGal4>UASdNf1/+; *dNf1^Null^*animals.

### The GRD domain of Neurofibromin suffices to regulate sleep in cholinergic neurons

Loss of dNf1 is thought to result in unregulated Ras activity, which underlies multiple deficits in models (Atsoniou, Giannopoulou et al. 2024) and patients (Siqueiros-Sanchez, Serur et al. 2025). Neurofibromin regulates Ras via its GTPase-activating protein-related domain (GRD), comprising roughly 15% of the protein (Atsoniou, Giannopoulou et al. 2024) and this negative regulation of Ras-MAPK signaling represents its best-understood function to date. Since dNf1 expression in cholinergic neurons is necessary and sufficient to mediate sleep consolidation, we asked whether restoring Ras regulation alone could reverse the sleep deficits of *dNf1^E2^* homozygotes. To that end we used two transgenes, one expressing the wild-type GRD (*dNf1^GRD^*, amino acids 1214-1574), and another carrying an Arginine1320-to-Proline mutation (*dNf1^GRD^*^-*RP*^) shown to abolish its activity (Walker, Tchoudakova et al. 2006). The analogous mutation in the human gene likewise produces negligible GTPase activity (Klose, Ahmadian et al. 1998).

Initially we determined that expression of the wild type GRD domain in the cholinergic system restored the elevated phosphorylated MAPK of *dNf1^E2^* homozygotes to control levels (Fig 6A). We determined via proteomics that MAPK levels remain relatively constant in controls, the null and the other mutants used herein (Suppl Fig 5). Therefore, we use syntaxin levels to normalize the blots since the anti-MAPK antibody is marginally effective in our hands (not shown).

**Figure 6.**
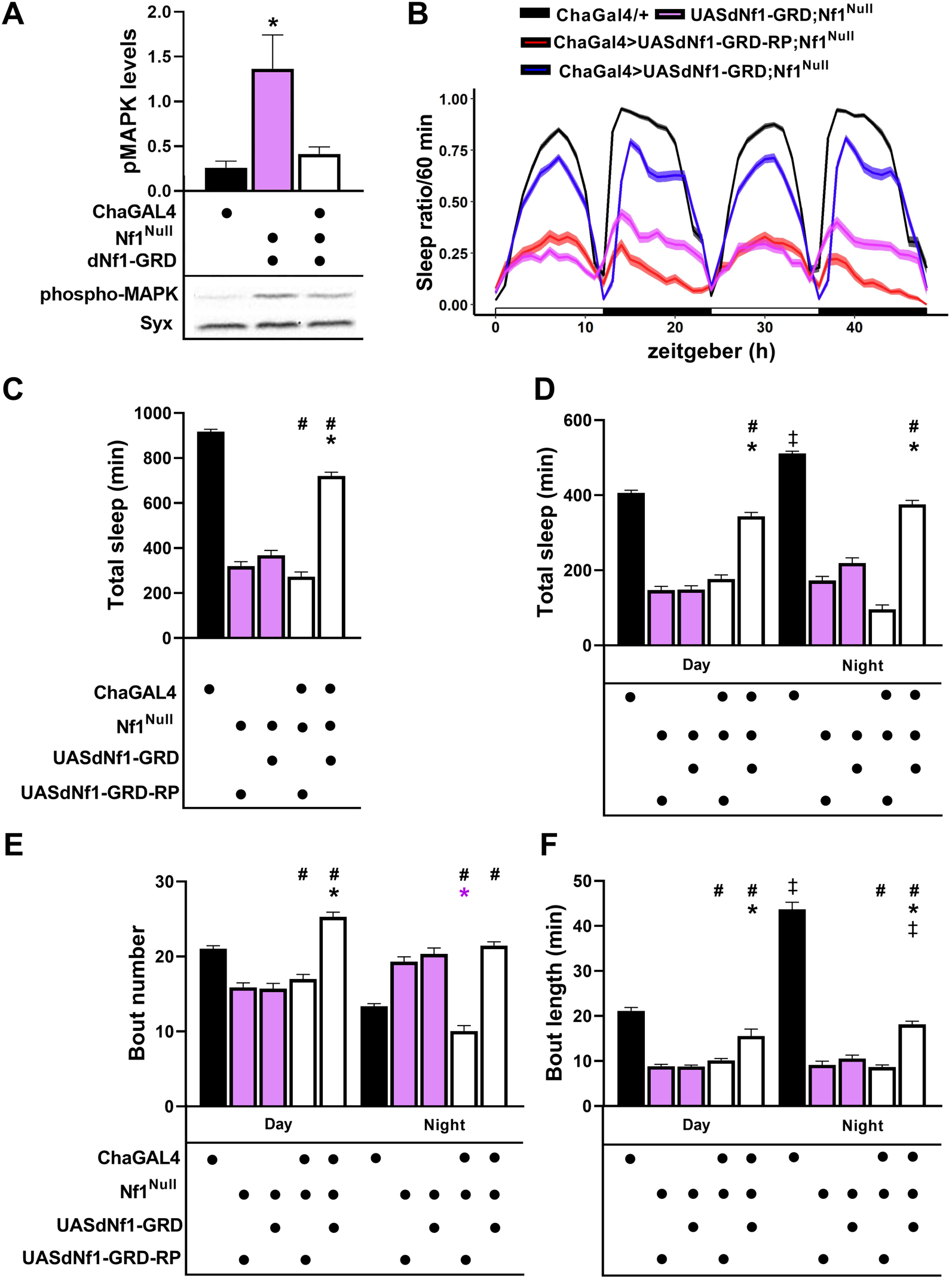
Transgenic expression of the dNf1 GRD in cholinergic neurons ameliorates the sleep deficit of dNf1^null^ homozygotes. **A.** Western blot analysis quantified below the blot demonstrated that phospho-MAPK levels in ChaGal4>UASdNf1-GRD; *dNf1^Null^* animals are not significantly different than those of ChaGal4/+ (p=0.3317) but were significantly elevated in UASdNf1-GRD/+; *dNf1^Null^* homozygotes (p=0.0043). Statistical details in Supplemental Table 1. **B.** An actogram quantifying sleep per hour over a 48 hr period as indicated for ChaGal4/+ controls (black line), UASdNf1/+; *dNf1^Null^* homozygotes (magenta line), and ChaGal4>UASdNf1GRD/+; *dNf1^Null^* (blue line) and ChaGal4>UASdNf1GRD-RP/+; *dNf1^Null^* (red line). The mean is indicated by the darker lines and the standard error of the mean (SEM) is indicated by the lighter width of the lines. Daytime is denoted by the open bars and nighttime by the filled bars on the x axis. **C-F.** Quantification of sleep and its attributes in ChaGal4/+ controls (black bars), UASdNf1GRD/+; *dNf1^Null^* homozygotes (magenta bars), UASdNf1GRD-RP/+; *dNf1^Null^* homozygotes (magenta bars), ChaGal4>UASdNf1GRD/+; *dNf1^Null^* animals and ChaGal4>UASdNf1GRD-RP/+; *dNf1^Null^* (open bars). As not all data were normally distributed (as determined by Shapiro-Wilk, a<0.05), statistical comparisons were performed with Mann-Whitney tests. Hashtags represent significant differences from ChaGal4/+ controls, and asterisks represent significant differences from the undriven UASdNf1-GRD/+; *dNf1^Null^*and UASdNf1-GRD-RP/+; *dNf1^Null^* mutants, with statistical details presented in Suppl. Table 1. Stars indicate that the mutant score is significantly different than that of the experimental animals, the double dagger signifies differences in controls or mutants in the day relative to night-time scores and # denotes significant differences of the experimental score compared to that of the controls. **C.** The total time spent asleep for ChaGal4/+ was significantly different than that for UASdNf1-GRD/+; *dNf1^Null^* (p<0.0001), but also from UASdNf1-GRD-RP/+;*dNf1^Null^* (p<0.0001) and Cha>UASdNf1-GRD-RP/+;*dNf1^Null^* (p<0.0001, hashtag) and significantly from Cha>UASdNf1-GRD/+;*dNf1^Null^* (p<0.0001, hashtag), indicating that reversal was not complete. However, sleep in Cha>UASdNf1-GRD/+;*dNf1^Null^* was significantly increased compared to UASdNf1-GRD/+; *dNf1^Null^* (p<0.0001, asterisk), demonstrating a partial rescue. In contrast, sleep of ChaGal4>UASdNf1-GRD-RP;Nf1^Null^ was not significantly different from that of UASdNf1-GRD-RP;Nf1^Null^ (p>0.9999), showing that the RP mutation abolishes rescue activity. **D.** Daytime sleep of ChaGal4/+ was significantly different than that of UASdNf1-GRD/+;*dNf1^Null^*(p<0.0001), UASdNf1-GRD-RP/+;*dNf1^Null^* (p<0.0001), Cha>UASdNf1-GRD-RP/+;*dNf1^Null^* (p<0.0001)and significantly from Cha>UASdNf1-GRD/+;*dNf1^Null^* (p=0.0076, hashtag), but significantly, daytime sleep of Cha>UASdNf1-GRD/+;*dNf1^Null^* was also significantly different than that of UASdNf1-GRD/+;*dNf1^Null^* (p<0.0001, asterisk). Night-time sleep was significantly elevated compared to that in the day for ChaGal4/+ (p<0.0001 double dagger) and was significantly different from that of UASdNf1-GRD/+;*dNf1^Null^* (p<0.0001), UASdNf1-GRD-RP/+;*dNf1^Null^* (p<0.0001), Cha>UASdNf1-GRD-RP/+;*dNf1^Null^* (p<0.0001) and from that of Cha>UASdNf1-GRD/+;*dNf1^Null^* (p<0.0001, hashtag). However, night-time sleep of Cha>UASdNf1-GRD/+;*dNf1^Null^* was also different from that of UASdNf1-GRD/+;*dNf1^Null^* (p<0.0001, asterisk). In contrast, sleep of Cha>UASdNf1-GRD-RP/+;*dNf1^Null^*was not significantly different from that of UASdNf1-GRD-RP/+;*dNf1^Null^* (p=0.0162). **E.** Daytime sleep bout number of ChaGal4/+ was significantly different than that of UASdNf1-GRD/+;*dNf1^Null^* (p<0.0001), UASdNf1-GRD-RP/+;*dNf1^Null^* (p<0.0001), ChaGal4>UASdNf1-GRD-RP/+;*dNf1^Null^*(p=0.001, hashtag) and Cha>UASdNf1-GRD/+;*dNf1^Null^* (p<0.0001 hashtag). Importantly, sleep bout number of ChaGal4>UASdNf1-GRD/+;*dNf1^Null^*was significantly different than that of UASdNf1-GRD-RP/+;*dNf1^Null^* (p<0.0001, asterisk) while that of ChaGal4>UASdNf1-GRD-RP/+;*dNf1^Null^*was not compared to UASdNf1-GRD-RP/+;*dNf1^Null^* (p=0.9999). Nighttime sleep episodes of ChaGal4/+ were significantly different than those of UASdNf1-GRD/+;*dNf1^Null^* (p<0.0001), UASdNf1-GRD-RP/+;*dNf1^Null^*(p<0.0001) and ChaGal4>UASdNf1-GRD/+;*dNf1^Null^* (p<0.0001, hashtag), but interestingly not from that of ChaGal4>UASdNf1-GRD-RP/+;*dNf1^Null^*(p=0.0433). Whereas ChaGal4>UASdNf1-GRD/+; *dNf1^Null^* did not perform significantly different than UASdNf1-GRD /+; *dNf1^Null^*, ChaGal4>UASdNf1-GRD-RP/+; *dNf1^Null^* was significantly different than that of UASdNf1-GRD-RP/+; *dNf1^Null^* (magenta asterisk). **F.** Daytime sleep bout length of ChaGal4/+ was significantly different than that of UASdNf1-GRD/+; *dNf1^Null^* (p<0.0001), UASdNf1-GRD-RP/+; *dNf1^Null^* (p<0.0001), ChaGal4>UASdNf1-GRD-RP/+; *dNf1^Null^*(p=0.001, hashtag) and importantly also from that of Cha>UASdNf1-GRD/+; *dNf1^Null^* (p<0.0001, hashtag). However, sleep episode length of Cha>UASdNf1-GRD/+; *dNf1^Null^* was also different from that of UASdNf1-GRD/+;*dNf1^Null^* (p<0.0001, asterisk), but was not for Cha>UASdNf1-GRD-RP/+; *dNf1^Null^* compared to UASdNf1-GRD-RP/+; *dNf1^Null^* (p=0.1258). Night-time sleep was significantly elevated compared to that in the day for ChaGal4/+ (p<0.0001 double dagger) and was also increased during the night for Cha>UASdNf1-GRD/+;*dNf1^Null^* (p<0.0001, double dagger). Bout length in ChaGal4/+ animals was significantly different than those of UASdNf1-GRD/+;*dNf1^Null^* (p<0.0001), UASdNf1-GRD-RP/+;*dNf1^Null^*(p<0.0001), ChaGal4>UASdNf1-GRD-RP/+;*dNf1^Null^* (p<0.0001, hashtag) and ChaGal4>UASdNf1-GRD/+;*dNf1^Null^* (p<0.0001, hashtag). However, bout length of ChaGal4>UASdNf1-GRD/+;*dNf1^Null^* was significantly different than that of UASdNf1-GRD/+;*dNf1^Null^* (p<0.0001, asterisk) while that of ChaGal4>UASdNf1-GRD-RP/+;*dNf1^Null^* was not compared to UASdNf1-GRD-RP/+;*dNf1^Null^*(p=0.9999).

Significantly, pan-cholinergic *dNf1^GRD^*expression also restored the day and night-time sleep patterns (Fig 6B) and levels (Fig 6C), as effectively as the full-length transgene (Fig 5B, C). Day and night-time sleep were equally restored (Fig 6D), as also were the respective number of sleep bouts (Fig 6E). Sleep episode length was also significantly improved (Fig 6F). Why control ChaGal4 heterozygotes present with excessively lengthy nighttime sleep bouts is unclear (Fig 6F) but was consistent in independent experiments (Fig 5E). Moreover, pan-neuronal expression of the *dNf1^GRD^* transgene yielded similar results, ameliorating the day and night-time sleep deficit (Suppl Fig 6A, B), normalizing of the number of sleep episodes (Suppl Fig 6C) and their duration (Suppl Fig 6D). In contrast, expression of the mutant *dNf1^GRD^*^-*RP*^ transgene did not ameliorate or alter the sleep deficits of null homozygotes (Fig 6B-D). Collectively the results demonstrate that the GTPase activity of Neurofibromin within cholinergic neurons is necessary and sufficient for normal sleep.

### Novel dNf1 missense mutants present with deficient and fragmented sleep

Modeling the effects of a loss of function allele is highly informative, but the number of patient-borne missense mutations in the *NF1* gene is significant and associated with pathological manifestations, (Fahsold, Hoffmeyer et al. 2000, Zhang, Tong et al. 2015, Koczkowska, Chen et al. 2018, Koczkowska, Callens et al. 2019, Koczkowska, Callens et al. 2020, Chen, Li et al. 2024). Despite the poor genotype/phenotype correlation, many missense mutation-bearing patients present with deficient and fragmented sleep (Marana Perez, Duat Rodriguez et al. 2015, Vassallo, Mughal et al. 2020, Carotenuto, Messina et al. 2023). Therefore, we wondered whether missense mutations in the Drosophila gene analogous to those in patients, would also present with deficient sleep. We focused on two missense mutations. First the *dNf1^C1045Y^*(Fig 7A), previously known as *dNf1^E4^*(Walker, Tchoudakova et al. 2006), in the region proximal to the Neurofibromin Tubulin Binding Domain, upstream of the GRD (Atsoniou, Giannopoulou et al. 2024), a domain of apparent functional relevance (Fahsold, Hoffmeyer et al. 2000) in the proximity of mutations (M1035R), associated with LEOPARD syndrome in humans (Wu, Legius et al. 1996). The second is a CrispR-generated mutation R1861C (*dNf1^R1861C^*) in the Drosophila gene, designed to mimic the R1809C missense mutation in patients (Pinna, Lanari et al. 2015, Rojnueangnit, Xie et al. 2015). This relatively common mutation affects the Plekstrin-Homology (PH) domain (Fig 7A) and is associated largely with cognitive deficits in humans.

**Figure 7:**
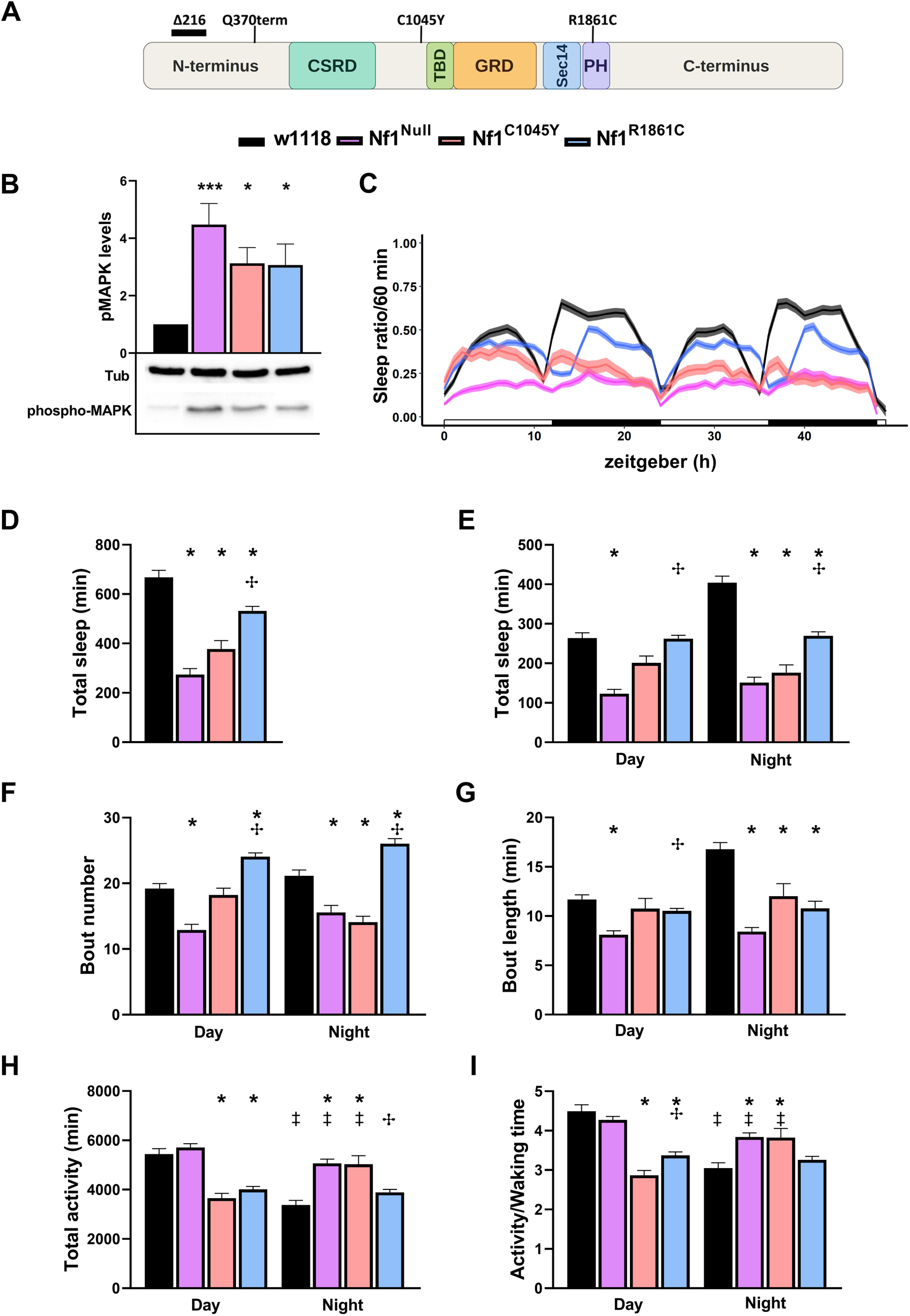
Characterization of pMAPK regulation, sleep profiles, and locomotor activity in dNf1 missense mutants. **A.** Schematic representation of the Drosophila Neurofibromin (dNf1) protein structure highlighting its functional domains: N-terminus, CSRD, TBD, GRD, Sec14, PH, and C-terminus. The locations of dNf1 Null Δ216, Q370term and point mutations C1045Y, and R1861C are marked above. Color-coded legend indicates the genotypes analyzed across panels: *w^1118^* control (black), *dNf1^Null^* (magenta), *dNf1^C1045Y^* (pink), and *dNf1^R1861C^* (light blue). **B.** Quantification and representative Western blot of phospho-MAPK (pMAPK) relative levels in adult head lysates. One-way ANOVA revealed a significant difference across genotypes (F_(3,29)_ = 6.998, p = 0.0011). Levels were highly significantly elevated in *dNf1^Null^* (p =0.0003, 3 stars) relative to *w^1118^*controls but to a lesser extent for *dNf1^C1045Y^* (p =0.019, single star) and *dNf1^R1861C^* (p = 0.022, single star). Tubulin (Tub) was used as a loading control because MAPK levels were found not significantly different among genotypes (Suppl Fig5). Full statistical details in Suppl Table 1. **C.** An actogram plotting the mean sleep ratio per 60 min interval over a 48-hour period. Open and filled horizontal bars on the x-axis represent daytime and nighttime (zeitgeber time in hours), respectively. **D-I.** Stars indicate that the mutant score is significantly different than that of controls, the double dagger signifies differences in controls or mutants in the day relative to night-time scores, and the cross denotes significant differences among the mutants. **D.** Total sleep duration accumulated over the monitored period. Kruskal-Wallis testing demonstrated highly significant variation among groups. Total sleep was severely compromised in *dNf1^Nul^*^l^ (p <0.0001), *dNf1^C1045Y^* (p < 0.0001), and *dNf1^R1861C^* (p = 0.0009) compared to *w^1118^* controls (asterisks). Sleep reduction in *dNf1^C1045Y^* was significantly more pronounced than in *dNf1^R1861C^* (Mann-Whitney U = 2817, p < 0.0001, cross). **E.** Total sleep duration during Day and Night periods. Group variations were highly significant during both the day (H_(3)_ = 91.24, p <0.0001) and night (H_(3)_ = 129.1, p < 0.0001). Daytime sleep was reduced in *dNf1^Null^*(p < 0.0001) and *dNf1^C1045Y^* (p = 0.0274), while *dNf1^R1861C^* was indistinguishable from controls (p > 0.9999). At night, all three mutants exhibited significantly less sleep than controls (p < 0.0001, asterisks). *dNf1^C1045Y^*flies slept significantly less than *dNf1^R1861C^* flies during both day and night intervals (p =0.0002 and p =0.0001, respectively). **F.** Sleep bout number quantified across the day and night intervals. Significant overall differences were identified during both phases (p <0.0001). *dNf1^R1861C^* displayed a substantial increase in daytime and nighttime bout number (p <0.0001) and (p = 0.0002), whereas *dNf1^Null^* showed reduced bout number during the day (p <0.0001) and at night (p = 0.0003). *dNf1^C1045Y^* exhibit decreased bouts only during the night (p=0.0001). Cross symbols indicate a significant difference between the *dNf1^C1045Y^* and *dNf1^R1861C^* mutants during day (U = 2780, p < 0.0001) and night (U = 1563, p < 0.0001). **G.** Mean sleep bout length (in minutes). Significant overall effects occurred in daytime (H_(3)_= 87.00, p <0.0001) and nighttime (H_(3)_ = 126.8, p <0.0001). Bout length was significantly reduced in *dNf1^Null^* during both phases (p <0.0001) and in *dNf1^C1045Y^* (p =0.0285) day, (p <0.0001) night). *dNf1^R1861C^* showed severely reduced bout duration exclusively at night (p <0.0001). Cross symbols indicate a significant difference between *dNf1^C1045Y^* and *dNf1^R1861C^* variant lines as determined by a pairwise Mann-Whitney U test during day (U =3526, p=0.0034). **H.** Total activity (in minutes). Daytime activity varied explicitly by genotype (H_(3)_ =93.92, p <0.0001), where *dNf1^C1045Y^* and *dNf1^R1861C^* exhibited decreased daytime values (p < 0.0001). Night-time hyperactivity was evident in *dNf1^Null^* (p <0.0001), *dNf1^C1045Y^* (p < 0.0001) but not for *dNf1^R1861C^* (p =0.0177). Diurnal modulation analysis confirmed a drop in night-time activity relative to daytime for *w^1118^* (p =0.0001) and *dNf1^Null^* (p =0.0053), whereas *dNf1^C1045Y^* activity increased at night (p =0.0005) and *dNf1^R1861C^* remained unchanged (p = 0.2785). Cross symbols indicate a significant difference between *dNf1^C1045Y^* and *dNf1^R1861C^* as determined by a pairwise Mann-Whitney U test (U= 3337, p=0.0007). **I.** Waking activity index (Activity/Waking time). Daytime differences were notable (H_(3)_ = 100.9, p <0.0001), driven by reduced activity intensity in *dNf1^C1045Y^* and *dNf1^R1861C^* (p <0.0001 for both). Nighttime waking intensity was altered (H_(3)_ = 30.77, p <0.0001), with elevated waking activity in *dNf1^Null^* (p <0.0001) and *dNf1^C1045Y^* (p=0.0039). Cross symbols indicate a significant difference between *dNf1^C1045Y^* and *dNf1^R1861C^* mutants as determined by a pairwise Mann-Whitney U test (U=3274, p=0.0004).

In these mutants, pMAPK levels, are elevated significantly (Fig 7B), though less so than in null head lysates. (p=0.0003 for the null, 0.0191 for *dNf1^C1045Y^*and 0.0224 for *dNf1^R1861C^*). Nonetheless, sleep was altered in both missense mutants relative to controls, with *dNf1^C1045Y^* exhibiting more severe deficits in sleep pattern, whereas *dNf1^R1861C^* homozygotes retained rudimentary sleep periodicity (Fig 7C). Quantification of total sleep demonstrated that it was highly reduced in *dNf1^C1045Y^*homozygotes, approximating the levels of null homozygotes, with a significant but milder deficit for *dNf1^R1861C^* animals (Fig 7D). The deficits appeared attributable primarily to deficient night-time sleep for both missense mutants, though milder for *dNf1^R1861C^* homozygotes. However, during the day *dNf1^R1861C^* homozygotes slept as control animals did and significantly more than null and *dNf1^C1045Y^* homozygotes (Fig 7E).

Quantification of the number of sleep episodes revealed that unlike the null homozygotes, daytime sleep episodes were not significantly different in the missense mutants than those in control animals. At night-time however, whereas sleep bout number was significantly reduced for *dNf1^C1045Y^* to the levels of null flies, it was not affected in *dNf1^R1861C^* homozygotes (Fig 7F). Conversely, the length of night-time sleep episodes was highly reduced in both missense mutants, to levels similar to those exhibited by null homozygotes, but day-time sleep bouts were not significantly affected (Fig 7G). Therefore, *dNf1^C1045Y^* homozygotes present with strongly decreased and fragmented sleep, whereas *dNf1^R1861C^* animals exhibit a much milder sleep deficit and fragmentation, attributable largely to shortened nighttime sleep bouts.

Interestingly, unlike for null homozygotes daytime activity was reduced in both missense mutants, and like nulls was elevated at night-time for *dNf1^C1045Y^*, but not for *dNf1^R1861C^* homozygotes (Fig 7H, I). Collectively the data demonstrate that although null and missense homozygotes present with deficient fragmented sleep and altered activity patterns, the pattern and magnitude of the effects are discernable and broadly mutation-specific. This is further demonstrated by examining sleep in heterozygous null and missense mutants. Null and *dNf1^C1045Y^* heterozygotes did not present significant pattern (Suppl Fig 7A), or quantitative sleep deficits (Suppl Fig 7B, C). In contrast, *dNf1^R1861C/+^* flies slept significantly more during the day and less at night-time than controls and the other two mutant heterozygotes (Suppl Fig 7A-C). However, their sleep bout number and length remained within control levels (Suppl Fig 7D, E), except for their increased day-time bout length (Suppl Fig 7E). Furthermore, relative to controls and other mutant heterozygotes, *dNf1^R1861C/+^* flies were hypoactive during the day and less active at night-time (Suppl Fig 7F, G). Taken together these results indicate differential and specific effects of these missense mutations in accord with the reported variable effects of their counterparts in patients.

### Restoration of GRD activity specifically in cholinergic neurons reverses the deficient and fragmented sleep of dNf1^C1045Y^ homozygotes

Because pan-neuronal (Suppl Fig 4), or cholinergic neuron-specific (Fig 5) expression of a wild type *dNf1* transgene sufficed to reverse the deficient sleep of null homozygotes, we examined whether likewise it would also reverse the sleep deficits of *dNf1^C1045Y^* homozygotes. Surprisingly however, pan-neuronal expression of the wild type transgene did not reverse the sleep deficit (Fig 8A) of *dNf1^C1045Y^* homozygous flies. Sleep fragmentation persisted as well, evidenced by the increased (Fig 8B), but significantly shorter (Fig 8C) sleep episodes.

**Figure 8.**
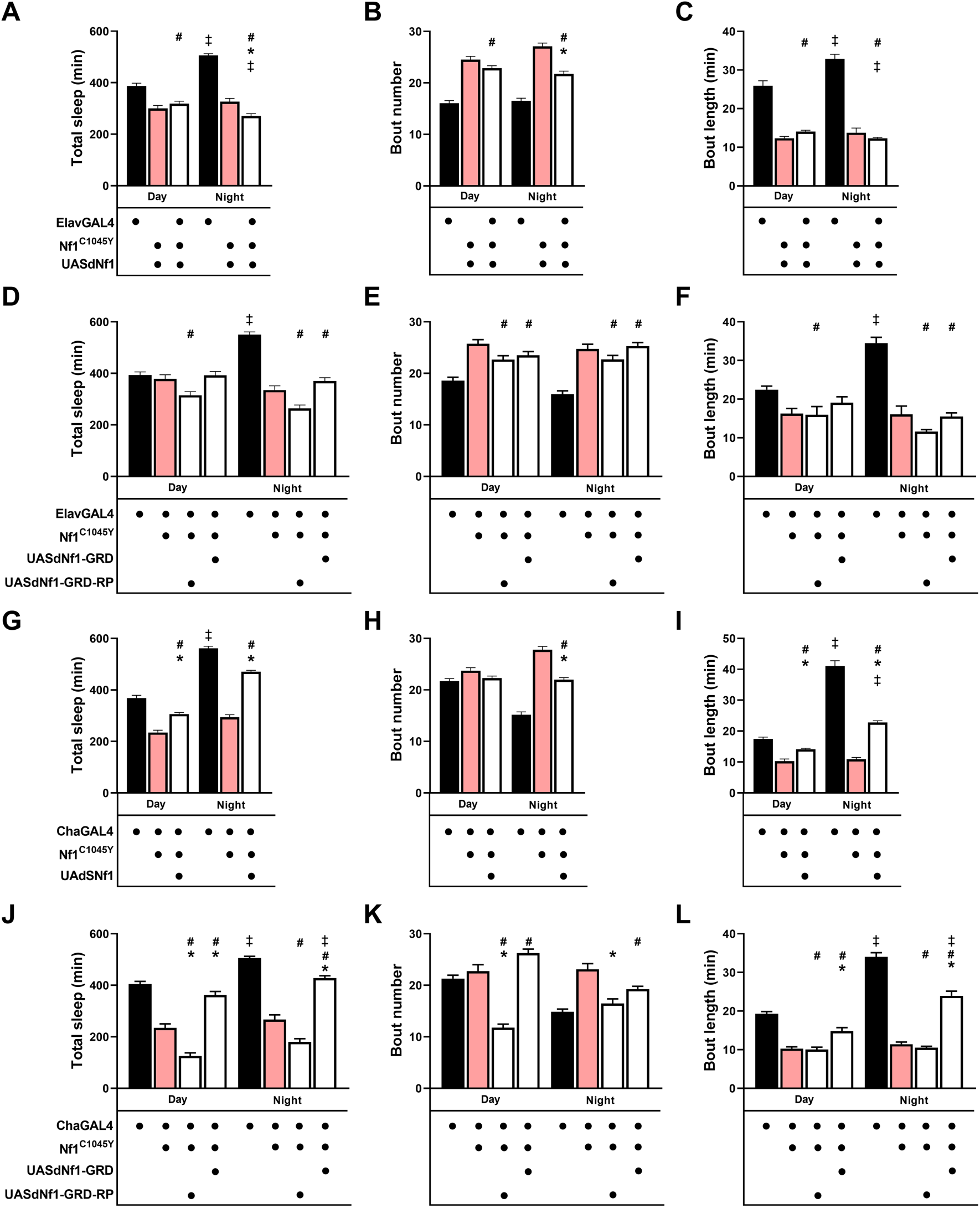
Tissue-specific rescue of the *dNf1^C1045Y^* missense mutant by wild-type *dNf1* and *dNf1-GRD* transgenes. **A–L.** Full statistical details in Suppl Table 1. Stars indicate that the mutant score is significantly different than that of controls, the double dagger signifies differences in controls or mutants in the day relative to night-time scores and # denotes significant differences of the experimental score compared to that of th controls. Controls are represented by black bars, mutants by pink and potential rescue strains by the open bars. **A-C.** Pan-neuronal expression of full-length wild-type UAS-dNf1. **A**The total time spent asleep for ElavGal4/+, during day and night, was significantly different than that for UASdNf1/+; d*Nf1^C1045Y^*(p<0.0001), but also from ElavGal4>UASdNf1/+;*dNf1^C1045Y^*(p<0.0001) indicating failure to rescue the sleep deficit. However, sleep in ElavGal4>UASdNf1/+; *dNf1^C1045Y^* was significantly decreased compared to UASdNf1/+; *dNf1^C1045Y^* (p<0.0001) at night-time. Night-time sleep was significantly elevated compared to that during the day for ElavGal4/+ (p<0.0001 double dagger) but decreased for ElavGal4>UASdNf1/+; *dNf1^C1045Y^*(p=0.0002). Sleep bout number for controls during day and night-time, was significantly different than that for the mutant UASdNf1/+; *dNf1^C1045Y^* (p<0.0001), but also from ElavGal4>UASdNf1/+; *dNf1^C1045Y^* (p<0.0001). However, bout number in ElavGal4>UASdNf1/+; *dNf1^C1045Y^* was significantly decreased compared to UASdNf1/+; *dNf1^C1045Y^* (p<0.0001, asterisk) during night-time. Mean sleep bout length presented with significant reduction during both day and night (p<0.0001) for the mutant and transgene-bearing groups relative to controls. Night-time bout length was significantly increased compared to that during the day for ElavGal4/+ (p<0.0001 double dagger) but decreased for ElavGal4>UASdNf1/+; *dNf1^C1045Y^* (p=0.002). The bout length for ElavGal4/+ during both day and night, was significantly different than that for UASdNf1/+; *dNf1^C1045Y^* (p<0.0001), but also from ElavGal4>UASdNf1/+; *dNf1^C1045Y^* (p<0.0001). **D–F.** Pan-neuronal expression of UAS-dNf1-GRD and the catalytic domain mutant UAS-dNf1-GRD-RP transgenes. Total sleep duration during the day was not significantly different among genotypes, except in mutant animals expressing the mutant transgene (p=0.0006). In contrast, night-time sleep was severely compromised in ElavGal4; *dNf1^C1045Y^* mutants (p<0.0001) compared to ElavGal4/+ controls and was not rescued by the expression of the wild type or mutant dNf1 catalytic domain transgenes (p<0.0001 compared to controls for both). Night-time sleep was significantly elevated compared to that during the day in ElavGal4/+ controls (double dagger), whereas this day–night difference was reduced or absent in the mutant and transgene-bearing groups. Sleep bout number indicating sleep fragmentation for the missense mutant and rescue groups compared to controls. ElavGal4/+ controls displayed fewer sleep bouts during the day and night compared with the *dNf1^C1045Y^*mutant and transgene-bearing groups (p<0.0001 for all). Expression of UASdNf1-GRD altered (p=0.0053) the day time number of sleep bouts relative controls, but not relative to *dNf1^C1045Y^* homozygotes (hashtags). During the night, the mutant and transgene-bearing groups also exhibited altered bout numbers (hashtags), compared with ElavGal4/+ controls, consistent with changes in sleep fragmentation. Mean sleep bout length showing significant changes in sleep consolidation across diurnal phases. Sleep bouts in ElavGal4/+ controls were longer than those in the *dNf1^C1045Y^*homozygotes (p<0.0001) and the corresponding transgene-bearing groups, particularly during the night. Expression of UASdNf1-GRD did not fully restore (p=0.0205) the reduced sleep bout length associated with the *dNf1^C1045Y^* mutation. The UASdNf1-GRD-RP construct resulted in an increase in night-time bout length relative to the mutant. Night-time bout length was significantly elevated compared with daytime values in ElavGal4/+ controls (‡), whereas this day–night increase was absent in the mutant and transgene-bearing groups. **G–I.** Cholinergic-specific expression of full-length wild-type UAS-dNf1 transgene. Total sleep duration during Day and Night. Cholinergic neuron targeted expression of the full-length wild-type protein improved significantly (p<0.0001) the nighttime sleep deficit of the mutant compared to UASNf1*; dNf1^C1045Y^* controls, but remained significantly different than sleep in controls (p<0.0001, hashtag). ChaGal4/+ controls slept significantly more at night (p<0.0001 double dagger) and the *dNf1^C1045Y^* homozygotes (p<0.0001). Expression of UASdNf1 in mutant homozygotes improved significantly their night-time sleep (p<0.0001), but significant differences remaining relative to the control. Sleep bout number during the day was relatively similar among the indicated genotypes, whereas during night-time the *dNf1^C1045Y^* mutant showed a marked increase (p<0.0001) in bout number compared with ChaGal4/+ controls. Expression of UASdNf1 altered this deficit (p<0.0001), although significant differences remained between the transgene-bearing and control groups. These changes in bout number are consistent with alterations in sleep architecture during the nighttime. Mean sleep bout length upon full-length wild-type transgene expression. ChaGal4/+ controls displayed significantly longer sleep bouts compared with both the Nf1^C1045Y^ mutant and the mutant expressing the transgene (p<0.0001 for both) during both diurnal phases. Expression of UASdNf1 improved sleep bout length in the mutant background both during the day and at night (p<0.0001) but in both cases not to control levels. Night-time bout length was significantly greater than during the day in the control group and the partially rescued mutant (double dagger). **J–L.** Cholinergic-specific expression of UAS-dNf1-GRD and catalytic domain mutant UAS-dNf1-GRD-RP transgenes. Expression of the GRD domain significantly rescued night-time sleep compared to the mutant (p<0.0001), whereas the catalytic-dead version did not have an effect. Controls showed significantly greater total sleep than the Nf1^C1045Y^ mutant during both diurnal phases. Expression of the GRD transgene rescued fully the sleep deficit (p=0.412 *vs* control, p<0.0001 *vs* mutant), whereas expression of the kinase dead GRD-RP did not. Night-time sleep was significantly elevated compared with daytime sleep in ChaGal4/+ controls and the rescued mutant (double dagger). Expression of the GRD-bearing transgene elevated bout number during the day and normalized sleep bouts at night-time similar to those of controls (p=0.0036), but not different than those of the mutant (p=0.3498). Mean sleep bout length confirmed significant recovery of night-time sleep consolidation upon GRD expression compared to the mutant or GRD-RP-expressing animals. Sleep bouts were significantly shorter in the Nf1^C1045Y^ mutant and the transgene-expressing animals during the day (p<0.0001 for all) compared with ChaGal4/+ controls, but GRD expression resulted in significantly improved bout length compared to the mutant (p=0.0056). At nighttime there is marked improvement in the GRD expressing animals (p<0.0001) relative to the mutants that remained different than controls (p=0.0014) and was different that daytime sleep as for controls (double daggers).

Although phospho-MAPK levels were mildly increased in *dNf1^C1045Y^* homozygotes (Fig 7B), we wondered whether pan-neuronal expression of the GRD domain itself might restore sleep in these animals as it did for null homozygotes (Fig 6). However, even upon expression of the *dNf1^GRD^* transgene, total sleep remained deficient (Fig 8D) and fragmented as indicated by the increased in number (Fig 8E), but highly attenuated sleep episodes, especially at night-time (Fig 8F). Statistically indistinguishable results were obtained upon expression of the inactive *dNf1^GRD-RP^* transgene (Fig 8D-F).

Because this mutation is not rescued by expression of the wild type transgene, these results suggest that the *dNf1^C1045Y^* mutation may in fact interfere with the activity of the wild type transgene and this would explain its relatively mild effect on pMAPK levels (Fig 7B). In fact, recent publications indicate that Nf1 forms dimers, at least *in vitro* and cultured vertebrate cells (Sherekar, Han et al. 2020, Lupton, Bayly-Jones et al. 2021, Naschberger, Baradaran et al. 2021), and certain variants destabilize such dimers (Young, Goldstein de Salazar et al. 2023). This notion is also supported by the fact that dNf1^C1045Y^ protein in head lysates is not reduced but surprisingly is significantly elevated (p<0.0001) over that in controls (Suppl Fig 8A). Such expression levels may not be attained by the transgenic wild type protein but even if they do, they may not support rescue if the mutant protein in the dimers interferes with their function. However, pan-neuronal expression of the wild type transgene rescued the day and night-time hypoactivity of *dNf1^C1045Y^* homozygotes (Suppl Fig 8B). This indicates that the transgenic wild type protein suffices to support for processes underlying the hypoactivity of this mutant, suggesting that dNf1 dimers are not required for regulated activity, or that the mutant dNf1^C1045Y^ protein does not interfere with the wild type protein activity in neurons mediating these behaviors.

Significantly, limiting expression of the wild type transgene to cholinergic neurons resulted in significant amelioration, but not full recovery, of night-time sleep (Fig 8G), bout number (Fig 8H) and bout length (Fig 8I). The reduced daytime activity of *dNf1^C1045Y^* homozygotes was also elevated (Suppl Fig 8C). Similar amelioration of the night-time sleep deficit (Fig 8J), decreased sleep bout number (Fig 8K) and longer sleep episodes (Fig 8L), were obtained by expression of the *dNf1^GRD^*, but not the *dNf1^GRD-RP^* transgene in cholinergic neurons.

Therefore, in accord with the results from null homozygotes (Fig 6), restoring Ras/MAPK regulation specifically within cholinergic neurons ameliorates the deficient, fragmented sleep of *dNf1^C1045Y^*homozygotes. Significantly, rescue is specific to the GAP-active dNf1^GRD^, potentially because the transgenic protein lacks domains that may mediate dimerization with its endogenous mutant counterpart. Furthermore, the failure of pan-neuronal expression to rescue the sleep deficits supports the notion that the dNf1^C1045Y^ protein does not merely reduce dNf1 activity but antagonizes the transgenic protein in a cell type-restricted manner. This is a property of this missense variant not shared by the null allele and may bear on therapeutic approaches on NF1 patients bearing this or analogous mutations.

### Sleep restoration in dNf1^R1861C^ homozygotes

Unlike the null and *dNf1^C1045Y^* homozygotes, the sleep deficits of *dNf1^R1861C^* animals are significant but milder (Fig 7). Interestingly, as for the null mutants (Suppl Fig 4), pan-neuronal expression of the wild type transgene ameliorated their night-time sleep deficit (Fig 9A) and normalized the number (Fig 9B) and duration (Fig 9C) of sleep episodes. In addition, expression of the wild type transgene normalized the activity of the mutants (not shown). Therefore, although the dNf1^R1861C^ protein was also elevated relative to controls (Suppl Fig 8A), unlike for *dNf1^C1045Y^* homozygotes (Fig 8A-C), pan-neuronal expression of the wild type transgene sufficed to ameliorate the sleep deficits of *dNf1^R1861C^* animals.

**Figure 9.**
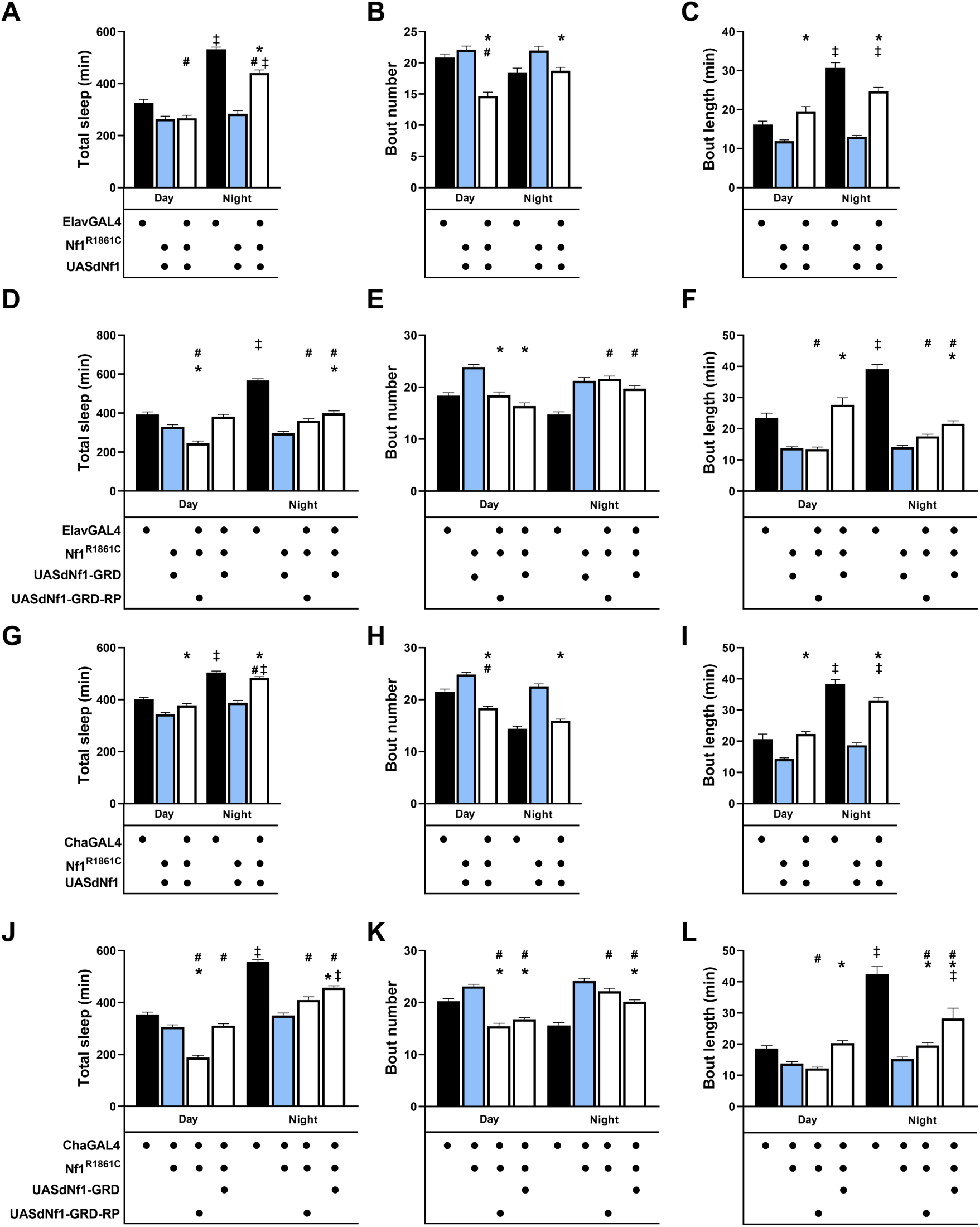
Expression of wild-type dNf1 and dNf1-GRD transgenes rescue the sleep deficits of the *dNf1^R1861C^* missense mutant. **A–L.** Full statistical details in Suppl Table 1. Stars indicate that the mutant score is significantly different than that of controls, the double dagger signifies differences in controls or mutants in the day relative to night-time scores and # denotes significant differences of the experimental score compared to that of the controls. Controls are represented by black bars, mutants by blue and potential rescue strains by the open bars. **A–C.** Pan-neuronal expression of full-length wild-type UAS-dNf1. **A.** Sleep in ElavGal4/+ controls was significantly different than both mutant and wild type transgene expressing animals during the day (p=0.0022 for both), but the transgene ameliorated the deficit at night (p<0.0001 *vs* the mutant) but not to control levels (p<0.0001). **(B)** Transgene expression significantly reduced sleep bouts during the day relative to controls and the mutant (p<0.0001 for both) and at night relative to the mutant only (p=0.0015). **(C)** Mean sleep bout length during the day was significantly different from that of the mutant upon transgene expression (p<0.0001), and similarly at night-time (p<0.0001), attaining control level (p=0.0273) and similarly to controls was elevated at night *versus* the day (p<0.0001 for both), indicating rescue of sleep fragmentation. **D–F.** Pan-neuronal expression of UAS-dNf1-GRD and catalytic domain mutant UAS-dNf1-GRD-RP transgenes. **D.** Total sleep duration across diurnal periods. Expression of the GRD fragment significantly rescued nighttime sleep deficits compared to UASdNf1-GRD; Nf1^R1861C^ controls (p<0.0001), whereas the catalytic-dead UASGRD-RP did not and in fact, significantly suppressed daytime sleep of the mutant (p<0.0001). Sleep of the mutants was improved at night-time by the GRD (p<0.0001) but not to the same extent by the GRD-RP transgene (p=0.0141). **E.** Sleep bout number was significantly reduced during the day by both transgenes compared to the mutant (p<0.0001), to control levels, but remained elevated as in the mutants at night nd significantly different from controls. **F.** Mean sleep bout length was significantly improved versus the mutant by the GRD (p<0.0001), but not GRD-RP transgene during the dau. At night-time, though improved over the mutant (p<0.0001), it remained significantly different from controls (p<0.0001), indicating partial amelioration. **G–I.** Cholinergic-specific expression of wild-type UAS-dNf1 transgene using Cha-GAL4. **G.** Expression of the wild type transgene improved (p=0.0015) daytime sleep *versus* the mutant, reaching control levels (p=0.0241). At night-time the improvement remained significantly different form the mutant (p<0.0001), but did not attain control levels (p<0.0001) and was higher than during the day as in was in controls (p<0.0001). **H.** Sleep bout number was reduced significantly relative to the mutant, but also the controls (p<0.0001 for both) during the day and similarly at night (p<0.0001 *vs* the mutant), but it attained nearly control levels (p=0.0658), indicating rescue of this sleep parameter. **I.** Mean sleep bout length was increased significantly *versus* the mutant (p<0.0001) to control levels (p=0.4667) during the day. At night a similar improvement was registered relative to the mutant (p<0.0001), approximating control levels (p=0.0127) indicating rescue of this sleep parameter. **J–L.** Cholinergic-specific expression of UAS-dNf1-GRD and the catalytic dead UAS-dNf1-GRD-RP transgenes. **J.** Total sleep duration during the day was not significantly impacted by GRD transgene expression (p=0.0026) compared to controls, or the mutant, while expression of the GRD-RP transgene significantly (p<0.0001) reduced sleep. At night-time there was a significant improvement (p<0.0001) over the mutant in the time asleep by expression of the GRD transgene, which however did not attain control levels (p<0.0001). Nighttime sleep was significantly elevated compared with daytime sleep in the control group. Nighttime sleep was significantly elevated compared with daytime sleep in the control group. **K.** Sleep bout number was significantly reduced by expression of both transgenes relative to that of the mutant (p<0.0001 for both) and remained significantly different than that of controls (p<0.0001) during the day. At night-time however, GRD, but not GRD-RP expression yielded a significant reduction of sleep bouts (p<0.0001) from the mutant in the direction of controls indicating normalization of bout numbers by GRD transgene expression. **L.** Mean sleep bout length supporting normalization of sleep consolidation in the mutant. GRD (p<0.0001), but not GRD-RP (p=0.8141) transgene expression increased bout length in the mutant during the day. The pattern held at night-time with expression only of the GRD transgene significantly (p<0.0001) increasing sleep bout length without attaining control levels (p<0.0001).

Pan-neuronal expression of the *dNf1^GRD^*transgene improved the sleep deficit of *dNf1^R1861C^* homozygotes (Fig 9D), but not at the level attained by expression of the wild type transgene (Fig 9A), largely because sleep bout number (Fig 9E) and duration (Fig 9F) were marginally changed. In contrast, expression of the mutated *dNf1^GRD-RP^* transgene did not alter the sleep deficits of the mutants (Fig 9D-F).

Expression of the wild type transgene specifically in cholinergic neurons resulted in complete reversal of the sleep deficit of *dNf1^R1861C^* homozygotes, especially at night-time (Fig 9G). This is consequent of the normalization to wild type levels of the number (Fig 9H) and duration (Fig 9I) of sleep bouts, particularly at night-time. Interestingly however, *dNf1^GRD^* transgene expression in cholinergic neurons improved significantly the sleep deficit of the mutants, especially at night-time (Fig 9J), but not to the level attained by expression of the wild type transgene (Fig 9G). This reflects the near normalization of sleep bout number (Fig 9K) and duration (Fig 9L). Expression of the *dNf1^GRD-RP^* transgene in cholinergic neurons did not have a significant impact (Fig 9 J-L).

Collectively, these results suggest that unlike in *dNf1^C1045Y^* homozygotes, sleep in *dNf1^R1861C^*animals is significantly ameliorated by pan-neuronal expression of the wild type transgene and fully rescued by its restriction to cholinergic neurons. Significantly, this occurs despite elevation of both mutant proteins (Suppl Fig 8A), demonstrating that excess mutant protein alone cannot account for the resistance of dNf1^C1045Y^ homozygotes to rescue, and supporting instead active interference by the dNf1^C1045Y^ protein with its transgenic wild type counterpart. In contrast, dNf1^R1861C^ behaves as a partial loss-of-function allele amenable to restored Ras/MAPK regulation, in accord with the milder pMAPK elevation and with the position of the mutation within the PH domain, which likely mediates membrane interactions and is distant and likely functionally upstream from the GRD and Nf1 GAP activity. Thus, although cholinergic neurons emerge as a common site of action across all three alleles, individual missense variants differ markedly in their capacity to be functionally restored.

## DISCUSSION

Our results demonstrate that loss of dNf1 results in reduced and poorly consolidated sleep in Drosophila, that this phenotype maps to cholinergic neurons rather than to the other dNf1-enriched circuits we examined, and that it depends on the Ras-GAP activity of Neurofibromin. Two missense alleles, *dNf1^C1045Y^* and *dNf1^R1861C^*, also disrupt sleep, but differ both in the parameters affected and in their response to transgenic rescue. Nf1-dependent sleep regulation therefore depends on neuronal context and on the particular molecular properties of the mutant protein that a given mutation disrupts.

Null mutants sleep approximately 60% less than controls, lack the normal night-time increase in bout number and duration and are hyperactive at night. Loss of dNf1 thus compromises both the amount and the consolidation of sleep, reproducing the two features that define sleep in NF1 patients as objectively fragmented rather than merely subjectively poor, characterized by frequent arousals and reduced sleep efficiency superimposed on shortened total sleep (Licis, Vallorani et al. 2013, Carotenuto, Messina et al. 2023). We demonstrate that the sleep deficits in Drosophila are attributable to loss of dNf1 itself as pan-neuronal knockdown reproduces them, consistent with previous studies of null and hypomorphic alleles (Bai and Sehgal 2015, Brown, Zhang et al. 2023). Significantly, knockdown restricted to cholinergic neurons reduces sleep and shortens sleep episodes without the accompanying hyperactivity, demonstrating that the sleep phenotype is not secondary to increased locomotion and that distinct neuronal populations underlie the sleep and activity components of the null phenotype.

Given the broad distribution of dNf1 in the adult brain, we asked whether sleep requires the protein within the circuits where it is enriched. Knockdown in the mushroom bodies with two independent drivers left total sleep, bout number and bout duration unaffected, whereas attenuation in all neurons except the mushroom bodies reproduced the nighttime deficit of the nulls. This is of interest because Nf1 levels oscillate in Kenyon cells and promote daytime wakefulness through cAMP/PKA signaling (Machado Almeida, Lago Solis et al. 2021), which predicts that reducing it there should increase daytime sleep. One potentially relevant difference between the studies is sex. Our experiments were performed exclusively in males while in the Machado-Almeida study primarily in virgin females, and sex-dependent differences in Nf1-associated sleep phenotypes have been reported (Bai and Sehgal 2015). Alternatively, the mushroom body contribution may be modest under our conditions, or masked by compensation within other sleep- and wake-regulatory circuits. In either case, mushroom body Nf1 is unlikely to account for the fragmentation caused by global dNf1 loss.

Our results similarly argue against a broad requirement in GABAergic neurons. Gad-Gal4-mediated knockdown did not reduce or fragment sleep, in agreement with the negative result reported for the Nf1-interacting protein Night Owl under Gad1-Gal4 (Maurer, Malita et al. 2020). Furthermore, attenuation in dorsal fan-shaped body or ellipsoid body neurons reproduce the null phenotype as sleep increased modestly in these genotypes relative to driver heterozygotes, but not relative to dNf1RNAi;Dcr-2 controls. This indicates that the effect largely arises from the RNAi background rather than from dNf1 loss in these neurons. This result bears on two reports that Nf1 knockdown under Rdl-GAL4 drivers fragments nighttime sleep, with a lower arousal threshold and loss of the sleep-dependent reduction in metabolic rate (Maurer, Malita et al. 2020, Brown, Zhang et al. 2023). The apparent discrepancy is informative rather than contradictory, because the Rdl-GAL4 driver we used (BDSC #49620) is restricted to the ellipsoid body, whereas those studies used broader drivers (#66509, #84688) that label additional Rdl-expressing populations. The fragmentation they report is therefore unlikely to originate in the ellipsoid body and is likely attributed to Rdl-positive neurons outside these neurons. Although, incomplete knockdown remains a caveat, we demonstrate that our conditions eliminate the protein in head lysates (Fig 2B). Collectively, these results narrow the circuits in which dNf1 is required for sleep and point away from several populations implicated in other Nf1-dependent behaviors.

In contrast, dNf1 attenuation in cholinergic neurons recapitulates the core features of the null phenotype, and expression of full-length dNf1 in these neurons of null homozygotes restores sleep amount, periodicity, bout number and bout duration. The incomplete recovery relative to controls is unlikely to indicate an additional requirement outside these neurons, because pan-neuronal expression o the wild type transgene does not yield complete rescue either (Suppl Fig 4), and more probably reflects the level or pattern of transgene expression. Cholinergic neurons therefore constitute a principal site of dNf1 action in sleep consolidation, which is a role for these circuits in Nf1-dependent sleep that has not previously been recognized.

How might loss of Neurofibromin from these neurons fragment sleep? A plausible interpretation is that dNf1 normally constrains cholinergic drive within a range that permits flies to initiate and sustain consolidated sleep, so that its loss shifts this balance toward excess drive and yields shorter, more frequent bouts. Such a mechanism would predict that lowering cholinergic output in the absence of dNf1 should restore consolidated sleep, a prediction that remains to be tested directly in the future. The fact that dNf1 knockdown in cholinergic neurons affects sleep without reproducing the hyperactivity of null mutants supports a division of labor, with the locomotor component of the null phenotype arising from different neuronal populations.

Our rescue experiments define the molecular activity of dNf1 required for normal consolidated sleep. The GRD alone restores pMAPK to control levels in null homozygotes and rescues sleep as effectively as the full-length protein, whereas the catalytically inactive GRD-RP construct (Klose et al., 1998; Walker et al., 2006) does not. Therefore the requirement for the Ras-GAP activity distinguishes sleep regulation from the cAMP/PKA-dependent processes implicated in growth and in mushroom body-mediated daytime wakefulness (Walker, Tchoudakova et al. 2006, Machado Almeida, Lago Solis et al. 2021), and aligns it with the role of Ras1 in regulating GABA release during learning (Georganta, Moressis et al. 2021). This conclusion does not exclude contributions from other Neurofibromin domains or from cAMP signaling in other neuronal contexts. Rather it indicates that restoring Ras regulation suffices to recover normal sleep. However how altered Ras/MAPK signaling ultimately changes cholinergic output remains an important question to be addressed in future work.

Because missense mutations account for approximately a third of known patient alleles, and genotype-phenotype correlation in NF1 is poor, we asked whether two such variants modeled in the fly gene would disrupt sleep as the null does. Our results (Fig7-Fig 9) demonstrate that they do, but not identically, and the differences are instructive. The *dNf1^C1045Y^* mutation near the Tubulin-Binding Domain, upstream of the GRD, results in a near-null sleep deficit. On the other hand, *dNf1^R1861C^* which models the human R1809C variant in the pleckstrin-homology (PH) domain, retains rudimentary periodicity and normal daytime sleep, with the deficit confined largely to shortened nighttime bouts. Importantly, the mutant heterozygotes differ as well because *dNf1^R1861C/+^*flies sleep more during the day and less at night than controls, while null and *dNf1^C1045Y^* heterozygotes are essentially unaffected. Because NF1 is a dominant disorder, and patients are typically heterozygous, this semi-dominant behavior of a relatively common variant associated principally with cognitive deficits (Pinna, Lanari et al. 2015, Rojnueangnit, Xie et al. 2015) is of particular interest.

The alleles diverge most strikingly in their response to genetic rescue. Pan-neuronal expression of full-length dNf1 restores sleep in null animals and ameliorates it in *dNf1^R1861C^* homozygotes, but fails in their *dNf1^C1045Y^* counterparts, as does pan-neuronal expression of the GRD. Critically, both mutant proteins are elevated in head lysates, so excess mutant protein cannot by itself explain why only *dNf1^C1045Y^* mutants resist rescue. The more parsimonious interpretation is active interference based on the reported Neurofibromin dimers (Sherekar, Han et al. 2020, Lupton, Bayly-Jones et al. 2021, Naschberger, Baradaran et al. 2021), which certain variants destabilize (Young, Goldstein de Salazar et al. 2023). Although Cysteine 1045 was not among the destabilizing mutations tested, the results are consistent with the notion that the dNf1^C1045Y^ protein may compromise the transgenic wild type within mixed dimers. Furthermore, the fact that pan-neuronal wild-type expression nonetheless restores the activity deficits of these animals argues against generalized interference and favors an effect confined to particular neuronal populations or processes.

Consistent with this view, restricting expression to cholinergic neurons ameliorates the night-time sleep deficit, bout number and bout duration of *dNf1^C1045Y^* homozygotes, with equivalent improvement obtained using the GAP-active GRD, but not the GRD-RP construct (Fig 8). The effectiveness of the GRD is notable because it lacks the domains implicated in dimerization, and would therefore escape interference by the endogenous mutant protein. A further comparison bears on the question of why restricting rescue to cholinergic neurons outperformed rescue with the broader pan-neuronal Elav driver. In null homozygotes, which contain no mutant protein, pan-neuronal and cholinergic rescue are equally effective. In contrast, restricting transgene expression to cholinergic neurons is consistently more effective for both missense mutants. Therefore, the advantage of rescue in the cholinergic system emerges only when the dNf1^C1045Y^ mutant Neurofibromin is present. This is not readily explained by driver strength alone and is predicted if the mutant protein antagonizes the transgenic one in a manner sensitive to their relative abundance within particular neurons.

On the other hand, the *dNf1^R1861C^* mutants present a graded rather than an all-or-none profile. Full-length dNf1 ameliorates sleep when expressed pan-neuronally and reverses the deficit completely when restricted to cholinergic neurons, whereas the GRD improves sleep only marginally pan-neuronally and substantially, in cholinergic neurons. Hence, Ras-GAP activity contributes to this phenotype but does not account for all of it, implicating an additional, GRD-independent contribution of Neurofibromin to sleep consolidation. This is the expected outcome for a partial loss-of-function allele whose lesion lies in the PH domain, distant from and functionally upstream of the GRD, where it plausibly perturbs membrane association rather than catalysis, and in in accord with the milder elevation of pMAPK in these animals.

The allele-specific effects reported here argue against treating NF1 missense variants as interchangeable partial loss-of-function alleles. *dNf1^C1045Y^* precipitates a severe, null-like sleep phenotype and resists pan-neuronal rescue, whereas *dNf1^R1861C^* yields a milder, largely night-time deficit that is readily reversed. Such differences may contribute to the variable neurological and behavioral manifestations associated with individual NF1 variants, and they bear on the design of therapeutic strategies. Accordingly, a variant that antagonizes restored wild-type protein would be expected to respond poorly to approaches aiming to supplement Neurofibromin function, whereas one that spares the GRD may respond to interventions targeting Ras/MAPK signaling, such as MAPK inhibitors.

Collectively then, our findings support a model in which Neurofibromin promotes consolidated sleep through its Ras-GAP activity within cholinergic neurons. Loss of dNf1 from these neurons suffices to reduce sleep and shorten sleep episodes, and restoration of the GRD suffices to reverse these deficits, with elevated cholinergic drive providing a candidate physiological link between the two. Furthermore, we identify a previously unrecognized role for cholinergic neurons in Nf1-dependent sleep regulation and place Ras-GAP activity therein as a central component of Nf1-dependent sleep regulation, while revealing allele-specific effects that distinguish sleep quantity from sleep consolidation. More broadly, they indicate that the behavioral consequences of Nf1 loss depend not only on how much functional Neurofibromin remains, but on where and how its molecular functions are disrupted.

## MATERIALS AND METHODS

### Drosophila culture and strains

Drosophila crosses were set up in standard wheat-flour-sugar food supplemented with soy flour and CaCl_2_ (Acevedo, Froudarakis et al. 2007) and cultured at 25°C and 50%-70% humidity with a 12 h light/dark cycle. The *dNf1^Null^* and *dNf1^C1045Y^* mutants, which contain EMS-induced nonsense and missense mutations, respectively, within the NF1 gene, have been described previously (Walker, Tchoudakova et al. 2006, Gouzi, Moressis et al. 2011, Georganta, Moressis et al. 2021). Novel *dNf1* mutants are described below.

Transgenic fly strains used in this work were as follows: UAS-*dNf1* (Walker, Tchoudakova et al. 2006), UAS-*dNf1^RNAi^*(Vienna Drosophila RNAi Center-VDRC, stock #109637), UAS-*Dcr-2* (BDRC stock #24651), UAS-247Gal80 (or Mef2Gal80, henceforth MBGal80, BDRC stock #64306). The UAS-*dNF1^GRD^*(*dNF1^GRD^*) and UAS-*dNF1^GRDR1320P^* (*dNF1^GRD-RP^*) have been described before (Walker, Tchoudakova et al. 2006). The Gal4-driver lines used in this study were Leo-Gal4 (Messaritou, Leptourgidou et al. 2009), Cha-Gal4 (BDRC stock #6798), Elav^C155^-Gal4 (BDSC stock #458), 247-Gal4 (BDRC stock # 27390), Gad-Gal4 (BDRC stock #51630), R23E10-Gal4 (BDRC stock #490323), Rdl-Gal4 (BDRC stock #49620). The novel dNf1-CRIMIC line containing a CRIMIC element in the 3’end of the gene was kindly provided by H. Bellen (Baylor College of Medicine Houston, TX USA). All strains were backcrossed into the resident Cantonised-*w^1118^* isogenic background for six generations.

### Generation of novel CRISPR/Cas9-mediated *dNf1* alleles

The *dNf1*^Δ216^ allele is a 261 bp deletion generated by CRISPR/Cas9-induced non-homologous end-joining. It removes 103 bp upstream of the ATG in exon 1 and 155 bp downstream extending into intron 1. sgRNA target sites were identified using the flyCRISPR target finder (flycrispr.org) (Gratz, Ukken et al. 2014). The two sgRNAs employed were: sgRNA1 NF1: GGGAGTGGCAATTGGTTGAGTGG and sgRNA2 NF1: GTCGCTGTGAGCATGTGTGGGGG, which were cloned into the pBFv-U6.2 expression vector (Genome Engineering Production Group at Harvard Medical School) for injection. The deleted sequence was confirmed as: 5’ACTCAACCAATTGCCTG CTGAAATGCAGTAAAAAACGCATAAACAACTGTTAAAACACCACAACAAGATGACCCAGAA GGAGGATCAGGCAAGTGTTTTCCAATCGCTTTTCCGCAGTTGCGTTTTCGCAAGCGAAAAAA TGTCACAGCGACC-3’. A validated mutant line was backcrossed for 6 generations to third chromosome balancers in the resident Cantonised *w^1118^* genetic background and the mutation-bearing chromosome was maintained balanced over TM3,Sb^1^.

The *dNf1^R1861C^* is also a CRISPR/Cas9-mediated missense mutation (Bassett, Tibbit et al. 2013), generated in the Drosophila *Nf1* gene at the position equivalent to the human gene R1809C, which has been associated largely with cognitive deficits. It was generated as follows: A sgRNA TAGTCAGCCGGACTCCGTGA-containing plasmid for the *dNf1* gene and donor oligonucleotide template were injected into NosCas9 attp40 expressing fly embryos (*w^1118^*; PBac{y[+mDint2]=vasCas9}VK00027) by Rainbow Transgenic Flies, Inc, (Camarillo, CA USA). The donor sequence contained the point mutation of interest, a silent mutation in the PAM site and an additional silent mutation generating a unique restriction site suitable for genotyping. Adults that emerged from the injected embryos were crossed with TM2/TM6B. Animals containing the R1809C point mutation were identified by the presence of the unique restriction site following amplification of the relevant genomic region by the polymerase chain reaction (PCR) with the primers: FWD: ACTGATTTCATCGACGCGGA, REV: AGTAGCGGTCAAAGCACACA and verified by sequencing. Validated lines containing only the sought missense mutation were backcrossed for 6 generations to third chromosome balancers in the resident Cantonised *w^1118^* genetic background and the mutation-bearing chromosome was maintained balanced over TM3,Sb^1^.

### Sleep and locomotor activity assay

2-5 day-old male flies were collected under CO_2_ into fresh standard food vials and maintained at 25°C, a 12:12 h light/dark (LD) cycle and 50% humidity for entraining at least two days prior to being assayed. Sleep assays were performed in the DAM5H Drosophila Activity Monitors (Trikinetics, Waltham, MA, USA).

Individual flies were placed in the 65 ×5mm glass assay tubes containing on one end a small plug, about 1 cm thick, of 1.5% agarose with 3% brown sugar, 1.5% fructose, and 0.5% Nipagin and a standard black cap on the other end per manufacturer’s instructions. To be placed in the tubes the flies were immobilized on ice. The tubes were placed into DAM monitors in a purpose designed incubator kept at 25°C at 12:12 LD. Monitors were kept perpendicular to and equidistant from, the white LED light source in the incubator. Flies were allowed to acclimatize to activity monitors and food for 16h before data acquisition for the following 2 days at 12:12 LD at 25 °C.

Sleep and waking activity were measured as previously described (Hendricks, Finn et al. 2000, Shaw, Cirelli et al. 2000). For each individual fly, the DAM system measures activity by counting the number of infrared beam crossings over time. Activity was measured in 1 min bins and sleep was identified as 5 min of consolidated inactivity. Data was processed using *SleepMat* (Sisobhan, Rosensweig et al. 2022) and *Rtivity* software (Silva, Pinho et al. 2022). The total time per hour flies spend sleeping on average throughout the subjective day and night day is represented by the Sleep Ratio/60 min and used to construct the respective graphs. The sum of time sleeping during the time assayed per fly, averaged over the number of flies per experiment is represented by the Total Sleep during the 48 hrs the animals were assayed, and conversely can be divided in total sleep during the day or at night. The average number of sleep bouts per day or night for all animals assayed is quantified by the bout number, while the mean length of each bout averaged over the animals assayed is reported as bout length. The total time the animals spend moving in the assay chamber averaged over all animals assayed is reported as the Total Activity, while the total time spend moving between sleep bouts averaged over all animals assayed is quantified as Activity per Waking Time.

### Western blot analyses

Eight fly heads, 2-5 d after eclosion were homogenized in 1X Laemmli buffer (50 mM Tris, pH 6.8, 100 mM DTT, 5% 2-mercaptoethanol, 2% SDS, 10% glycerol, and 0,01% bromophenol blue) and loaded per lane. Proteins were transferred to PVDF membrane and probed with the following antibodies: mouse anti-β-tubulin E7 (Developmental Studies Hybridoma Bank-DSHB) at 1:5000, mouse anti-syntaxin 8C3 rabbit (DSHB) at 1:5000, rabbit anti-phospho-p44/42 MAPK (Thr202/Tyr204) (#4370, Cell Signaling Technology) at 1:500, and rabbit anti-44/42 total MAPK (Erk1/2) (#9102, Cell Signaling Technology) at 1:500. Appropriate HRP-conjugated secondary antibodies were applied at 1:5000. Proteins were visualized with chemiluminescence (Immobilon Crescendo, Millipore).

### Confocal imaging

The distribution of dNf1 in the adult head was reported through Nf1-CRIMIC driven UAS-GFP expression. Confocal imaging was performed as described previously (Georganta, Moressis et al. 2021). In brief, dissected brains were fixed in 4% paraformaldehyde/1xPBS for 20min at room temperature. Brains were then washed with PBS + 0.3% Triton X-100 (PBT) to remove residual fixative. Samples were mounted on a slide using Dako fluorescent mounting medium and imaged with a Leica Microsystems TCS SP8 Confocal Microscope under 40x oil objective lens. Images were formatted using Fiji software and Adobe Photoshop CS2.

### Proteomic determination of dNf1 and MAPK levels via quantitative proteomics

Three biological and two to three technical replicates from each genotype were used for this experiment, as described previously (Papanikolopoulou, Roussou et al. 2019). Briefly, 10 fly heads or dissected brains per genotype (*w^1118^, dNf1^null^*, Elav-Gal4>+, Elav-Gal4>UAS-Dcr2, Elav-Gal4>UAS-dNf1^RNAi^, Elav-Gal4> UAS-dNf1^RNAi^; UAS-Dcr2) were homogenized and lysed in 4%SDS, 0.1M DTT and 0.1M Triethylammonium bicarbonate buffer (TEAB) followed by boiling for 3 min. Samples were sonicated in a sonicating water bath for 10 min and centrifuged at 13000rpm for 10 min at room temperature. The supernatant was processed using the single-pot, solid-phase-enhanced sample preparation Sp3-mediated protein digestion with Platinum grade Trypsin (Promega). The next day, the supernatants were collected and the peptides were purified using a modified Sp3 clean-up protocol and finally solubilized in the mobile phase A (0.1% formic acid in water), sonicated and the peptide concentration was determined in NanoDrop (absorbance at 280 nm).

Samples were run on a liquid chromatography-tandem mass spectrometry (LC-MS/MS) setup consisting of a Dionex UltimateRSLC online with a Thermo Q Exactive HF-X Orbitrap mass spectrometer (Thermo Scientific, Waltham, MA, USA). Peptide samples were directly injected and separated on a 25 cm-long analytical C18 column (1.9 µm ^3^ beads, 75 µm ID, Pepsep) using a 1-hour long run, starting with a gradient consisting of 7% Buffer B (0.1% formic acid in 80% acetonitrile) to 35% for 40 min and followed by an increase to 45% in 5 min and a second increase to 99% in 0.5 min and then kept constant for equilibration for 14.5 min. The flow rate was set to 400 nL/min in the sample loading phase and lowered to 250 nL/min in the main separation phase of sample analysis. A full MS was acquired in profile and positive mode using a mass spectrometer, operating in the scan range of 375–1400 m/z using 120 K resolving power with an AGC of 3 × 10^6^ and max IT of 60 ms followed by data independent analysis using 8 Th windows (39 loop counts) with 15 K resolving power with an AGC of 3 × 10^5^ and max IT of 22 ms and a normalized collision energy of 26.

Orbitrap raw data were analyzed in DIA-NN 2.0 (Data-Independent Acquisition by Neural Networks) through searching against the *Drosophila melanogaster* Uniprot reference database containing 13892 proteins using the library-free mode of the software and allowing up to two tryptic missed cleavages. The match-between-runs feature was used for all analyses and the output (precursor) was filtered at 0.01 FDR. The protein inference was performed on the level of genes using only proteotypic peptides. dNf1 and MAPK protein abundance was calculated on the basis of the normalized spectral protein intensity as label-free quantitation (LFQ intensity). Visualization was performed in Perseus (version 1.6.15.0) and statistical analysis using GraphPad Prism (v.8.1) as described below.

### Statistics and reproducibility

GraphPad Prism (v.8.1) was used for statistical analysis and data plotting. All experiments were performed at least in independent triplicates, irrespective of the total number of animals examined per genotype in each set. Data was tested for normal ditribution with Shapiro-Wilk tests with α<0.05 leading to nonparametric analysis of the entire data set. Normally distributed data were analysed parametrically using Welch’s or Dunnett’s tests as appropriate following initail possitive ANOVA. Conversely, non-normally distributed data were alanysed non-parametrically using Kruskal-Wallis and Mann-Whitney tests. Significance cuttoff was kept at p<0.005, although in the majority of cases was much below that as indicated along with all relevant statistical details in Supplemental Table 1 and in the figure legends as appropriate.

## ACKNOWLEDGEMENTS

The authors would like to thank Dr. Gregg Roman for help establishing the *dNf1^R1681C^*, Maro Loizou for technical help and the Bloomington Drosophila Stock Center for strains. This work was supported by Grants from the US Department of Defense Neurofibromatosis program (W81XWH-20-1-0255) and the Children’s Tumor Foundation Grant (CTF-2022-05-004) to EMCS.

## REFERENCES

Acevedo, S. F., E. I. Froudarakis, A. Kanellopoulos and E. M. Skoulakis (2007). "Protection from premature habituation requires functional mushroom bodies in Drosophila." Learn Mem 14(5): 376–384.

Allada, R., C. Cirelli and A. Sehgal (2017). "Molecular Mechanisms of Sleep Homeostasis in Flies and Mammals." Cold Spring Harb Perspect Biol 9(8).

Artiushin, G. and A. Sehgal (2020). "The Glial Perspective on Sleep and Circadian Rhythms." Annu Rev Neurosci 43: 119–140.

Aso, Y., K. Grübel, S. Busch, A. B. Friedrich, I. Siwanowicz and H. Tanimoto (2009). "The Mushroom Body of Adult Drosophila Characterized by GAL4 Drivers." Journal of Neurogenetics 23(1-2): 156–172.

Atsoniou, K., E. Giannopoulou, E. M. Georganta and E. M. C. Skoulakis (2024). "Drosophila Contributions towards Understanding Neurofibromatosis 1." Cells 13(8).

Bai, L. and A. Sehgal (2015). "Anaplastic Lymphoma Kinase Acts in the Drosophila Mushroom Body to Negatively Regulate Sleep." PLoS Genet 11(11): e1005611.

Bassett, A., C. Tibbit, C. Ponting and J. Liu (2013). "Highly efficient targeted mutagenesis of Drosophila with the CRISPR/Cas9 system." Cell Rep. 4(1): 220–228.

Beckwith, E. J. and A. S. French (2019). "Sleep in Drosophila and Its Context." Front Physiol 10: 1167.

Bergoug, M., M. Doudeau, F. Godin, C. Mosrin, B. Vallee and H. Benedetti (2020). "Neurofibromin Structure, Functions and Regulation." Cells 9(11).

Bergqvist, C., A. Servy, L. Valeyrie-Allanore, S. Ferkal, P. Combemale, P. Wolkenstein and N. F. F. Network (2020). "Neurofibromatosis 1 French national guidelines based on an extensive literature review since 1966." Orphanet J Rare Dis 15(1): 37.

Blum, I. D., B. Bell and M. N. Wu (2018). "Time for Bed: Genetic Mechanisms Mediating the Circadian Regulation of Sleep." Trends in Genetics 34(5): 379–388.

Bonnet, M. H. and D. L. Arand (2003). "Clinical effects of sleep fragmentation versus sleep deprivation." Sleep Med Rev 7(4): 297–310.

Brown, E. B., J. Zhang, E. Lloyd, E. Lanzon, V. Botero, S. Tomchik and A. C. Keene (2023). "Neurofibromin 1 mediates sleep depth in Drosophila." PLoS Genet 19(12): e1011049.

Carotenuto, M., G. Messina, M. Esposito, C. Santoro, D. Iacono and K. Spruyt (2023). "Polysomnographic study in pediatric neurofibromatosis type 1." Front Neurol 14: 1213430.

Chaturvedi, R. and P. Emery (2024). "Fly into tranquility: GABA’s role in Drosophila sleep." Curr Opin Insect Sci 64: 101219.

Chen, J., Z. Li, Y. Wu, X. Li, Z. Chen, P. Chen, Y. Ding, C. Wu and L. Hu (2024). "Identification of Pathogenic Missense Mutations of NF1 Using Computational Approaches." J Mol Neurosci 74(4): 94.

Crittenden, J. R., E. M. Skoulakis, K. A. Han, D. Kalderon and R. L. Davis (1998). "Tripartite mushroom body architecture revealed by antigenic markers." Learn Mem 5(1-2): 38–51.

Delidakis, C., M. Monastirioti and S. S. Magadi (2014). Chapter Six -E(spl): Genetic, Developmental, and Evolutionary Aspects of a Group of Invertebrate Hes Proteins with Close Ties to Notch Signaling. Current Topics in Developmental Biology. R. Taneja, Academic Press. 110: 217–262.

Dissel, S. (2020). "Drosophila as a Model to Study the Relationship Between Sleep, Plasticity, and Memory." Front Physiol 11: 533.

Donlea, J. M., D. Pimentel and G. Miesenböck (2014). "Neuronal machinery of sleep homeostasis in Drosophila." Neuron 81(4): 860–872.

Donlea, J. M., D. Pimentel, C. B. Talbot, A. Kempf, J. J. Omoto, V. Hartenstein and G. Miesenböck (2018). "Recurrent Circuitry for Balancing Sleep Need and Sleep." Neuron 97(2): 378–389.e374.

Durkin, J., A. R. Poe, S. J. Belfer, A. Rodriguez, S. H. Tang, J. A. Walker and M. S. Kayser (2023). "Neurofibromin 1 regulates early developmental sleep in Drosophila." Neurobiol Sleep Circadian Rhythms 15: 100101.

Eckstein, N., A. S. Bates, A. Champion, M. Du, Y. Yin, P. Schlegel, A. K.-Y. Lu, T. Rymer, S. Finley-May, T. Paterson, R. Parekh, S. Dorkenwald, A. Matsliah, S.-C. Yu, C. McKellar, A. Sterling, K. Eichler, M. Costa, S. Seung, M. Murthy, V. Hartenstein, G. S. X. E. Jefferis and J. Funke (2024). "Neurotransmitter classification from electron microscopy images at synaptic sites in Drosophila melanogaster." Cell 187(10): 2574–2594.e2523.

Fahsold, R., S. Hoffmeyer, C. Mischung, C. Gille, C. Ehlers, N. Kücükceylan, M. Abdel-Nour, A. Gewies, H. Peters, D. Kaufmann, A. Buske, S. Tinschert and P. Nürnberg (2000). "Minor Lesion Mutational Spectrum of the Entire NF1 Gene Does Not Explain Its High Mutability but Points to a Functional Domain Upstream of the GAP-Related Domain." The American Journal of Human Genetics 66(3): 790–818.

Fiala, A. and K. R. Kaun (2024). "What do the mushroom bodies do for the insect brain? Twenty-five years of progress." Learn Mem 31(5).

Foka, K., E. Georganta, O. Semelidou and E. Skoulakis (2022). "Loss of the Schizophrenia-linked Furin protein from Drosophila mushroom body neurons results in antipsychotic-reversible habituation deficits." J Neurosci. 42(39): 7496–7511.

Georganta, E. M., A. Moressis and E. M. C. Skoulakis (2021). "Associative Learning Requires Neurofibromin to Modulate GABAergic Inputs to Drosophila Mushroom Bodies." J Neurosci 41(24): 5274–5286.

Gouzi, J. Y., A. Moressis, J. A. Walker, A. A. Apostolopoulou, R. H. Palmer, A. Bernards and E. M. Skoulakis (2011). "The receptor tyrosine kinase Alk controls neurofibromin functions in Drosophila growth and learning." PLoS Genet 7(9): e1002281.

Gratz, S. J., F. P. Ukken, C. D. Rubinstein, G. Thiede, L. K. Donohue, A. M. Cummings and K. M. O’Connor-Giles (2014). "Highly specific and efficient CRISPR/Cas9-catalyzed homology-directed repair in Drosophila." Genetics 196(4): 961–971.

Harrisingh, M. C. and A. C. Lloyd (2004). "Ras/Raf/ERK signalling and NF1." Cell Cycle 3(10): 1255–1258.

Hasegawa, T., J. Tomita, R. Hashimoto, T. Ueno, S. Kume and K. Kume (2017). "Sweetness induces sleep through gustatory signalling independent of nutritional value in a starved fruit fly." Sci Rep 7(1): 14355.

Hendricks, J. C., S. M. Finn, K. A. Panckeri, J. Chavkin, J. A. Williams, A. Sehgal and A. I. Pack (2000). "Rest in Drosophila is a sleep-like state." Neuron 25(1): 129–138.

Johannessen, C. M., E. E. Reczek, M. F. James, H. Brems, E. Legius and K. Cichowski (2005). "The NF1 tumor suppressor critically regulates TSC2 and mTOR." Proc Natl Acad Sci U S A 102(24): 8573–8578.

Johnson, H., L. Wiggs, G. Stores and S. M. Huson (2005). "Psychological disturbance and sleep disorders in children with neurofibromatosis type 1." Dev Med Child Neurol 47(4): 237–242.

Joiner, W. J. (2016). "Unraveling the Evolutionary Determinants of Sleep." Curr Biol 26(20): R1073–R1087.

Joiner, W. J., A. Crocker, B. H. White and A. Sehgal (2006). "Sleep in Drosophila is regulated by adult mushroom bodies." Nature 441(7094): 757–760.

Kanca, O., J. Zirin, Y. Hu, B. Tepe, D. Dutta, W.-W. Lin, L. Ma, M. Ge, Z. Zuo, L.-P. Liu, R. W. Levis, N. Perrimon and H. J. Bellen (2022). "An expanded toolkit for Drosophila gene tagging using synthesized homology donor constructs for CRISPR-mediated homologous recombination." eLife 11: e76077.

Keene, A. C. and E. R. Duboue (2018). "The origins and evolution of sleep." J Exp Biol 221(Pt 11).

Klose, A., M. R. Ahmadian, M. Schuelke, K. Scheffzek, S. Hoffmeyer, A. Gewies, F. Schmitz, D. Kaufmann, H. Peters, A. Wittinghofer and P. Nürnberg (1998). "Selective disactivation of neurofibromin GAP activity in neurofibromatosis type 1." Hum Mol Genet 7(8): 1261–1268.

Koczkowska, M., T. Callens, Y. Chen, A. Gomes, A. D. Hicks, A. Sharp, E. Johns, K. A. Uhas, L. Armstrong, K. A. Bosanko, D. Babovic-Vuksanovic, L. Baker, D. G. Basel, M. Bengala, J. T. Bennett, C. Chambers, L. K. Clarkson, M. Clementi, F. M. Cortés, M. Cunningham, M. D. D’Agostino, M. B. Delatycki, M. C. Digilio, L. Dosa, S. Esposito, S. Fox, M. L. Freckmann, C. Fauth, T. Giugliano, S. Giustini, A. Goetsch, Y. Goldberg, R. S. Greenwood, C. Griffis, K. W. Gripp, P. Gupta, E. Haan, R. K. Hachen, T. L. Haygarth, C. Hernández-Chico, K. Hodge, R. J. Hopkin, L. Hudgins, S. Janssens, K. Keller, G. Kelly-Mancuso, A. Kochhar, B. R. Korf, A. M. Lewis, J. Liebelt, A. Lichty, R. H. Listernick, M. J. Lyons, I. Maystadt, M. Martinez Ojeda, C. McDougall, L. K. McGregor, D. Melis, N. Mendelsohn, M. J. M. Nowaczyk, J. Ortenberg, K. Panzer, J. G. Pappas, M. E. Pierpont, G. Piluso, V. Pinna, E. K. Pivnick, D. A. Pond, C. M. Powell, C. Rogers, N. Ruhrman Shahar, S. L. Rutledge, V. Saletti, S. A. Sandaradura, C., Santoro, U. A. Schatz, A. Schreiber, D. A. Scott, E. A. Sellars, R. Sheffer, E. Siqveland, J. M. Slopis, R. Smith, A. Spalice, D. W. Stockton, H. Streff, A. Theos, G. E. Tomlinson, G. Tran, P. L. Trapane, E. Trevisson, N. J. Ullrich, J. Van den Ende, S. A. Schrier Vergano, S. E. Wallace, M. F. Wangler, D. D. Weaver, K. H. Yohay, E. Zackai, J. Zonana, V. Zurcher, K. B. M. Claes, M. Eoli, Y. Martin, K. Wimmer, A. De Luca, E. Legius and L. M. Messiaen (2020). "Clinical spectrum of individuals with pathogenic NF1 missense variants affecting p.Met1149, p.Arg1276, and p.Lys1423: genotype-phenotype study in neurofibromatosis type 1." Hum Mutat 41(1): 299–315.

Koczkowska, M., T. Callens, A. Gomes, A. Sharp, Y. Chen, A. D. Hicks, A. S. Aylsworth, A. A. Azizi, D. G. Basel, G. Bellus, L. M. Bird, M. A. Blazo, L. W. Burke, A. Cannon, F. Collins, C. DeFilippo, E. Denayer, M. C. Digilio, S. K. Dills, L. Dosa, R. S. Greenwood, C. Griffis, P. Gupta, R. K. Hachen, C. Hernández-Chico, S. Janssens, K. J. Jones, J. T. Jordan, P. Kannu, B. R. Korf, A. M. Lewis, R. H. Listernick, F. Lonardo, M. J. Mahoney, M. M. Ojeda, M. T. McDonald, C. McDougall, N. Mendelsohn, D. T. Miller, M. Mori, R. Oostenbrink, S. Perreault, M. E. Pierpont, C. Piscopo, D. A. Pond, L. M. Randolph, K. A. Rauen, S. Rednam, S. L. Rutledge, V. Saletti, G. B. Schaefer, E. K. Schorry, D. A. Scott, A. Shugar, E. Siqveland, L. J. Starr, A. Syed, P. L. Trapane, N. J. Ullrich, E. G. Wakefield, L. E. Walsh, M. F. Wangler, E. Zackai, K. B. M. Claes, K. Wimmer, R. van Minkelen, A. De Luca, Y. Martin, E. Legius and L. M. Messiaen (2019). "Expanding the clinical phenotype of individuals with a 3-bp in-frame deletion of the NF1 gene (c.2970_2972del): an update of genotype-phenotype correlation." Genet Med 21(4): 867–876.

Koczkowska, M., Y. Chen, T. Callens, A. Gomes, A. Sharp, S. Johnson, M. Hsiao, Z. Chen, M. Balasubramanian, C. Barnett, T. Becker, S. Ben-Shachar, D. Bertola, J. Blakeley, E. Burkitt-Wright, A. Callaway, M. Crenshaw, K. Cunha, M. Cunningham, M. D’Agostino, K. Dahan, A. De Luca, A. Destrée, R. Dhamija, M. Eoli, D. Evans, P. Galvin-Parton, J. George-Abraham, K. Gripp, J. Guevara-Campos, N. Hanchard, C. Hernández-Chico, L. Immken, S. Janssens, K. Jones, B. Keena, A. Kochhar, J. Liebelt, A. Martir-Negron, M. Mahoney, I. Maystadt, C. McDougall, M. McEntagart, N. M. Mendelsohn, DT, G. Mortier, J. Morton, J. Pappas, S. Plotkin, D. Pond, K. Rosenbaum, K. Rubin, L. Russell, L. Rutledge, V. Saletti, R. Schonberg, A. Schreiber, M. Seidel, E. Siqveland, D. Stockton, E. Trevisson, N. Ullrich, M. Upadhyaya, R. van Minkelen, H. Verhelst, M. Wallace, Y. Yap, E. Zackai, J. Zonana, V. Zurcher, K. Claes, Y. Martin, B. Korf, E. Legius and L. Messiaen (2018). "Genotype-Phenotype Correlation in NF1: Evidence for a More Severe Phenotype Associated with Missense Mutations Affecting NF1 Codons 844-848." Am J Hum Genet. 102(1): 69–87.

Leschziner, G. D., J. F. Golding and R. E. Ferner (2013). "Sleep disturbance as part of the neurofibromatosis type 1 phenotype in adults." Am J Med Genet A 161A(6): 1319–1322.

Licis, A. K., A. Vallorani, F. Gao, C. Chen, J. Lenox, K. A. Yamada, S. P. Duntley and D. H. Gutmann (2013). "Prevalence of Sleep Disturbances in Children With Neurofibromatosis Type 1." J Child Neurol 28(11): 1400–1405.

Lupton, C. J., C. Bayly-Jones, L. D’Andrea, C. Huang, R. B. Schittenhelm, H. Venugopal, J. C. Whisstock, M. L. Halls and A. M. Ellisdon (2021). "The cryo-EM structure of the human neurofibromin dimer reveals the molecular basis for neurofibromatosis type 1." Nat Struct Mol Biol 28(12): 982–988.

Ly, S., A. I. Pack and N. Naidoo (2018). "The neurobiological basis of sleep: Insights from Drosophila." Neurosci Biobehav Rev 87: 67–86.

Machado Almeida, P., B. Lago Solis, L. Stickley, A. Feidler and E. Nagoshi (2021). "Neurofibromin 1 in mushroom body neurons mediates circadian wake drive through activating cAMP-PKA signaling." Nat Commun 12(1): 5758.

Marana Perez, A. I., A. Duat Rodriguez, V. Soto Insuga, J. Dominguez Carral, V. Puertas Martin and L. Gonzalez Gutierrez Solana (2015). "Prevalence of sleep disorders in patients with neurofibromatosis type 1." Neurologia 30(9): 561–565.

Martin, G. A., D. Viskochil, G. Bollag, P. C. McCabe, W. J. Crosier, H. Haubruck, L. Conroy, R. Clark, P. O’Connell, R. M. Cawthon and et al. (1990). "The GAP-related domain of the neurofibromatosis type 1 gene product interacts with ras p21." Cell 63(4): 843–849.

Maurer, G. W., A. Malita, S. Nagy, T. Koyama, T. M. Werge, K. A. Halberg, M. J. Texada and K. Rewitz (2020). "Analysis of genes within the schizophrenia-linked 22q11.2 deletion identifies interaction of night owl/LZTR1 and NF1 in GABAergic sleep control." PLoS Genet 16(4): e1008727.

Messaritou, G., F. Leptourgidou, M. Franco and E. M. Skoulakis (2009). "A third functional isoform enriched in mushroom body neurons is encoded by the Drosophila 14-3-3zeta gene." FEBS Lett 583(17): 2934–2938.

Moodley, M. and K. R. Lopez (2024). "Neurofibromatosis type 1 -an update." Semin Pediatr Neurol 52: 101172.

Naschberger, A., R. Baradaran, B. Rupp and M. Carroni (2021). "The structure of neurofibromin isoform 2 reveals different functional states." Nature 599(7884): 315–319.

Papanikolopoulou, K., I. G. Roussou, J. Y. Gouzi, M. Samiotaki, G. Panayotou, L. Turin and E. M. C. Skoulakis (2019). "Drosophila Tau Negatively Regulates Translation and Olfactory Long-Term Memory, But Facilitates Footshock Habituation and Cytoskeletal Homeostasis." J Neurosci 39(42): 8315–8329.

Pinna, V., V. Lanari, P. Daniele, F. Consoli, E. Agolini, K. Margiotti, I. Bottillo, I. Torrente, A. Bruselles, C. Fusilli, A. Ficcadenti, S. Bargiacchi, E. Trevisson, M. Forzan, S. Giustini, C. Leoni, G. Zampino, M. Digilio, B. Dallapiccola, M. Clementi, M. Tartaglia and A. De Luca (2015). "p.Arg1809Cys substitution in neurofibromin is associated with a distinctive NF1 phenotype without neurofibromas." Eur J Hum Genet. 23(8): 1068–1071.

Pitman, J. L., J. J. McGill, K. P. Keegan and R. Allada (2006). "A dynamic role for the mushroom bodies in promoting sleep in Drosophila." Nature 441(7094): 753–756.

Rojnueangnit, K., J. Xie, A. Gomes, A. Sharp, T. Callens, Y. Chen, Y. Liu, M. Cochran, M. A. Abbott, J. Atkin, D. Babovic-Vuksanovic, C. P. Barnett, M. Crenshaw, D. W. Bartholomew, L. Basel, G. Bellus, S. Ben-Shachar, M. G. Bialer, D. Bick, B. Blumberg, F. Cortes, K. L. David, A. Destree, A. Duat-Rodriguez, D. Earl, L. Escobar, M. Eswara, B. Ezquieta, I. M. Frayling, M. Frydman, K. Gardner, K. W. Gripp, C. Hernandez-Chico, K. Heyrman, J. Ibrahim, S. Janssens, B. A. Keena, I. Llano-Rivas, K. Leppig, M. McDonald, V. K. Misra, J. Mulbury, V. Narayanan, N. Orenstein, P. Galvin-Parton, H. Pedro, E. K. Pivnick, C. M. Powell, L. Randolph, S. Raskin, J. Rosell, K. Rubin, M. Seashore, C. P. Schaaf, A. Scheuerle, M. Schultz, E. Schorry, R. Schnur, E. Siqveland, A. Tkachuk, J. Tonsgard, M. Upadhyaya, I. C. Verma, S. Wallace, C. Williams, E. Zackai, J. Zonana, C. Lazaro, K. Claes, B. Korf, Y. Martin, E. Legius and L. Messiaen (2015). "High Incidence of Noonan Syndrome Features Including Short Stature and Pulmonic Stenosis in Patients carrying NF1 Missense Mutations Affecting p.Arg1809: Genotype-Phenotype Correlation." Hum Mutat 36(11): 1052–1063.

Rosato, E. and C. P. Kyriacou (2006). "Analysis of locomotor activity rhythms in Drosophila." Nat Protoc 1(2): 559–568.

Roussou, I. G., K. Papanikolopoulou, C. Savakis and E. M. C. Skoulakis (2019). "Drosophila Bruton’s Tyrosine Kinase Regulates Habituation Latency and Facilitation in Distinct Mushroom Body Neurons." J Neurosci 39(44): 8730–8743.

Shafer, O. T. and A. C. Keene (2021). "The Regulation of Drosophila Sleep." Curr Biol 31(1): R38–R49.

Shaw, P., C. Cirelli, R. Greenspan and G. Tononi (2000). "Correlates of sleep and waking in Drosophila melanogaster." Science 287(5459): 1834–1837.

Sherekar, M., S. W. Han, R. Ghirlando, S. Messing, M. Drew, D. Rabara, T. Waybright, P. Juneja, H. O’Neill, C. B. Stanley, D. Bhowmik, A. Ramanathan, S. Subramaniam, D. V. Nissley, W. Gillette, F. McCormick and D. Esposito (2020). "Biochemical and structural analyses reveal that the tumor suppressor neurofibromin (NF1) forms a high-affinity dimer." J Biol Chem 295(4): 1105–1119.

Silva, R. F. O., B. R. Pinho, N. M. Monteiro, M. M. Santos and J. M. A. Oliveira (2022). "Automated analysis of activity, sleep, and rhythmic behaviour in various animal species with the Rtivity software." Sci Rep 12(1): 4179.

Siqueiros-Sanchez, M., Y. Serur, C. A. McGhee, T. F. Smith and T. Green (2025). "Social Communication in Ras Pathway Disorders: A Comprehensive Review From Genetics to Behavior in Neurofibromatosis Type 1 and Noonan Syndrome." Biol Psychiatry 97(5): 461–498.

Sisobhan, S., C. Rosensweig, B. C. Lear and R. Allada (2022). "SleepMat: a new behavioral analysis software program for sleep and circadian rhythms." Sleep 45(12): zsac195.

The, I., G. E. Hannigan, G. S. Cowley, S. Reginald, Y. Zhong, J. F. Gusella, I. K. Hariharan and A. Bernards (1997). "Rescue of a Drosophila NF1 mutant phenotype by protein kinase A." Science 276(5313): 791–794.

Tong, J., F. Hannan, Y. Zhu, A. Bernards and Y. Zhong (2002). "Neurofibromin regulates G protein-stimulated adenylyl cyclase activity." Nat Neurosci 5(2): 95–96.

Vassallo, G., Z. Mughal, L. Robinson, D. Weisberg, S. Roberts, E. Hupton, J. Eelloo, E. Burkitt Wright, S. Garg, L. Lewis, D. Evans and S. Stivaros (2020). "Perceived fatigue in children and young adults with neurofibromatosis type 1." J Paediatr Child Health. 56(6): 878–883.

Walker, J. A., A. V. Tchoudakova, P. T. McKenney, S. Brill, D. Wu, G. S. Cowley, I. K. Hariharan and A. Bernards (2006). "Reduced growth of Drosophila neurofibromatosis 1 mutants reflects a non-cell-autonomous requirement for GTPase-Activating Protein activity in larval neurons." Genes Dev 20(23): 3311–3323.

Weiss, J. T. and J. M. Donlea (2021). "Roles for Sleep in Neural and Behavioral Plasticity: Reviewing Variation in the Consequences of Sleep Loss." Front Behav Neurosci 15: 777799.

Wu, L. and C. Liu (2023). "Integrated neural circuits of sleep and memory regulation in Drosophila." Curr Opin Insect Sci 59: 101105.

Wu, R., E. Legius, W. Robberecht, M. Dumoulin, J. J. Cassiman and J. P. Fryns (1996). "Neurofibromatosis type I gene mutation in a patient with features of LEOPARD syndrome." Hum Mutat 8(1): 51–56.

Young, L. C., R. Goldstein de Salazar, S. W. Han, Z. Y. S. Huang, A. Merk, M. Drew, J. Darling, V. Wall, R. Grisshammer, A. Cheng, M. R. Allison, M. J. Sale, D. V. Nissley, D. Esposito, J. Ognjenovic and F. McCormick (2023). "Destabilizing NF1 variants act in a dominant negative manner through neurofibromin dimerization." Proc Natl Acad Sci U S A 120(5): e2208960120.

Yurgel, M. E. and A. C. Keene (2018). "Sleep: Helicon Cells Charge the Circuit." Curr Biol 28(7): R317–r319.

Zhang, J., H. Tong, X. A. Fu, Y. Zhang, J. Liu, R. Cheng, J. Liang, J. Peng, Z. Sun, H. Liu, F. Zhang, W. Lu, M. Li and Z. Yao (2015). "Molecular Characterization of NF1 and Neurofibromatosis Type 1 Genotype-Phenotype Correlations in a Chinese Population." Scientific Reports 5(1): 11291.

