## Supplemental material for "Neurofibromin Regulates Sleep Consolidation via Cholinergic Neurons in a *Drosophila* Model of Neurofibromatosis 1"

### SUPPLEMENTAL FIGURES AND LEGENDS

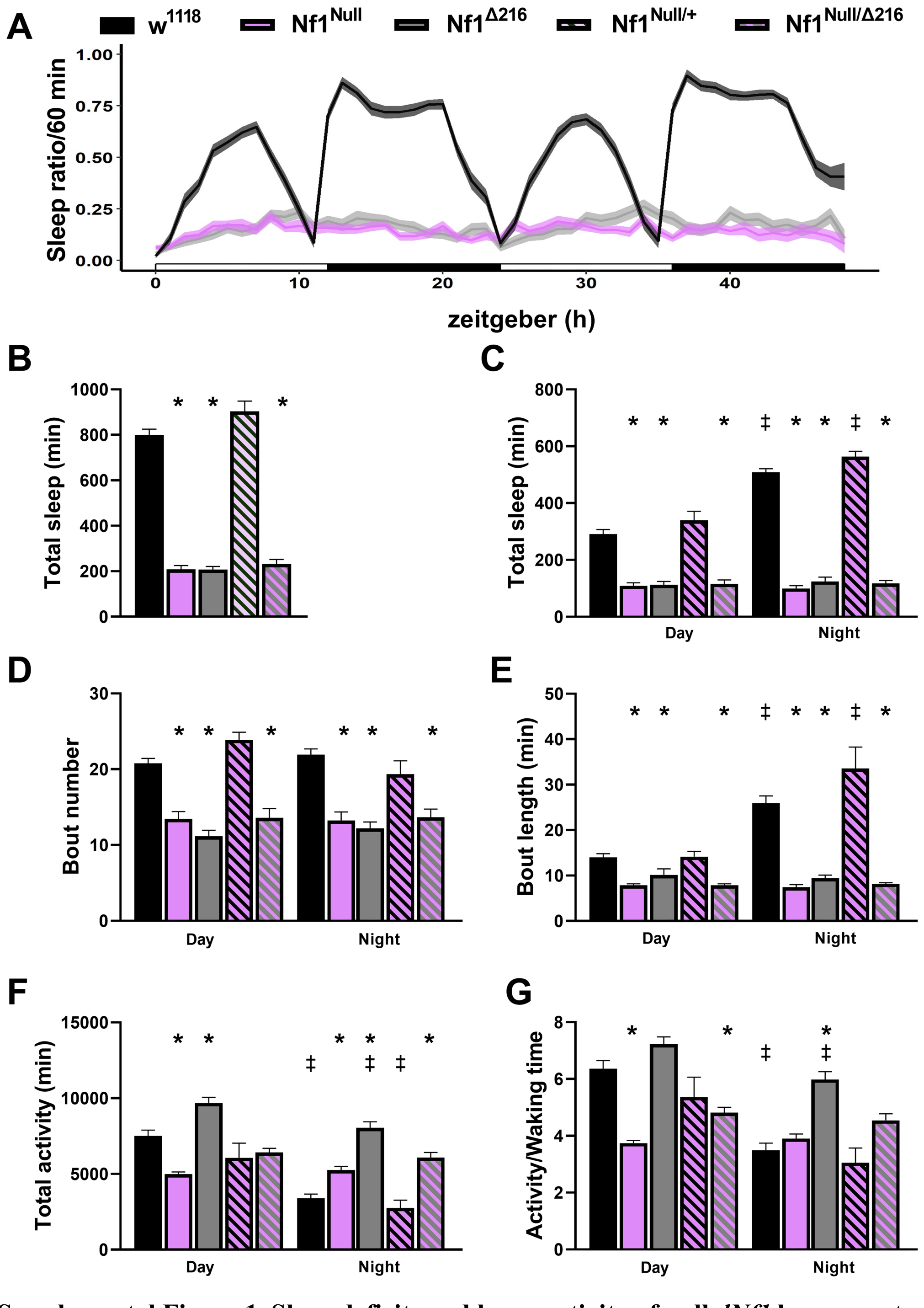

**Supplemental Figure 1. Sleep deficits and hyperactivity of null *dNf1* homozygotes and heteroallelics.**

**A.** An actogram quantifying the amount of time spent asleep per hour over a 48 hr period as indicated for $w^{1118}$ controls (black line, n= 57), $dNf1^{Null}$ homozygotes (magenta line, n= 37), and $dNf1^{\Delta 216}$ homozygotes (n= 54). The mean is indicated by the darker lines, and the standard error of the mean (SEM) is indicated by the lighter width of the lines. Daytime is denoted by the open bars and nighttime by the filled bars on the x axis.

**B-G.** Quantification of sleep and its attributes in $w^{1118}$ control animals (black bars), *dNf1*$^{Null}$ homozygotes (magenta bars), *dNf1*$^{\Delta216}$ homozygotes (grey bars), *dNf1*$^{Null}$ heterozygotes (black-striped magenta bars, n=11) and *dNf1*$^{Null/\Delta216}$ heteroallelics (n=37). Multiple non-parametric comparisons (Shapiro-Wilk a<0.05) were performed with Kruskal-Wallis tests, while two-way comparisons were performed with Mann-Whitney tests. Multiple comparisons of normally distributed data were performed with Dunnett's test following initial ANOVA. Stars indicate that the mutant score is significantly different than that of controls, the double dagger signifies differences in controls or mutants in the day relative to night-time scores and # denotes significant differences of the experimental score compared to that of the controls. All statistical comparisons are detailed in Suppl. Table 1.

**B**. The total time spent asleep over the monitored period was significantly less (p<0.0001 for all) for *dNf1*$^{Null}$ homozygotes, *dNf1*$^{\Delta216}$ homozygotes and *dNf1*$^{Null/\Delta216}$ heteroallelics relative to $w^{1118}$ controls indicated by the asterisks. In contrast, the time *dNf1*$^{Null}$ heterozygotes spent asleep was not different than that of controls (p=0.9999).

**C.** The total time spent asleep during the day and at night revealed that $w^{1118}$ control animals and *dNf1*$^{Null}$ heterozygotes slept significantly more at night than during the day (p<0.0001, double daggers). In contrast, *dNf1*$^{Null}$ homozygotes, *dNf1*$^{\Delta216}$ homozygotes and *dNf1*$^{Null/\Delta216}$ heteroallelics slept significantly less than controls both during the day (p<0.0001 for all comparisons) and at night (p<0.0001 for all comparisons), denoted by the asterisks, and their sleep at night was not increased relative to daytime.

**D.** The number of sleep episodes was significantly reduced in *dNf1*$^{Null}$ homozygotes, *dNf1*$^{\Delta216}$ homozygotes and *dNf1*$^{Null/\Delta216}$ heteroallelics compared to those of $w^{1118}$ controls both during the day and at night (p<0.0001 for all comparisons), denoted by the asterisks, while that of *dNf1*$^{Null}$ heterozygotes remained similar to controls (p= 0.1086 or day and p=0.2193 for night-time).

**E.** Sleep bout length was increased at night for control animals (p<0.0001, double daggers), but not *dNf1*$^{Null}$ homozygotes, *dNf1*$^{\Delta216}$ homozygotes (p=0.7544) and *dNf1*$^{Null/\Delta216}$ heteroallelics, which presented significantly reduced bout length (p<0.0001) relative to controls (asterisks). Again, bout length in *dNf1*$^{Null}$ heterozygotes did not differ from that of controls (p=0.9999 for day and night).

**F.** Total activity decreased at night relative to daytime for controls (p<0.0001) and *dNf1*$^{Null}$ heterozygotes, as indicated by the double daggers. However, activities of *dNf1*$^{Null}$ homozygotes (p<0.0001) and *dNf1*$^{\Delta216}$ homozygotes (p=0.0016) were different during the day relative to controls, while surprisingly those of *dNf1*$^{Null/\Delta216}$ heteroallelics were not (p=0.1823). In contrast, at nighttime, the activities of *dNf1*$^{Null}$ homozygotes (p=0.003), *dNf1*$^{\Delta216}$ homozygotes and *dNf1*$^{Null/\Delta216}$ heteroallelics (p<0.0001 for both) were significantly different than that of controls, while that of *dNf1*$^{Null}$ heterozygotes was not (p=0.9999).

**G.** Activity between sleep episodes decreased at night relative to daytime for controls and for the *dNf1*$^{Null}$ heterozygotes ($p<0.0001$ for both), as indicated by the double daggers. However, the waking time activity during the day for *dNf1*$^{Null}$ homozygotes and *dNf1*$^{Null/\Delta 216}$ heteroallelics relative to controls was significantly reduced ($p<0.0001$ and $p=0.0021$, respectively), but surprisingly, that of *dNf1*$^{\Delta 216}$ homozygotes was not ($p=0.1603$). Interestingly, at night-time only the activity of *dNf1*$^{\Delta 216}$ homozygotes was significantly elevated relative to controls ($p<0.0001$).

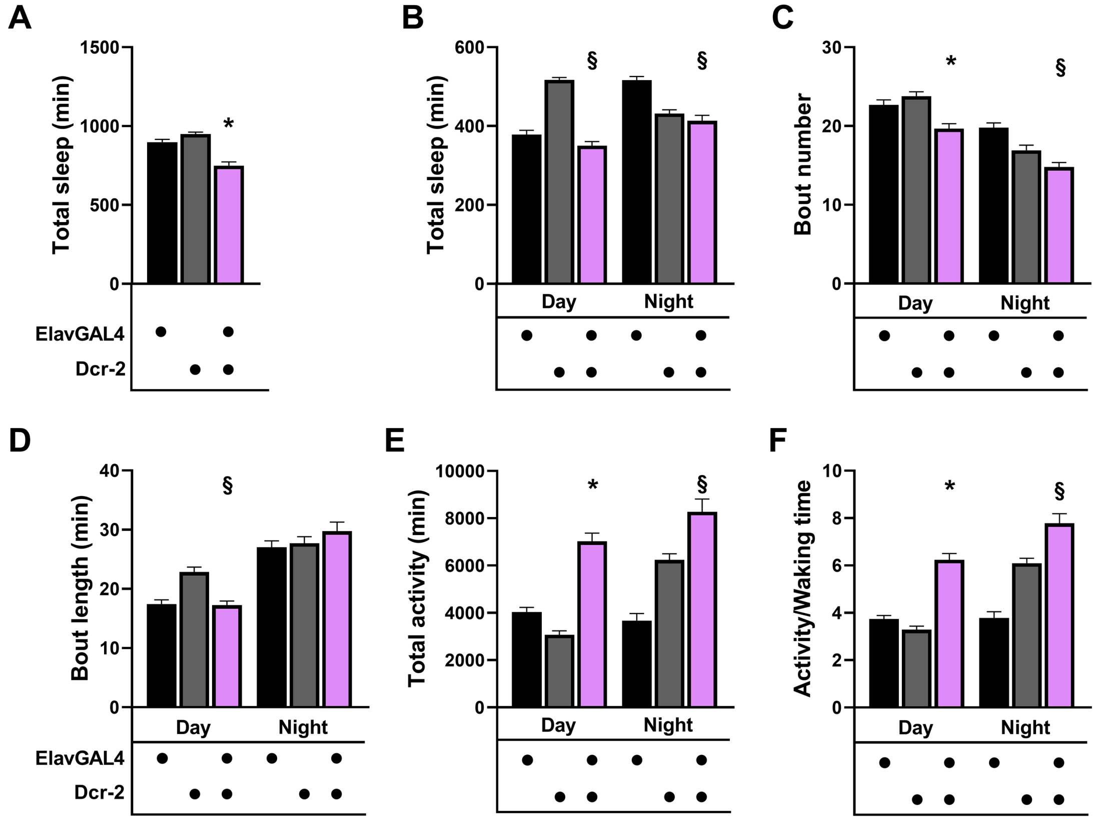

**Supplemental Figure 2. Pan-neuronal expression of Dcr-2 results in hyperactivity, while sleep architecture undergoes minor changes but remains largely intact.**

Quantification of sleep and its attributes in ElavGal4/+ control animals (black bars), flies carrying, but not expressing, the Dcr-2 transgene (gray bars), and those expressing Dcr-2 pan-neuronally (magenta bars). Multiple non-parametric (as determined by Shapiro-Wilk, a<0.05) comparisons were performed with Kruskal-Wallis tests, while two-way comparisons were performed with Mann-Whitney tests. Multiple comparisons of normally distributed data were performed with Dunnett's test following initial ANOVA. Stars indicate that the mutant score is significantly different than that of controls, while § indicates differences from one , but the other controlAll statistical comparisons are detailed in Suppl. Table 1.

**A**. The total time spent asleep over the monitored period was significantly less for ElavGal4>Dcr-2-expressing animals, relative to that of ElavGal4/+ and Dcr-2/+ (ANOVA p<0.0001, Dunnett's p<0.0001 from both controls)

**B.** The total time spent asleep during the day and at night revealed the reason total sleep was reduced for ElavGal4>Dcr-2 flies, as their daytime sleep was significantly different from Dcr-2/+ (p<0.0001, section sign), but not from the other control ElavGal4/+ (p=0.2316). Similarly, nighttime sleep for ElavGal4>Dcr-2-expressing flies was significantly different from that of ElavGal4/+ (p<0.0001, section sign), but not from Dcr-2/+ flies (p=0.9999).

**C.** The number of daytime sleep episodes were significantly reduced (asterisk) in ElavGal4>Dcr-2-expressing animals compared to those of ElavGal4/+ (p=0.0069) and Dcr-2/+ (p<0.0001). However, at nighttime was significantly different from that of ElavGal4/+ (p<0.0001, section sign), but not from Dcr-2/+ (p=0.0461).

**D.** Daytime sleep bout length of ElavGal4>Dcr-2 flies was significantly different from that of Dcr-2/+ (p<0.0001 section sign), but not from ElavGal4/+ (p=0.9999). In contrast, at night, sleep bout length was not significantly different from both controls (p=0.3159 and 0.9023 respectively).

**E.** Total activity was highly elevated for ElavGal4>Dcr-2 relative to both controls during the day (p<0.0001, asterisk), but not at nighttime, where ElavGal4>Dcr-2 activity was significantly different from that of ElavGal4/+ (p<0.0001, section sign), but not from Dcr-2/+ (p=0.9999).

**F.** Activity when awake was significantly elevated for ElavGal4>Dcr-2 relative to both controls during the day p<0.0001, asterisk), but not at nighttime, where ElavGal4>Dcr-2 activity was significantly different from that of ElavGal4/+ (p<0.0001, section sign), but not from Dcr-2/+ (p=0.0402).

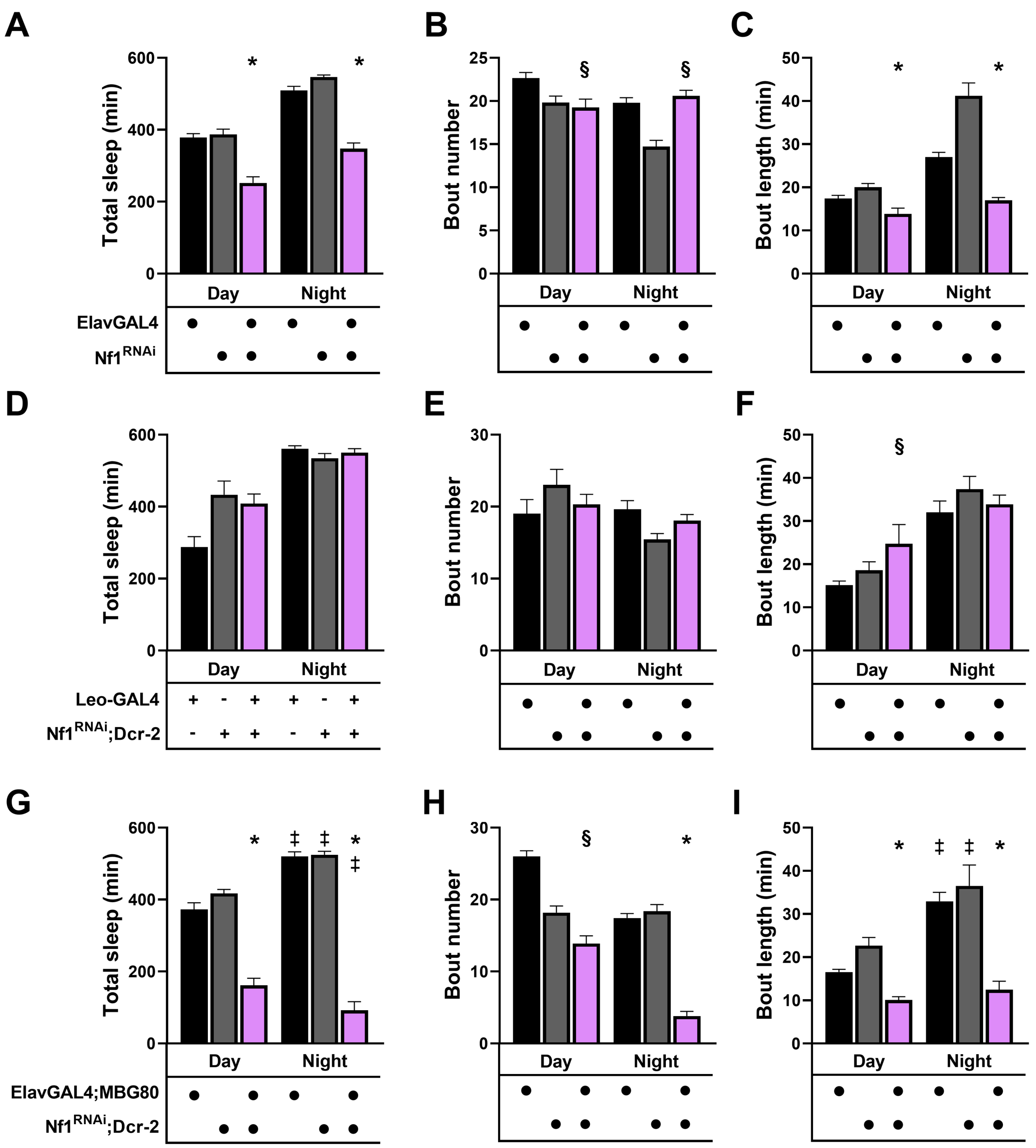

**Supplemental Figure 3. Targeted abrogation of dNf1 in distinct neuronal types**

Quantification of sleep and its attributes in animals where dNf1 is abrogated, as indicated below by the tissue-specific drivers (magenta bars) and the respective controls driver/+ (black bars) and *dNf1*$^{RNAi}$/+;Dcr-2/+ (grey bars). Multiple non-parametric (as determined by Shapiro-Wilk, a<0.05) comparisons were performed with Kruskal-Wallis tests, while two-way comparisons were performed with Mann-Whitney. Stars indicate that the mutant score is significantly different than that of controls, the double dagger signifies differences in controls or mutants in the day relative to night-time scores, while § indicates differences from one , but the other control. All statistical comparisons are detailed in Suppl. Table 1.

**A-C.** Selective pan-neuronal dNf1 knockdown in the absence of Dcr2.

**A.** Pan-neuronal attenuation of dNf1 precipitated significant ($p<0.0001$) deficits in total sleep during the day and at night.

**B.** Pan-neuronal attenuation of dNf1 significantly reduced sleep bout number relative to Elav/+ controls ($p=0.0086$), but not $Nf1^{RNAi}$/+ ($p=0.9999$) during the day, but the opposite at night ($p<0.0001$ *vs* $Nf1^{RNAi}$/+, but $p=0.6199$ *vs* Elav/+).

**C.** In contrast, pan-neuronal attenuation of dNf1significantly reduced sleep bout duration ($p<0.0001$ *vs* $Nf1^{RNAi}$/+, $p=0.0002$ *vs* Elav/+) during the day and similarly at night-time ($p<0.0001$ for both controls).

**D-F**. Selective dNf1 abrogation in the mushroom bodies. No significant effects were revealed except for daytime bout length ($p=0.0051$).

**G-I.** Selective pan-neuronal dNf1 knockdown sparing the mushroom bodies.

**G.** Total daytime and night-time sleep were significantly ($p<0.0001$ for both) reduced upon pan-neuronal dNf1 loss sparing the mushroom bodies.

**H.** Sleep bout number was significantly reduced ($p<0.0001$) relative to ElavGal4;MBG80/+, but not the other control ($p=0.0369$) during the day, but it was against both controls at night-time ($p<0.0001$).

**I**. Similarly, sleep bout length was significantly reduced during the day and night ($p<0.0001$).

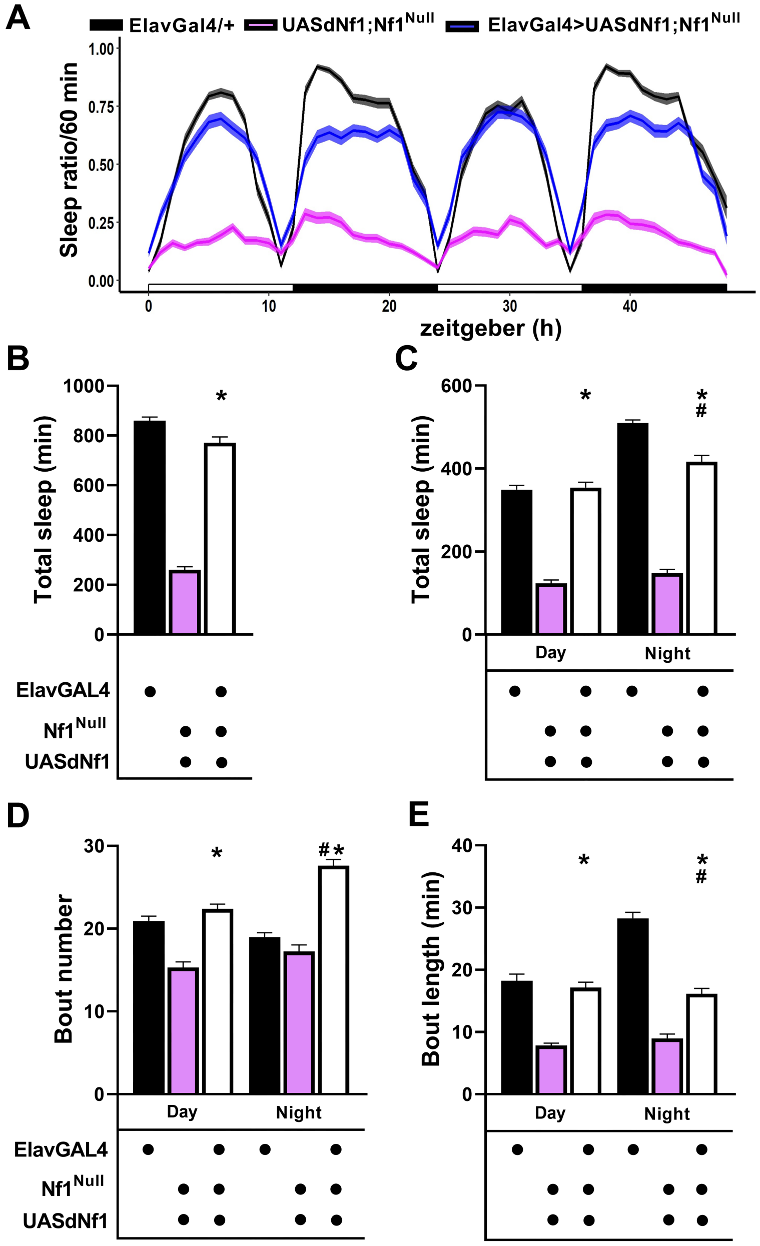

**Supplemental Figure 4. Pan-neuronal restoration of Neurofibromin in the null mutant ameliorates its sleep deficits**

**A.** An actogram quantifying the amount of time spend asleep per hour over a 48 hr period as indicated for ElavGal4/+ controls (black line) and *UASdNf1*/+;*dNf1*$^{Null}$ homozygotes (magenta line) and ElavGal4>*UASdNf1*/+;*dNf1*$^{Null}$ (blue line). The mean is indicated by the darker lines and the standard error of the mean (SEM) is indicated by the lighter width of the lines. Daytime is denoted by the open bars and nighttime by the filled bars on the x axis.

**B-E.** Quantification of sleep and its attributes in ElavGal4/+ controls (black bars) and *UASdNf1*/+;*dNf1*$^{Null}$ homozygotes (magenta bars) and ElavGal4>*UASdNf1*/+;*dNf1*$^{Null}$ flies (open bars). Stars indicate that the mutant score is significantly different than that of controls, the double dagger signifies differences in controls or mutants in the day relative to night-time scores and # denotes significant differences of the experimental score compared to that of the controls. As all data were not normally distributed (as determined by Shapiro-Wilk, a<0.05), statistical comparisons were performed with Mann-Whitney tests with statistical details presented in Suppl. Table 1.

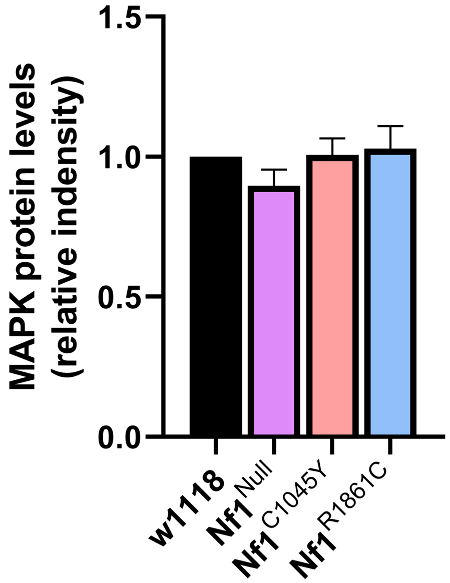

**Supplemental Figure 5. MAPK levels are not changed in the mutants.**

The abundance of MAPK tryptic peptides in head lysates of *w*$^{1118}$ control animals (dark bars) was arbitrarily set to 1 and the relative levels of MAPK based on the normalized spectral intensity in *dNf1*$^{E2}$ homozygotes (magenta bar), *dNf1*$^{C1045Y}$ (light red bar) and *dNf1*$^{R1861C}$ (blue bar) homozygotes are shown. The difference was not significant (ANOVA F= 1.341, p=0.2892, n=6)

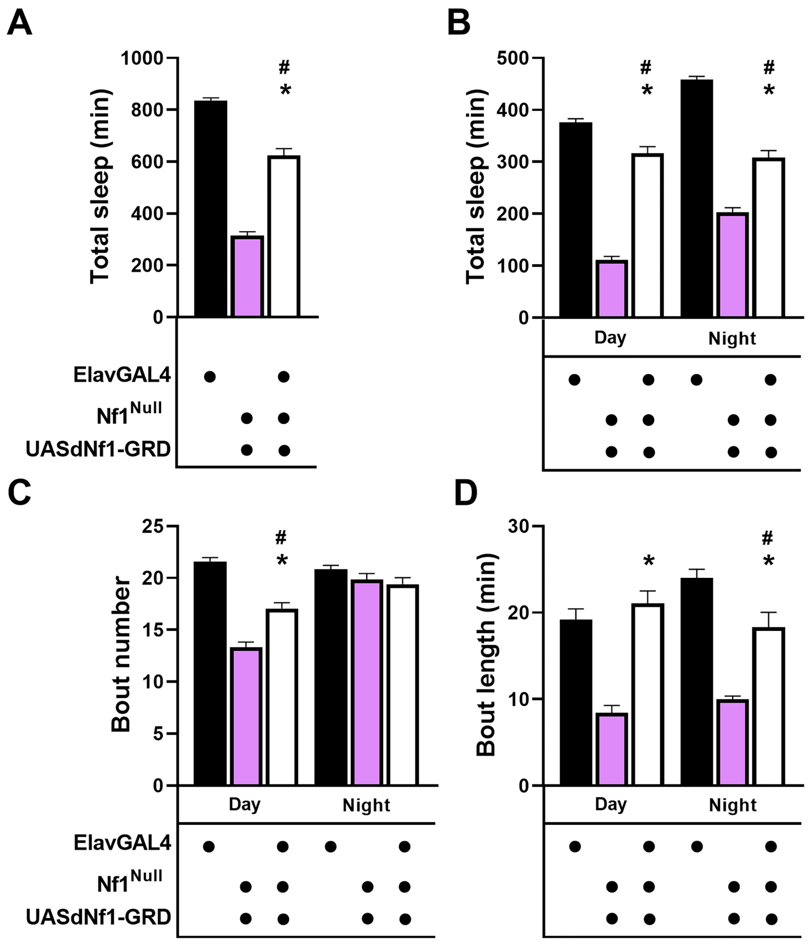

**Supplemental Figure 6. Pan-neuronal expression of the GRD transgene ameliorates the sleep phenotypes of the null mutant.**

Quantification of sleep and its attributes in ElavGal4/+ controls (black bars) and *UASdNf1-GRD*/+;*dNf1*$^{Null}$ homozygotes (magenta bars) and ElavGal4> *UASdNf1-GRD*/+;*dNf1*$^{Null}$ flies (open bars). Stars indicate that the mutant score is significantly different than that of controls and # denotes significant differences of the experimental score compared to that of the controls. Statistical details presented in Suppl. Table 1.

**A.** Pan-neuronal expression of the GRD transgene ameliorated the sleep deficit of the mutants ($p<0.0001$), which however did not attain control levels.

**B.** Amelioration of the sleep deficit was evenly parsed during the day and at night-time ($p<0.0001$ for both), but did not attain control sleep levels at both periods.

**C.** Sleep bout number was increased during the day upon GRD transgene expression relative to the levels presented by the mutant ($p<0.0001$), but nonetheless did not attain the sleep bout level of control flies.

**D.** Sleep bout length was significantly increased ($p<0.0001$) upon GRD transgene expression during the day and at night and reached control levels during the day, but at night-time.

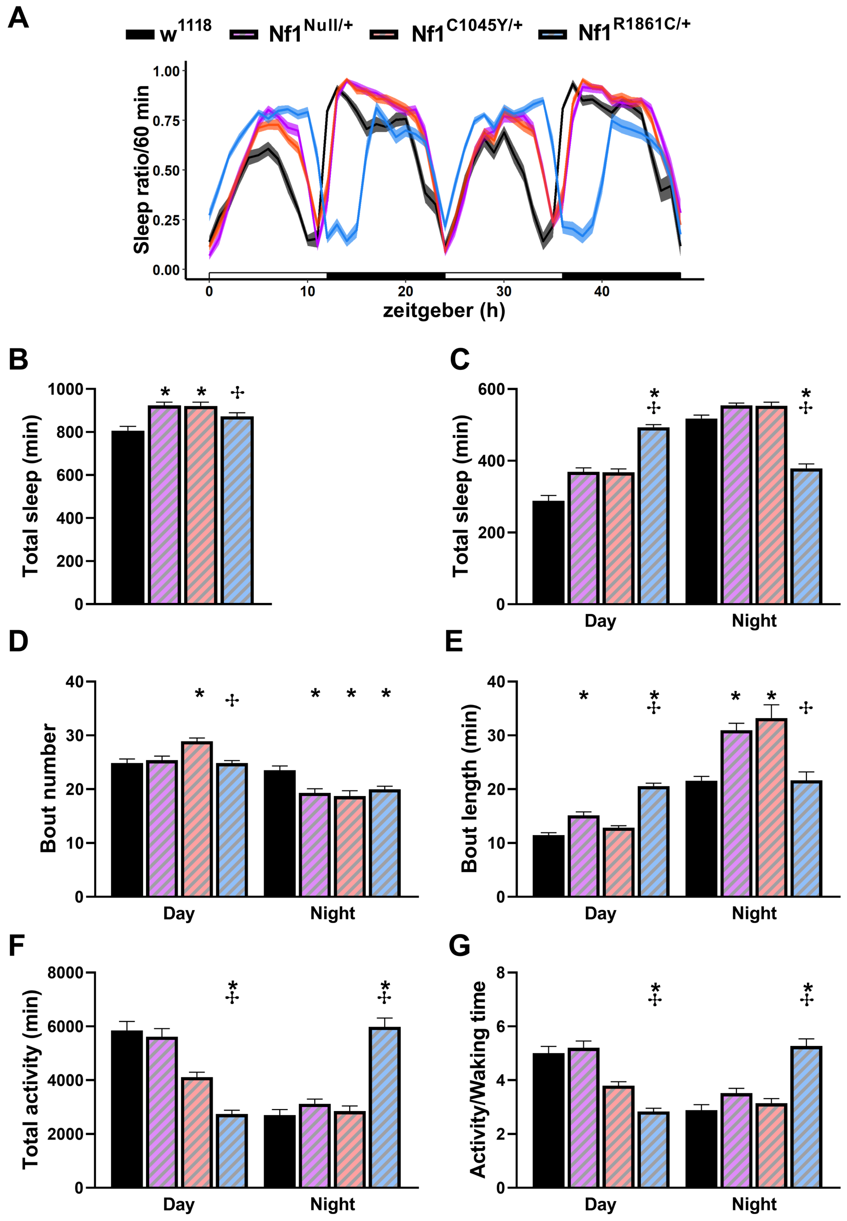

**Supplemental Figure 7. Sleep deficits and hyperactivity of null *dNf1* heterozygotes.**

Stars indicate that the mutant score is significantly different than that of controls, the cross signifies differences among mutants. Statistical details in Suppl Table 1.

**A.** Average sleep length plotted in 60-minute intervals over a 48-hour period under a 12-hour light / 12-hour dark (LD) cycle. The horizontal bar underneath indicates light (white) and dark (black) phases. Zeitgeber time (ZT, in hours) is shown on the x-axis against the sleep ratio on the y-axis for control w1118 (black line), $Nf1^{Null/+}$ (magenta line), $Nf1^{C1045Y/+}$ (pink line), and $Nf1^{R1861C/+}$ (blue line) flies.

**B.** Quantification of total sleep duration averaged over a 24-hour cycle across the indicated genotypes. Both *dNf1*$^{Null/+}$ and *dNf1*$^{C1045Y/+}$ animals showed significantly increased total sleep compared with *w*$^{1118}$ controls ($p<0.0001$). Total sleep in *dNf1*$^{R1861C/+}$ was also altered ($p=0.0073$) compared to the *dNf1*$^{C1045Y/+}$, as indicated by the corresponding statistical comparison (cross).

**C.** Total sleep duration separated into daytime (light phase) and nighttime (dark phase) conditions. During the daytime, *dNf1*$^{R1861C/+}$ flies showed significantly increased ($p<0.0001$) total sleep compared with controls. During the night, *dNf1*$^{R1861C/+}$ showed reduced ($p<0.0001$) sleep compared with controls.

**D.** The average number of discrete sleep episodes (bouts) quantified during the day and night. During the daytime, *dNf1*$^{R1861C/+}$ flies displayed a significantly increased number of sleep bouts compared with controls ($p=0.0004$), consistent with increased sleep fragmentation. During the nighttime, sleep bout number was significantly reduced in *dNf1*$^{Null/+}$, *dNf1*$^{C1045Y/+}$ and *dNf1*$^{R1861C/+}$ relative to the control group ($p=0.0006$, 0.0008 and 0.0041 respectively).

**E.** Mean sleep bout length during the day and night showing significant alterations in sleep consolidation across the different Nf1 genotypes. During the day, *dNf1*$^{Null/+}$and *dNf1*$^{R1861C/+}$ flies exhibited significantly longer sleep bouts compared with controls ($p=0.003$ and $p<0.00001$ respectively), whereas Nf1$^{C1045Y/+}$ showed a comparatively shorter bout duration. During the nighttime, sleep bout length was increased in *dNf1*$^{Null/+}$ and *dNf1*$^{C1045Y/}$ animals relative to controls, while the *dNf1*$^{R1861C/+}$ showed a smaller increase.

**F.** Total locomotor activity counts measured during the diurnal and nocturnal periods. Total locomotor activity during the day and nighttime in controls, *dNf1*$^{Null/+}$, *dNf1*$^{C1045Y/+}$ and *dNf1*$^{R1861C/+}$ flies. During the daytime, *dNf1*$^{C1045Y/+}$ ($p=0.0126$) and particularly *dNf1*$^{R1861C/+}$ ($p<0.0001$) flies showed reduced total activity relative to controls. In contrast, nighttime activity was markedly increased in *dNf1*$^{R1861C/+}$ flies ($p<0.0001$) compared with the other genotypes, indicating altered locomotor activity across the diurnal cycle.

**G.** Waking Activity Index: Total activity normalized over total waking time (Activity/Waking time) calculated for the day and night phases. The performance of *dNf1*$^{R1861C/+}$ flies was significantly different than that of controls ($p<0.0001$) and from the other mutant heterozygotes both at day and night-time.

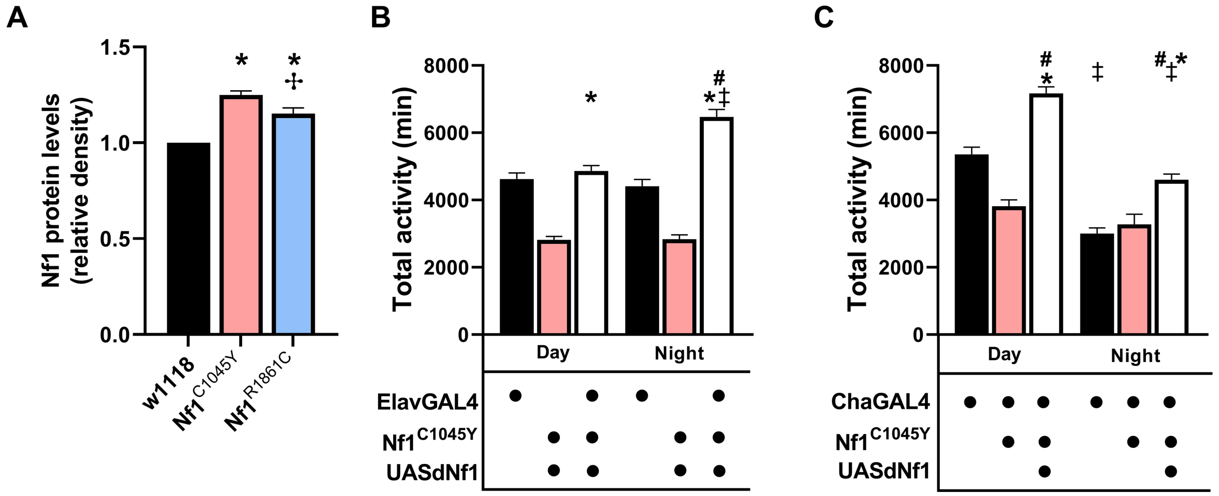

**Supplemental Figure 8. Amelioration of activity deficits of *dNf1*$^{C1045Y}$ homozygotes by pan-neuronal and cholinergic neuron-specific restoration of dNF1.**

Stars indicate that the mutant score is significantly different than that of controls and # denotes significant differences of the experimental score compared to that of the controls, while § indicates differences from one, but the other control. The crosses indicate differences among the mutants Statistical details presented in Suppl. Table 1.

**A.** The abundance of Neurofibromin-derived tryptic peptides in head lysates of $w^{1118}$ control animals (dark bars) was arbitrarily set to 1 and the relative levels in *dNf1*$^{C1045Y}$ (light red bar) and *dNf1*$^{R1861C}$ (blue bar) homozygotes are shown. ANOVA F= 38.9, p<0.0001, n=6. Unpaired t-tests with Welch's correlation indicated that the level of dNF1 is significantly different in *dNf1*$^{C1045Y}$ (p<0.0001) and *dNf1*$^{R1861C}$ (p=0.0039) relative to that of controls. (n=6).

**B.** Pan-neuronal expression of the wild type transgene rescued the day and nighttime hypoactivity of Nf1$^{C1045Y}$ homozygotes.

**C.** The reduced daytime activity of dNf1$^{C1045Y}$ homozygotes was also elevated by dNf1 transgene expression in cholinergic neurons.

**SUPPLEMENTAL TABLE 1**

| Genotype | Mean ± SEM | n | p |
|---|---|---|---|
| **Fig. 1A** Mann-Whitney (U)= 0 **p=0.0022** | | | |
| $w^{1118}$ | $1.4e^6$ ± 28070 | 6 | |
| $Nf1^{Null}$ | $1.3e^5$ ± 12485 | 6 | **0.0022** |
| **Fig. 1C** Mann-Whitney (U)= 3911 **p<0.0001** | | | |
| $w^{1118}$ | 668.3 ± 28 | 156 | |
| $Nf1^{Null}$ | 274.6 ± 24 | 127 | **<0.0001** |
| **Fig. 1D** Mann-Whitney (U)= 4722 **p<0.0001** | | | |
| $w^{1118}$ (Day) | 263.9 ± 13 | 156 | |
| $Nf1^{Null}$ (Day) | 123.0 ± 11 | 127 | **<0.0001** |
| Mann-Whitney (U)= 3644 **p<0.0001** | | | |
| $w^{1118}$ (Night) | 404.3 ± 16 | 156 | |
| $Nf1^{Null}$ (Night) | 151.5 ± 13 | 127 | **<0.0001** |
| Mann-Whitney (U)= 6591 **p<0.0001** | | | |
| $w^{1118}$ (Day) | 263.9 ± 13 | 156 | |
| $w^{1118}$ (Night) | 404.3 ± 16 | 156 | **<0.0001** |
| Mann-Whitney (U)= 7170 p=0.1267 | | | |
| $Nf1^{Null}$ (Day) | 123.0 ± 11 | 127 | |
| $Nf1^{Null}$ (Night) | 151.5 ± 13 | 127 | 0.1267 |
| **Fig. 1E** Mann-Whitney (U)=6381 **p<0.0001** | | | |
| $w^{1118}$ (Day) | 19.22 ± 0.73 | 156 | |
| $Nf1^{Null}$ (Day) | 12.90 ± 0.87 | 127 | **<0.0001** |
| Mann-Whitney (U)=7250 **p<0.0001** | | | |
| $w^{1118}$ (Night) | 21.16 ± 0.87 | 156 | |
| $Nf1^{Null}$ (Night) | 15.57 ± 1.1 | 127 | **<0.0001** |
| Mann-Whitney (U)=10413 p=0.0276 | | | |
| $w^{1118}$ (Day) | 19.22 ± 0.73 | 156 | |
| $w^{1118}$ (Night) | 21.16 ± 0.87 | 156 | 0.0276 |
| Mann-Whitney (U)=7206 p=0.1427 | | | |
| $Nf1^{Null}$ (Day) | 12.90 ± 0.87 | 127 | |
| $Nf1^{Null}$ (Night) | 15.57 ± 1.1 | 127 | 0.1427 |
| **Fig. 1F** Mann-Whitney (U)= 4427 **p<0.0001** | | | |
| $w^{1118}$ (Day) | 11.69 ± 0.47 | 150 | |
| $Nf1^{Null}$ (Day) | 8.110 ± 0.40 | 126 | **<0.0001** |
| Mann-Whitney (U)= 3237 **p<0.0001** | | | |
| $w^{1118}$ (Night) | 16.77 ± 0.68 | 149 | |
| $Nf1^{Null}$ (Night) | 8.420 ± 0.42 | 126 | **<0.0001** |
| Mann-Whitney (U)= 6566 **p<0.0001** | | | |

| | | | |
|---|---|---|---|
| $w^{1118}$ (Day) | 11.69 ± 0.47 | 150 | |
| $w^{1118}$ (Night) | 16.77 ± 0.68 | 149 | **<0.0001** |
| Mann-Whitney (U)= 6898 p=0.0724 | | | |
| $Nf1^{Null}$ (Day) | 8.110 ± 0.40 | 126 | |
| $Nf1^{Null}$ (Night) | 8.420 ± 0.42 | 126 | 0.0724 |
| **Fig. 1G** Mann-Whitney (U)= 8807 p=0.1085 | | | |
| $w^{1118}$ (Day) | 5443 ± 220 | 156 | |
| $Nf1^{Null}$ (Day) | 5712 ± 150 | 127 | 0.1085 |
| Mann-Whitney (U)= 5333 **p<0.0001** | | | |
| $w^{1118}$ (Night) | 3379 ± 180 | 156 | |
| $Nf1^{Null}$ (Night) | 5068 ± 170 | 127 | **<0.0001** |
| Mann-Whitney (U)= 6588 **p<0.0001** | | | |
| $w^{1118}$ (Day) | 5443 ± 220 | 156 | |
| $w^{1118}$ (Night) | 3379 ± 180 | 156 | **<0.0001** |
| Mann-Whitney (U)= 6431 **p=0.0053** | | | |
| $Nf1^{Null}$ (Day) | 5712 ± 150 | 127 | |
| $Nf1^{Null}$ (Night) | 5068 ± 170 | 127 | **0.0053** |
| **Fig. 1H** Mann-Whitney (U)= 9521 p=0.5744 | | | |
| $w^{1118}$ (Day) | 4.492 ± 0.17 | 156 | |
| $Nf1^{Null}$ (Day) | 4.272 ± 0.089 | 127 | 0.5744 |
| Mann-Whitney (U)= 6623 **p<0.0001** | | | |
| $w^{1118}$ (Night) | 3.050 ± 0.13 | 156 | |
| $Nf1^{Null}$ (Night) | 3.839 ± 0.10 | 127 | **<0.0001** |
| Mann-Whitney (U)= 6754 **p<0.0001** | | | |
| $w^{1118}$ (Day) | 4.492 ± 0.17 | 156 | |
| $w^{1118}$ (Night) | 3.050 ± 0.13 | 156 | **<0.0001** |
| Mann-Whitney (U)= 6189 **p=0.0014** | | | |
| $Nf1^{Null}$ (Day) | 4.272 ± 0.089 | 127 | |
| $Nf1^{Null}$ (Night) | 3.839 ± 0.10 | 127 | **0.0014** |
| **Fig. 2B** Kruskal Wallis$_{(4)}$ = 41.48 **p<0.0001** | | | |
| ElavGal4/+ | 233192 ± 510 | 10 | |
| ElavGal4>Dcr-2 | 231490 ± 510 | 10 | >0.9999 |
| $Nf1^{Null}$ | 0 | 10 | **<0.0001** |
| ElavGal4>Nf1RNAi; Dcr-2 | 0 | 10 | **<0.0001** |
| **Fig. 2C** Kruskal Wallis$_{(2)}$ = 82.15 **p<0.0001** | | | |

| | | | |
|---|---|---|---|
| ElavGal4>Nf1RNAi; Dcr-2 | 161.5 ± 37 | 32 | |
| ElavGal4/+ | 887.6 ± 20 | 64 | **<0.0001** |
| Nf1RNAi; Dcr-2/+ | 1030 ± 20 | 32 | **<0.0001** |
| **Fig. 2D** Kruskal Wallis$_{(2)}$ = 86.77 **p<0.0001** | | | |
| ElavGal4>Nf1RNAi; Dcr-2 (Day) | 89.02 ± 16 | 32 | |
| ElavGal4/+ (Day) | 378.3 ± 11 | 64 | **<0.0001** |
| Nf1RNAi; Dcr-2/+ (Day) | 492.6 ± 12 | 32 | **<0.0001** |
| Kruskal Wallis$_{(2)}$ = 70.89 **p<0.0001** | | | |
| ElavGal4>Nf1RNAi; Dcr-2 (Night) | 72.34 ± 23 | 32 | |
| ElavGal4/+ (Night) | 509.2 ± 12 | 64 | **<0.0001** |
| Nf1RNAi; Dcr-2/+ (Night) | 536.9 ± 11 | 32 | **<0.0001** |
| Mann-Whitney (U)= 509 **p<0.0001** | | | |
| ElavGal4/+ (Day) | 378.3 ± 11 | 64 | |
| ElavGal4/+ (Night) | 509.2 ± 12 | 64 | **<0.0001** |
| Mann-Whitney (U)= 309.5 **p=0.0060** | | | |
| Nf1RNAi; Dcr-2/+ (Day) | 492.6 ± 12 | 32 | |
| Nf1RNAi; Dcr-2/+ (Night) | 536.9 ± 11 | 32 | **0.0060** |
| Mann-Whitney (U)= 250.5 **p=0.0003** | | | |
| ElavGal4>Nf1RNAi; Dcr-2 (Day) | 89.02 ± 16 | 32 | |
| ElavGal4>Nf1RNAi; Dcr-2 (Night) | 72.34 ± 23 | 32 | **0.0003** |
| **Fig. 2E** Kruskal Wallis$_{(2)}$ = 44.01 **p<0.0001** | | | |
| ElavGal4>Nf1RNAi; Dcr-2 (Day) | 10.02 ± 1.3 | 32 | |
| ElavGal4/+ (Day) | 22.66 ± 0.65 | 64 | **<0.0001** |
| Nf1RNAi; Dcr-2/+ (Day) | 22.72 ± 0.92 | 32 | **<0.0001** |
| Kruskal Wallis$_{(2)}$ = 41.36 **p<0.0001** | | | |
| ElavGal4>Nf1RNAi; Dcr-2 (Night) | 5.953 ± 1.6 | 32 | |
| ElavGal4/+ (Night) | 19.78 ± 0.61 | 64 | **<0.0001** |
| Nf1RNAi; Dcr-2/+ (Night) | 17.25 ± 0.86 | 32 | **0.0001** |
| **Fig. 2F** Kruskal Wallis$_{(2)}$ = 75.49 **p<0.0001** | | | |
| ElavGal4>Nf1RNAi; Dcr-2 (Day) | 7.891 ± 0.40 | 32 | |
| ElavGal4/+ (Day) | 17.44 ± 0.71 | 64 | **<0.0001** |
| Nf1RNAi; Dcr-2/+ (Day) | 23.01 ± 1.2 | 32 | **<0.0001** |
| Kruskal Wallis$_{(2)}$ = 66.21 **p<0.0001** | | | |
| ElavGal4>Nf1RNAi; Dcr-2 (Night) | 8.349 ± 0.72 | 32 | |
| ElavGal4/+ (Night) | 27.04 ± 1.1 | 64 | **<0.0001** |
| Nf1RNAi; Dcr-2/+ (Night) | 34.35 ± 2.3 | 32 | **<0.0001** |
| Mann-Whitney (U)= 617 **p<0.0001** | | | |
| ElavGal4/+ (Day) | 17.44 ± 0.71 | 64 | |

| | | | |
|---|---|---|---|
| ElavGal4/+ (Night) | 27.04 ± 1.1 | 64 | **<0.0001** |
| Mann-Whitney (U)= 211 **p<0.0001** | | | |
| Nf1RNAi; Dcr-2/+ (Day) | 23.01 ± 1.2 | 32 | |
| Nf1RNAi; Dcr-2/+ (Night) | 34.35 ± 2.3 | 32 | **<0.0001** |
| Mann-Whitney (U)= 363.5 p=0.3014 | | | |
| ElavGal4>Nf1RNAi; Dcr-2 (Day) | 7.891 ± 0.40 | 32 | |
| ElavGal4>Nf1RNAi; Dcr-2 (Night) | 8.349 ± 0.72 | 32 | 0.3014 |
| **Fig. 2G** Kruskal Wallis $H_{(2)}$ = 71.48 **p<0.0001** | | | |
| ElavGal4>Nf1RNAi; Dcr-2 (Day) | 7233 ± 330 | 32 | |
| ElavGal4/+ (Day) | 3954 ± 170 | 63 | **<0.0001** |
| Nf1RNAi; Dcr-2/+ (Day) | 2440 ± 170 | 32 | **<0.0001** |
| Kruskal Wallis $H_{(2)}$ = 60.01 **p<0.0001** | | | |
| ElavGal4>Nf1RNAi; Dcr-2 (Night) | 10062 ± 550 | 32 | |
| ElavGal4/+ (Night) | 3535 ± 260 | 63 | **<0.0001** |
| Nf1RNAi; Dcr-2/+ (Night) | 3776 ± 260 | 32 | **<0.0001** |
| **Fig. 2H** ANOVA $F_{(2, 125)}$ = 48.75 **p<0.0001** | | | |
| ElavGal4>Nf1RNAi; Dcr-2 (Day) | 5.307 ± 0.21 | 32 | |
| ElavGal4/+ (Day) | 3.734 ± 0.15 | 64 | **<0.0001** |
| Nf1RNAi; Dcr-2/+ (Day) | 2.531 ± 0.16 | 32 | **<0.0001** |
| ANOVA $F_{(2, 125)}$ = 38.49 **p<0.0001** | | | |
| ElavGal4>Nf1RNAi; Dcr-2 (Night) | 7.262 ± 0.34 | 32 | |
| ElavGal4/+ (Night) | 3.789 ± 0.26 | 64 | **<0.0001** |
| Nf1RNAi; Dcr-2/+ (Night) | 4.112 ± 0.25 | 32 | **<0.0001** |
| **Fig. 3A** Kruskal Wallis $H_{(2)}$=8.763 p=0.0125 | | | |
| MB247Gal4>Nf1RNAi; Dcr-2 (Day) | 418.7 ± 7.7 | 65 | |
| MB247Gal4/+ (Day) | 448.2 ± 9.9 | 49 | 0.0533 |
| Nf1RNAi; Dcr-2/+ (Day) | 401.0 ± 11 | 69 | 0.9818 |
| Kruskal Wallis $H_{(2)}$=111.3 **p<0.0001** | | | |
| MB247Gal4>Nf1RNAi; Dcr-2 (Night) | 313.4 ± 11 | 65 | |
| MB247Gal4/+ (Night) | 304.5 ± 10 | 49 | >0.9999 |
| Nf1RNAi; Dcr-2/+ (Night) | 502.0 ± 8.4 | 69 | **<0.0001** |
| **Fig. 3B** Kruskal Wallis $H_{(2)}$=44.60 **p<0.0001** | | | |
| MB247Gal4>Nf1RNAi; Dcr-2 (Day) | 27.75 ± 0.56 | 65 | |
| MB247Gal4/+ (Day) | 27.46 ± 0.71 | 49 | >0.9999 |
| Nf1RNAi; Dcr-2/+ (Day) | 22.06 ± 0.64 | 69 | **<0.0001** |
| Kruskal Wallis $H_{(2)}$=13.53 **p=0.0012** | | | |

| | | | |
|---|---|---|---|
| MB247Gal4>Nf1RNAi; Dcr-2 (Night) | 13.39 ± 0.43 | 65 | |
| MB247Gal4/+ (Night) | 14.60 ± 0.53 | 49 | 0.2150 |
| Nf1RNAi; Dcr-2/+ (Night) | 16.28 ± 0.55 | 69 | 0.0005 |
| **Fig. 3C** Kruskal Wallis $H_{(2)}$=12.06 **p=0.0024** | | | |
| MB247Gal4>Nf1RNAi; Dcr-2 (Day) | 15.61 ± 0.52 | 65 | |
| MB247Gal4/+ (Day) | 17.13 ± 0.78 | 49 | 0.3642 |
| Nf1RNAi; Dcr-2/+ (Day) | 19.05 ± 0.73 | 69 | **0.0011** |
| Kruskal Wallis $H_{(2)}$=46.38 **p<0.0001** | | | |
| MB247Gal4>Nf1RNAi; Dcr-2 (Night) | 24.67 ± 1.1 | 65 | |
| MB247Gal4/+ (Night) | 21.80 ± 0.95 | 49 | 0.2269 |
| $Nf1^{RNAi}$;UASDcr-2/+ (Night) | 33.25 ± 1.2 | 69 | **<0.0001** |
| **Fig. 3D** Kruskal Wallis $H_{(2)}$=13.44 **p=0.0012** | | | |
| GadGal4>Nf1RNAi; Dcr-2 (Day) | 421.7 ± 26 | 38 | |
| GadGal4/+ (Day) | 323.1 ± 32 | 22 | **0.0056** |
| Nf1RNAi; Dcr-2/+ (Day) | 433.1 ± 38 | 22 | 0.7563 |
| Kruskal Wallis $H_{(2)}$=1.599 **p=0.4494** | | | |
| GadGal4>Nf1RNAi; Dcr-2 (Night) | 546.2 ± 8.2 | 38 | |
| GadGal4/+ (Night) | 525.3 ± 13 | 22 | 0.5884 |
| Nf1RNAi; Dcr-2/+ (Night) | 534.7 ± 13 | 22 | 0.5949 |
| **Fig. 3E** Kruskal Wallis $H_{(2)}$=12.39 **p=0.0020** | | | |
| GadGal4>Nf1RNAi; Dcr-2 (Day) | 22.27 ± 1.8 | 36 | |
| GadGal4/+ (Day) | 14.27 ± 0.63 | 19 | **0.0009** |
| Nf1RNAi; Dcr-2/+ (Day) | 18.61 ± 1.9 | 22 | 0.3906 |
| Kruskal Wallis $H_{(2)}$=20.10 **p<0.0001** | | | |
| GadGal4>Nf1RNAi; Dcr-2 (Night) | 43.18 ± 2.7 | 38 | |
| GadGal4/+ (Night) | 27.94 ± 1.9 | 22 | **<0.0001** |
| Nf1RNAi; Dcr-2/+ (Night) | 37.39 ± 3.0 | 22 | 0.3890 |
| **Fig. 3F** Kruskal Wallis $H_{(2)}$=12.39 **p=0.0020** | | | |
| GadGal4>Nf1RNAi; Dcr-2 (Day) | 22.27 ± 1.8 | 36 | |
| GadGal4/+ (Day) | 14.27 ± 0.63 | 19 | **0.0009** |
| Nf1RNAi; Dcr-2/+ (Day) | 18.61 ± 1.9 | 22 | 0.3906 |
| Kruskal Wallis $H_{(2)}$=20.10 **p<0.0001** | | | |
| GadGal4>Nf1RNAi; Dcr-2 (Night) | 43.18 ± 2.7 | 38 | |
| GadGal4/+ (Night) | 27.94 ± 1.9 | 22 | **<0.0001** |
| Nf1RNAi; Dcr-2/+ (Night) | 37.39 ± 3.0 | 22 | 0.3890 |
| **Fig. 3G** Kruskal Wallis $H_{(2)}$= 11.47 **p=0.0032** | | | |

| | | | |
|---|---|---|---|
| R23E10Gal4>Nf1RNAi; Dcr-2 (Day) | 383.2 ± 8.5 | 40 | |
| R23E10Gal4/+ (Day) | 351.0 ± 9.7 | 42 | 0.0471 |
| Nf1RNAi; Dcr-2/+ (Day) | 329.7 ± 12 | 50 | **0.0017** |
| Kruskal Wallis $H_{(2)}$= 16.77 **p=0.0002** | | | |
| R23E10Gal4>Nf1RNAi; Dcr-2 (Night) | 533.0 ± 10 | 40 | |
| R23E10Gal4/+ (Night) | 483.9 ± 8.9 | 42 | **0.0004** |
| Nf1RNAi; Dcr-2/+ (Night) | 524.1 ± 8.3 | 50 | >0.9999 |
| **Fig. 3H** Kruskal Wallis $H_{(2)}$=6.14 p=0.0463 | | | |
| R23E10Gal4>Nf1$^{RNAi}$;UASDcr-2 (Day) | 21.75 ± 0.62 | 40 | |
| R23E10Gal4/+ (Day) | 21.75 ± 0.62 | 42 | >0.9999 |
| Nf1$^{RNAi}$;UASDcr-2/+ (Day) | 19.76 ± 0.70 | 50 | 0.0584 |
| Kruskal Wallis $H_{(2)}$=21.69 **p<0.0001** | | | |
| R23E10Gal4>Nf1RNAi; Dcr-2 (Night) | 17.05 ± 0.66 | 40 | |
| R23E10Gal4/+ (Night) | 20.19 ± 0.86 | 42 | 0.0257 |
| Nf1RNAi; Dcr-2/+ (Night) | 15.32 ± 0.60 | 50 | 0.0954 |
| **Fig. 3I** Kruskal Wallis $H_{(2)}$=2.155 p=0.3405 | | | |
| R23E10Gal4>Nf1RNAi; Dcr-2 (Day) | 18.25 ± 0.72 | 40 | |
| R23E10Gal4/+ (Day) | 16.59 ± 0.61 | 42 | 0.2891 |
| Nf1RNAi; Dcr-2/+ (Day) | 17.24 ± 0.65 | 50 | 0.7154 |
| Kruskal Wallis $H_{(2)}$=22.78 **p<0.0001** | | | |
| R23E10Gal4>Nf1$^{RNAi}$;UASDcr-2 (Night) | 33.55 ± 1.7 | 40 | |
| R23E10Gal4/+ (Night) | 26.15 ± 1.5 | 42 | **0.0025** |
| Nf1$^{RNAi}$;UASDcr-2/+ (Night) | 36.95 ± 1.6 | 50 | 0.4448 |
| **Fig. 3J** Kruskal Wallis $H_{(2)}$=35.16 **p<0.0001** | | | |
| RdlGal4>Nf1RNAi; Dcr-2 (Day) | 384.9 ± 5.7 | 214 | |
| RdlGal4/+ (Day) | 326.5 ± 7.8 | 136 | **<0.0001** |
| Nf1RNAi; Dcr-2/+ (Day) | 391.5 ± 10 | 93 | >0.9999 |
| Kruskal Wallis $H_{(2)}$=35.63 **p<0.0001** | | | |
| RdlGal4>Nf1RNAi; Dcr-2 (Night) | 501.7 ± 4.5 | 214 | |
| RdlGal4/+ (Night) | 449.4 ± 8.4 | 136 | **<0.0001** |
| Nf1RNAi; Dcr-2/+ (Night) | 513.5 ± 7.9 | 93 | 0.3632 |
| **Fig. 3K** Kruskal Wallis $H_{(2)}$=27.80 **p<0.0001** | | | |
| RdlGal4>Nf1RNAi; Dcr-2 (Day) | 22.43 ± 0.32 | 214 | |
| RdlGal4/+ (Day) | 19.96 ± 0.44 | 136 | **<0.0001** |
| Nf1RNAi; Dcr-2/+ (Day) | 23.72 ± 0.51 | 93 | 0.2092 |
| Kruskal Wallis $H_{(2)}$=76.68 **p<0.0001** | | | |

| | | | |
|---|---|---|---|
| RdlGal4>Nf1RNAi; Dcr-2 (Night) | 16.05 ± 0.30 | 214 | |
| RdlGal4/+ (Night) | 20.86 ± 0.41 | 136 | **<0.0001** |
| Nf1RNAi; Dcr-2/+ (Night) | 17.19 ± 0.48 | 93 | 0.1327 |
| **Fig. 3L** Kruskal Wallis $H_{(2)}$=2.781 p=0.2490 | | | |
| RdlGal4>Nf1RNAi; Dcr-2 (Day) | 18.16 ± 0.49 | 214 | |
| RdlGal4/+ (Day) | 16.93 ± 0.48 | 136 | 0.2448 |
| Nf1RNAi; Dcr-2/+ (Day) | 17.07 ± 0.58 | 93 | 0.5253 |
| Kruskal Wallis $H_{(2)}$=86.42 **p<0.0001** | | | |
| RdlGal4>Nf1RNAi; Dcr-2 (Night) | 34.03 ± 0.80 | 214 | |
| RdlGal4/+ (Night) | 23.12 ± 0.77 | 136 | **<0.0001** |
| Nf1RNAi; Dcr-2/+ (Night) | 32.49 ± 1.2 | 93 | 0.5858 |
| **Fig. 4A** ANOVA $F_{(2, 231)}$ = 98.69 **p<0.0001** | | | |
| ChaGal4>Nf1RNAi; Dcr-2 | 641.2 ± 30 | 63 | |
| ChaGal4/+ | 967.5 ± 12 | 95 | **<0.0001** |
| Nf1RNAi; Dcr-2/+ | 992.0 ± 16 | 76 | **<0.0001** |
| **Fig. 4B** Kruskal Wallis $H_{(2)}$ = 63.62 **p<0.0001** | | | |
| ChaGal4>Nf1RNAi; Dcr-2 (Day) | 286.8 ± 15 | 63 | |
| ChaGal4/+ (Day) | 411.4 ± 8.4 | 95 | **<0.0001** |
| Nf1RNAi; Dcr-2/+ (Day) | 447.8 ± 11 | 76 | **<0.0001** |
| Kruskal Wallis $H_{(2)}$ = 94.31 **p<0.0001** | | | |
| ChaGal4>Nf1RNAi; Dcr-2(Night) | 364.4 ± 18 | 61 | |
| ChaGal4/+ (Night) | 556.1 ± 6.3 | 95 | **<0.0001** |
| Nf1RNAi; Dcr-2/+ (Night) | 544.2 ± 6.7 | 76 | **<0.0001** |
| Mann-Whitney (U)=731.5 **p<0.0001** | | | |
| ChaGal4/+ (Day) | 411.4 ± 8.4 | 95 | |
| ChaGal4/+ (Night) | 556.1 ± 6.3 | 95 | **<0.0001** |
| Mann-Whitney (U)=1128 **p<0.0001** | | | |
| Nf1RNAi; Dcr-2/+ (Night) | 544.2 ± 6.7 | 76 | |
| Nf1RNAi; Dcr-2/+ (Night) | 544.2 ± 6.7 | 76 | **<0.0001** |
| Mann-Whitney (U)=1250 **p=0.0007** | | | |
| ChaGal4>Nf1RNAi; Dcr-2 (Day) | 286.8 ± 15 | 63 | |
| ChaGal4>Nf1RNAi; Dcr-2(Night) | 364.4 ± 18 | 61 | **0.0007** |
| **Fig. 4C** Kruskal Wallis $H_{(2)}$ = 10.88 **p=0.0043** | | | |
| ChaGal4>Nf1RNAi; Dcr-2 (Day) | 22.84 ± 0.88 | 63 | |
| ChaGal4/+ (Day) | 19.68 ± 0.43 | 95 | **0.0044** |
| Nf1RNAi; Dcr-2/+ (Day) | 21.74 ± 0.66 | 76 | 0.8990 |

| | | | |
|---|---|---|---|
| Kruskal Wallis $H_{(2)}$ = 50.26 **p<0.0001** | | | |
| ChaGal4>Nf1RNAi; Dcr-2(Night) | 18.48 ± 0.96 | 63 | |
| ChaGal4/+ (Night) | 11.08 ± 0.45 | 95 | **<0.0001** |
| Nf1RNAi; Dcr-2/+ (Night) | 14.99 ± 0.67 | 76 | 0.0170 |
| **Fig. 4D** Kruskal Wallis $H_{(2)}$ = 66.84 **p<0.0001** | | | |
| ChaGal4>Nf1RNAi; Dcr-2 (Day) | 12.88 ± 0.63 | 63 | |
| ChaGal4/+ (Day) | 21.90 ± 0.72 | 95 | **<0.0001** |
| Nf1RNAi; Dcr-2/+ (Day) | 22.61 ± 1.1 | 76 | **<0.0001** |
| Kruskal Wallis $H_{(2)}$ = 100.9 **p<0.0001** | | | |
| ChaGal4>Nf1RNAi; Dcr-2(Night) | 20.75 ± 1.3 | 63 | |
| ChaGal4/+ (Night) | 59.12 ± 2.7 | 95 | **<0.0001** |
| Nf1RNAi; Dcr-2/+ (Night) | 43.56 ± 2.3 | 76 | **<0.0001** |
| Mann-Whitney (U)= 363 **p<0.0001** | | | |
| ChaGal4/+ (Day) | 21.90 ± 0.72 | 95 | |
| ChaGal4/+ (Night) | 59.12 ± 2.7 | 95 | **<0.0001** |
| Mann-Whitney (U)= 912 **p<0.0001** | | | |
| Nf1RNAi; Dcr-2/+ (Day) | 22.61 ± 1.1 | 76 | |
| Nf1RNAi; Dcr-2/+ (Night) | 43.56 ± 2.3 | 76 | **<0.0001** |
| Mann-Whitney (U)= 1124 **p<0.0001** | | | |
| ChaGal4>Nf1RNAi; Dcr-2 (Day) | 12.88 ± 0.63 | 63 | |
| ChaGal4>Nf1RNAi; Dcr-2(Night) | 20.75 ± 1.3 | 63 | **<0.0001** |
| **Fig. 4E** Kruskal Wallis $H_{(2)}$ = 36.38 **p<0.0001** | | | |
| ChaGal4>Nf1RNAi; Dcr-2 (Day) | 3021 ± 150 | 63 | |
| ChaGal4/+ (Day) | 4496 ± 160 | 95 | **<0.0001** |
| Nf1RNAi; Dcr-2/+ (Day) | 3786 ± 210 | 76 | 0.0158 |
| Kruskal Wallis $H_{(2)}$ = 6.892 p=0.0319 | | | |
| ChaGal4>Nf1RNAi; Dcr-2(Night) | 4898 ± 550 | 63 | |
| ChaGal4/+ (Night) | 3903 ± 200 | 95 | >0.9999 |
| Nf1RNAi; Dcr-2/+ (Night) | 4544 ± 210 | 76 | 0.0413 |
| **Fig. 4F** Kruskal Wallis $H_{(2)}$ = 70.91 **p<0.0001** | | | |
| ChaGal4>Nf1RNAi; Dcr-2 (Day) | 2.562 ± 0.11 | 63 | |
| ChaGal4/+ (Day) | 4.297 ± 0.13 | 95 | **<0.0001** |
| Nf1RNAi; Dcr-2/+ (Day) | 3.709 ± 0.17 | 76 | **<0.0001** |
| Kruskal Wallis $H_{(2)}$ = 15.17 **p=0.0005** | | | |
| ChaGal4>Nf1RNAi; Dcr-2(Night) | 4.147 ± 0.40 | 63 | |
| ChaGal4/+ (Night) | 4.307 ± 0.19 | 95 | 0.1197 |
| Nf1RNAi; Dcr-2/+ (Night) | 4.992 ± 0.21 | 76 | **0.0002** |
| **Fig. 5B** Kruskal Wallis $_{(2)}$ = 202.3 **p<0.0001** | | | |

| | | | |
|---|---|---|---|
| ChaGal4/+ | 960.6 ± 11 | 103 | |
| UASNf1; $Nf1^{Null}$ | 235.9 ± 16 | 72 | **<0.0001** |
| ChaGal4>dUASNf1;$Nf1^{Null}$ | 769.1 ± 14 | 110 | **<0.0001** |
| ChaGal4>UASdNf1; Nf1Null | 769.1 ± 14 | 110 | |
| UASdNf1 $Nf1^{Null}$ | 235.9 ± 16 | 72 | **<0.0001** |
| **Fig. 5C** Kruskal Wallis $_{(2)}$ = 173.1 **p<0.0001** | | | |
| ChaGal4/+ (Day) | 440.3 ± 7.6 | 103 | |
| UASdNf1; Nf1Null (Day) | 91.95 ± 8.0 | 72 | **<0.0001** |
| ChaGal4>UASdNf1; Nf1Null (Day) | 358.7 ± 10 | 110 | **<0.0001** |
| ChaGal4>UASdNf1; Nf1Null (Day) | 358.7 ± 10 | 110 | |
| UASdNf1; Nf1Null (Day) | 91.95 ± 8.0 | 72 | **<0.0001** |
| Kruskal Wallis $_{(2)}$ = 203.4 **p<0.0001** | | | |
| ChaGal4/+ (Night) | 520.2 ± 7.3 | 103 | |
| UASdNf1; Nf1Null (Night) | 145.8 ± 9.2 | 72 | **<0.0001** |
| ChaGal4>UASdNf1; Nf1Null (Night) | 410.2 ± 6.3 | 110 | **<0.0001** |
| ChaGal4>UASdNf1; Nf1Null (Night) | 410.2 ± 6.3 | 110 | |
| UASdNf1; Nf1Null (Night) | 145.8 ± 9.2 | 72 | **<0.0001** |
| Mann-Whitney (U)=2400 **p<0.0001** | | | |
| ChaGal4/+ (Day) | 440.3 ± 7.6 | 103 | |
| ChaGal4/+ (Night) | 520.2 ± 7.3 | 103 | **<0.0001** |
| Mann-Whitney (U)=4152 **p<0.0001** | | | |
| ChaGal4>UASdNf1; Nf1Null (Day) | 358.7 ± 10 | 110 | |
| ChaGal4>UASdNf1; Nf1Null (Night) | 410.2 ± 6.3 | 110 | **<0.0001** |
| **Fig. 5D** Kruskal Wallis $_{(2)}$ =127.9 **p<0.0001** | | | |
| ChaGal4/+ (Day) | 22.31 ± 0.49 | 103 | |
| UASNf1 ;Nf1Null (Day) | 11.56 ± 0.79 | 72 | **<0.0001** |
| ChaGal4>UASdNf1; Nf1Null (Day) | 26.02 ± 0.55 | 110 | **<0.0001** |
| ChaGal4>UASdNf1; Nf1Null (Day) | 26.02 ± 0.55 | 110 | |
| UASdNf1; Nf1Null (Day) | 11.56 ± 0.79 | 72 | **<0.0001** |
| Kruskal Wallis $_{(2)}$ =49.73 **p<0.0001** | | | |
| ChaGal4/+ (Night) | 14.87 ± 0.47 | 103 | |
| UASdNf1; Nf1Null (Night) | 17.13 ± 0.82 | 72 | **0.0081** |
| ChaGal4>UASdNf1; Nf1Null (Night) | 19.75 ± 0.40 | 110 | **<0.0001** |

| | | | |
|---|---|---|---|
| ChaGal4>UASdNf1; Nf1Null (Night) | 19.75 ± 0.40 | 110 | |
| UASdNf1; Nf1Null (Night) | 17.13 ± 0.82 | 72 | **0.0026** |
| **Fig. 5E** Kruskal Wallis $_{(2)}$ = 187.6 **p<0.0001** | | | |
| ChaGal4/+ (Day) | 21.11 ± 0.80 | 103 | |
| UASdNf1; Nf1Null (Day) | 7.374 ± 0.19 | 72 | **<0.0001** |
| ChaGal4>UASdNf1; Nf1Null (Day) | 14.34 ± 0.59 | 110 | **<0.0001** |
| ChaGal4>UASdNf1; Nf1Null (Day) | 14.34 ± 0.59 | 110 | |
| UASdNf1; Nf1Null (Day) | 7.374 ± 0.19 | 72 | **<0.0001** |
| Kruskal Wallis $_{(2)}$ = 206.7 **p<0.0001** | | | |
| ChaGal4/+ (Night) | 39.65 ± 1.7 | 103 | |
| UASdNf1; Nf1Null (Night) | 8.043 ± 0.21 | 72 | **<0.0001** |
| ChaGal4>UASdNf1; Nf1Null (Night) | 22.11 ± 0.72 | 110 | **<0.0001** |
| ChaGal4>UASdNf1; Nf1Null (Night) | 22.11 ± 0.72 | 110 | |
| UASdNf1; Nf1Null (Night) | 8.043 ± 0.21 | 72 | **<0.0001** |
| Mann-Whitney (U)=1249 **p<0.0001** | | | |
| ChaGal4/+ (Day) | 21.11 ± 0.80 | 103 | |
| ChaGal4/+ (Night) | 39.65 ± 1.7 | 103 | **<0.0001** |
| Mann-Whitney (U)=1930 **p<0.0001** | | | |
| ChaGal4>UASdNf1; Nf1Null (Day) | 14.34 ± 0.59 | 110 | |
| ChaGal4>UASdNf1; Nf1Null (Night) | 22.11 ± 0.72 | 110 | **<0.0001** |
| **Fig. 6A** Kruskal Wallis $_{(2)}$ = 9.629 **p=0.0030** | | | |
| ChaGal4/+ | 0.2589 ± 0.076 | 11 | |
| UASdNf1-GRD; Nf1Null | 1.367 ± 0.38 | 4 | **0.0043** |
| ChaGal4>UASdNf1-GRD; Nf1Null | 0.4140 ± 0.079 | 6 | 0.3317 |
| **Fig. 6C** Kruskal Wallis $_{(4)}$ =357.6 **p<0.0001** | | | |
| ChaGal4/+ | 917.8 ± 9.8 | 163 | |
| UASdNf1-GRD-RP; Nf1Null | 319.7 ± 20 | 80 | **<0.0001** |
| UASdNf1-GRD; Nf1Null | 368.1 ± 22 | 101 | **<0.0001** |
| ChaGal4>UASdNf1-GRD-RP; Nf1Null | 273.1 ± 20 | 76 | **<0.0001** |
| ChaGal4>UASdNf1-GRD; Nf1Null | 720.1 ± 17 | 112 | **<0.0001** |
| UASdNf1-GRD; Nf1Null | 368.1 ± 22 | 101 | |
| ChaGal4>UASdNf1-GRD; Nf1Null | 720.1 ± 17 | 112 | **<0.0001** |

| | | | |
|---|---|---|---|
| UASdNf1-GRD-RP; Nf1Null | 319.7 ± 20 | 80 | |
| ChaGal4>UASdNf1-GRD-RP; Nf1Null | 273.1 ± 20 | 76 | >0.9999 |
| **Fig. 6D** Kruskal Wallis $_{(4)}$ =312.2 **p<0.0001** | | | |
| ChaGal4/+ (Day) | 406.2 ± 6.9 | 163 | |
| UASdNf1-GRD-RP; Nf1Null (Day) | 147.0 ± 10 | 80 | **<0.0001** |
| UASdNf1-GRD; Nf1Null (Day) | 148.5 ± 10 | 101 | **<0.0001** |
| ChaGal4>UASdNf1-GRD-RP; Nf1Null (Day) | 176.8 ± 11 | 76 | **<0.0001** |
| ChaGal4>UASdNf1-GRD; Nf1Null (Day) | 344.0 ± 10 | 112 | **0.0076** |
| UASdNf1-GRD; Nf1Null (Day) | 148.5 ± 10 | 101 | |
| ChaGal4>UASdNf1-GRD; Nf1Null (Day) | 344.0 ± 10 | 112 | **<0.0001** |
| UASdNf1-GRD-RP; Nf1Null (Day) | 147.0 ± 10 | 80 | |
| ChaGal4>UASdNf1-GRD-RP; Nf1Null (Day) | 176.8 ± 11 | 76 | >0.9999 |
| Kruskal Wallis $_{(4)}$ =360.3 **p<0.0001** | | | |
| ChaGal4/+ (Night) | 511.3 ± 6.1 | 163 | |
| UASdNf1-GRD-RP; Nf1Null (Night) | 172.6 ± 11 | 80 | **<0.0001** |
| UASdNf1-GRD; Nf1Null (Night) | 219.3 ± 14 | 101 | **<0.0001** |
| ChaGal4>UASdNf1-GRD-RP; Nf1Null (Night) | 96.19 ± 11 | 76 | **<0.0001** |
| ChaGal4>UASdNf1-GRD; Nf1Null (Night) | 375.8 ± 10 | 112 | **<0.0001** |
| UASdNf1-GRD; Nf1Null (Night) | 219.3 ± 14 | 101 | |
| ChaGal4>UASdNf1-GRD; Nf1Null (Night) | 375.8 ± 10 | 112 | **<0.0001** |
| UASdNf1-GRD-RP; Nf1Null (Night) | 172.6 ± 11 | 80 | |
| ChaGal4>UASdNf1-GRD-RP; Nf1Null (Night) | 96.19 ± 11 | 76 | 0.0162 |
| Mann-Whitney (U)=4794 **p<0.0001** | | | |
| ChaGal4/+ (Day) | 406.2 ± 6.9 | 163 | |
| ChaGal4/+ (Night) | 511.3 ± 6.1 | 163 | **<0.0001** |
| Mann-Whitney (U)=5195 p=0.0264 | | | |
| ChaGal4>UASdNf1-GRD; Nf1Null (Day) | 344.0 ± 10 | 112 | |
| ChaGal4>UASdNf1-GRD; Nf1Null (Night) | 375.8 ± 10 | 112 | 0.0264 |
| **Fig. 6E** Kruskal Wallis $_{(4)}$ =146.9 **p<0.0001** | | | |

| | | | |
|---|---|---|---|
| ChaGal4/+ (Day) | 21.03 ± 0.42 | 163 | |
| UASdNf1-GRD-RP; Nf1Null (Day) | 15.89 ± 0.59 | 80 | **<0.0001** |
| UASdNf1-GRD; Nf1Null (Day) | 15.71 ± 0.70 | 101 | **<0.0001** |
| ChaGal4>UASdNf1-GRD-RP; Nf1Null (Day) | 16.98 ± 0.64 | 76 | **0.001** |
| ChaGal4>UASdNf1-GRD; Nf1Null (Day) | 25.29 ± 0.62 | 112 | **<0.0001** |
| UASdNf1-GRD; Nf1Null (Day) | 15.71 ± 0.70 | 101 | |
| ChaGal4>UASdNf1-GRD; Nf1Null (Day) | 25.29 ± 0.62 | 112 | **<0.0001** |
| UASdNf1-GRD-RP; Nf1Null (Day) | 15.89 ± 0.59 | 80 | |
| ChaGal4>UASdNf1-GRD-RP; Nf1Null (Day) | 16.98 ± 0.64 | 76 | >0.9999 |
| Kruskal Wallis $_{(4)}$ =185.2 **p<0.0001** | | | |
| ChaGal4/+ (Night) | 13.36 ± 0.37 | 163 | |
| UASdNf1-GRD-RP; Nf1Null (Night) | 19.31 ± 0.64 | 80 | **<0.0001** |
| UASdNf1-GRD; Nf1Null (Night) | 20.35 ± 0.79 | 101 | **<0.0001** |
| ChaGal4>UASdNf1-GRD-RP; Nf1Null (Night) | 10.07 ± 0.71 | 76 | 0.0433 |
| ChaGal4>UASdNf1-GRD; Nf1Null (Night) | 21.45 ± 0.52 | 112 | **<0.0001** |
| UASdNf1-GRD; Nf1Null (Night) | 20.35 ± 0.79 | 101 | |
| ChaGal4>UASdNf1-GRD; Nf1Null (Night) | 21.45 ± 0.52 | 112 | 0.9209 |
| UASdNf1-GRD-RP; Nf1Null (Night) | 19.31 ± 0.64 | 80 | |
| ChaGal4>UASdNf1-GRD-RP; Nf1Null (Night) | 10.07 ± 0.71 | 76 | **<0.0001** |
| **Fig. 6F** Kruskal Wallis $_{(4)}$ =309.0 **p<0.0001** | | | |
| ChaGal4/+ (Day) | 21.14 ± 0.77 | 163 | |
| UASdNf1-GRD-RP; Nf1Null (Day) | 8.841 ± 0.41 | 80 | **<0.0001** |
| UASdNf1-GRD; Nf1Null (Day) | 8.776 ± 0.32 | 101 | **<0.0001** |
| ChaGal4>UASdNf1-GRD-RP; Nf1Null (Day) | 10.14 ± 0.44 | 76 | **<0.0001** |
| ChaGal4>UASdNf1-GRD; Nf1Null (Day) | 15.55 ± 1.6 | 112 | **<0.0001** |
| UASdNf1-GRD; Nf1Null (Day) | 8.776 ± 0.32 | 101 | |
| ChaGal4>UASdNf1-GRD; Nf1Null (Day) | 15.55 ± 1.6 | 112 | **<0.0001** |
| UASdNf1-GRD-RP; Nf1Null (Day) | 8.841 ± 0.41 | 80 | |
| ChaGal4>UASdNf1-GRD-RP; Nf1Null (Day) | 10.14 ± 0.44 | 76 | 0.1258 |
| Kruskal Wallis $_{(4)}$ =395.8 **p<0.0001** | | | |
| ChaGal4/+ (Night) | 43.69 ± 1.6 | 163 | |
| UASdNf1-GRD-RP; Nf1Null (Night) | 9.161 ± 0.84 | 80 | **<0.0001** |
| UASdNf1-GRD; Nf1Null (Night) | 10.55 ± 0.78 | 101 | **<0.0001** |
| ChaGal4>UASdNf1-GRD-RP; Nf1Null (Night) | 8.647 ± 0.51 | 76 | **<0.0001** |
| ChaGal4>UASdNf1-GRD; Nf1Null (Night) | 18.14 ± 0.73 | 112 | **<0.0001** |

| | | | |
|---|---|---|---|
| UASdNf1-GRD; Nf1Null (Night) | 10.55 ± 0.78 | 101 | |
| ChaGal4>UASdNf1-GRD; Nf1Null (Night) | 18.14 ± 0.73 | 112 | **<0.0001** |
| UASdNf1-GRD-RP; Nf1Null (Night) | 9.161 ± 0.84 | 80 | |
| ChaGal4>UASdNf1-GRD-RP; Nf1Null (Night) | 8.647 ± 0.51 | 76 | >0.9999 |
| Mann-Whitney (U)=2762 **p<0.0001** | | | |
| ChaGal4/+ (Day) | 21.14 ± 0.77 | 163 | |
| ChaGal4/+ (Night) | 43.69 ± 1.6 | 163 | **<0.0001** |
| Mann-Whitney (U)=3408 **p<0.0001** | | | |
| ChaGal4>UASdNf1-GRD; Nf1Null (Day) | 15.55 ± 1.6 | 112 | |
| ChaGal4>UASdNf1-GRD; Nf1Null (Night) | 18.14 ± 0.73 | 112 | **<0.0001** |
| **Fig. 7B** ANOVA $F_{(3,29)} = 6.998$ **p=0.0011** | | | |
| $w^{1118}$ | 1.000 ± 0 | 9 | |
| $Nf1^{Null}$ | 4.478 ± 0.75 | 9 | **0.0003** |
| $Nf1^{C1045Y}$ | 3.127 ± 0.55 | 9 | **0.0191** |
| $Nf1^{R1861C}$ | 3.068 ± 0.73 | 6 | **0.0224** |
| **Fig. 7D** Kruskal Wallis $H_{(3)} = 114.1$ **p<0.0001** | | | |
| $w^{1118}$ | 668.3 ± 28 | 156 | |
| $Nf1^{Null}$ | 274.6 ± 24 | 127 | **<0.0001** |
| $Nf1^{C1045Y}$ | 377.5 ± 34 | 60 | **<0.0001** |
| $Nf1^{R1861C}$ | 532.1 ± 18 | 158 | **0.0009** |
| Mann-Whitney (U)= 2817 **p<0.0001** | | | |
| $Nf1^{C1045Y}$ | 377.5 ± 34 | 60 | |
| $Nf1^{R1861C}$ | 532.1 ± 18 | 158 | **<0.0001** |
| **Fig. 7E** Kruskal Wallis $H_{(3)} = 91.24$ **p<0.0001** | | | |
| $w^{1118}$ (Day) | 263.9 ± 13 | 156 | |
| $Nf1^{Null}$ (Day) | 123.0 ± 11 | 127 | **<0.0001** |
| $Nf1^{C1045Y}$ (Day) | 200.9 ± 18 | 60 | 0.0274 |
| $Nf1^{R1861C}$ (Day) | 262.2 ± 9.0 | 158 | >0.9999 |
| Mann-Whitney (U)= 3205 **p=0.0002** | | | |
| $Nf1^{C1045Y}$ (Day) | 200.9 ± 18 | 60 | |
| $Nf1^{R1861C}$ (Day) | 262.2 ± 9.0 | 158 | **0.0002** |
| Kruskal Wallis $H_{(3)} = 129.1$ **p<0.0001** | | | |
| $w^{1118}$ (Night) | 404.3 ± 16 | 156 | |
| $Nf1^{Null}$ (Night) | 151.5 ± 13 | 127 | **<0.0001** |
| $Nf1^{C1045Y}$ (Night) | 176.5 ± 19 | 60 | **<0.0001** |

| | | | |
|---|---|---|---|
| $Nf1^{R1861C}$ (Night) | 269.6 ± 10 | 158 | **<0.0001** |
| Mann-Whitney (U)= 2660 **p<0.0001** | | | |
| $Nf1^{C1045Y}$ (Night) | 176.5 ± 19 | 60 | |
| $Nf1^{R1861C}$ (Night) | 269.6 ± 10 | 158 | **0.0001** |
| **Fig. 7F** Kruskal Wallis $H_{(3)}$ = 88.98 **p<0.0001** | | | |
| $w^{1118}$ (Day) | 19.22 ± 0.73 | 156 | |
| $Nf1^{Null}$ (Day) | 12.90 ± 0.87 | 127 | **<0.0001** |
| $Nf1^{C1045Y}$ (Day) | 18.24 ± 1.0 | 60 | >0.9999 |
| $Nf1^{R1861C}$ (Day) | 24.07 ± 0.57 | 158 | **<0.0001** |
| Mann-Whitney (U)= 2780 **p<0.0001** | | | |
| $Nf1^{C1045Y}$ (Day) | 18.24 ± 1.0 | 60 | |
| $Nf1^{R1809C}$ (Day) | 24.07 ± 0.57 | 158 | **<0.0001** |
| Kruskal Wallis $H_{(3)}$ = 81.58 **p<0.0001** | | | |
| $w^{1118}$ (Night) | 21.16 ± 0.87 | 156 | |
| $Nf1^{Null}$ (Night) | 15.57 ± 1.1 | 127 | **0.0003** |
| $Nf1^{C1045Y}$ (Night) | 14.09 ± 0.89 | 60 | **0.0001** |
| $Nf1^{R1861C}$ (Night) | 26.04 ± 0.77 | 158 | **0.0002** |
| Mann-Whitney (U)= 1563 **p<0.0001** | | | |
| $Nf1^{C1045Y}$ (Night) | 14.09 ± 0.89 | 60 | |
| $Nf1^{R1861C}$ (Night) | 26.04 ± 0.77 | 158 | **0.0001** |
| **Fig. 7G** Kruskal Wallis $H_{(3)}$ = 87.00 **p<0.0001** | | | |
| $w^{1118}$ (Day) | 11.69 ± 0.47 | 150 | |
| $Nf1^{Null}$ (Day) | 8.110 ± 0.40 | 126 | **<0.0001** |
| $Nf1^{C1045Y}$ (Day) | 10.76 ± 1.0 | 60 | 0.0285 |
| $Nf1^{R1861C}$ (Day) | 10.53 ± 0.24 | 158 | >0.9999 |
| Mann-Whitney (U)= 3526 **p=0.0034** | | | |
| $Nf1^{C1045Y}$ (Day) | 10.76 ± 1.0 | 60 | |
| $Nf1^{R1861C}$ (Day) | 10.53 ± 0.24 | 158 | **0.0034** |
| Kruskal Wallis $H_{(3)}$ =126.8 **p<0.0001** | | | |
| $w^{1118}$ (Night) | 16.77 ± 0.68 | 149 | |
| $Nf1^{Null}$ (Night) | 8.420 ± 0.42 | 126 | **<0.0001** |
| $Nf1^{C1045Y}$ (Night) | 12.03 ± 1.3 | 59 | **<0.0001** |
| $Nf1^{R1861C}$ (Night) | 10.78 ± 0.73 | 158 | **<0.0001** |
| Mann-Whitney (U)= 4438 **p=0.5893** | | | |
| $Nf1^{C1045Y}$ (Night) | 12.03 ± 1.3 | 59 | |
| $Nf1^{R1861C}$ (Night) | 10.78 ± 0.73 | 158 | 0.5893 |
| **Fig. 7H** Kruskal Wallis $H_{(3)}$ =93.92 **p<0.0001** | | | |
| $w^{1118}$ (Day) | 5443 ± 220 | 156 | |
| $Nf1^{Null}$ (Day) | 5712 ± 150 | 127 | 0.1528 |
| $Nf1^{C1045Y}$ (Day) | 3657 ± 190 | 60 | **<0.0001** |
| $Nf1^{R1861C}$ (Day) | 4012 ± 120 | 158 | **<0.0001** |
| Mann-Whitney (U)= 3973 p=0.0650 | | | |

| | | | |
|---|---|---|---|
| $Nf1^{C1045Y}$ (Day) | 3657 ± 190 | 60 | |
| $Nf1^{R1861C}$ (Day) | 4012 ± 120 | 158 | 0.0650 |
| Kruskal Wallis $H_{(3)}$ =61.81 **p<0.0001** | | | |
| $w^{1118}$ (Night) | 3379 ± 180 | 156 | |
| $Nf1^{Null}$ (Night) | 5068 ± 170 | 127 | **<0.0001** |
| $Nf1^{C1045Y}$ (Night) | 5031 ± 340 | 60 | **<0.0001** |
| $Nf1^{R1861C}$ (Night) | 3888 ± 120 | 158 | 0.0177 |
| Mann-Whitney (U)= 3337 p=0.0007 | | | |
| $Nf1^{C1045Y}$ (Night) | 5031 ± 340 | 60 | |
| $Nf1^{R1861C}$ (Night) | 3888 ± 120 | 158 | **0.0007** |
| Mann-Whitney (U)=6588 **p<0.0001** | | | |
| $w^{1118}$ (Day) | 5443 ± 220 | 156 | |
| $w^{1118}$ (Night) | 3379 ± 180 | 156 | **0.0001** |
| Mann-Whitney (U)= 6431 **p=0.0053** | | | |
| $Nf1^{Null}$ (Day) | 5712 ± 150 | 127 | |
| $Nf1^{Null}$ (Night) | 5068 ± 170 | 127 | **0.0053** |
| Mann-Whitney (U)=1144 **p=0.0005** | | | |
| $Nf1^{C1045Y}$ (Day) | 3657 ± 190 | 60 | |
| $Nf1^{C1045Y}$ (Night) | 5031 ± 340 | 60 | **0.0005** |
| Mann-Whitney (U)=11602 p=0.2785 | | | |
| $Nf1^{R1861C}$ (Day) | 4012 ± 120 | 158 | |
| $Nf1^{R1861C}$ (Night) | 3888 ± 120 | 158 | 0.2785 |
| **Fig. 7I** Kruskal Wallis $H_{(3)}$ =100.9 **p<0.0001** | | | |
| $w^{1118}$ (Day) | 4.492 ± 0.17 | 156 | |
| $Nf1^{Null}$ (Day) | 4.272 ± 0.089 | 127 | >0.9999 |
| $Nf1^{C1045Y}$ (Day) | 2.867 ± 0.12 | 60 | **<0.0001** |
| $Nf1^{R1861C}$ (Day) | 3.372 ± 0.088 | 158 | **<0.0001** |
| Mann-Whitney (U)=3274 **p=0.0004** | | | |
| $Nf1^{C1045Y}$ (Day) | 2.867 ± 0.12 | 60 | |
| $Nf1^{R1861C}$ (Day) | 3.372 ± 0.088 | 158 | **0.0004** |
| Kruskal Wallis $H_{(3)}$ =30.77 **p<0.0001** | | | |
| $w^{1118}$ (Night) | 3.050 ± 0.13 | 156 | |
| $Nf1^{Null}$ (Night) | 3.839 ± 0.10 | 127 | **<0.0001** |
| $Nf1^{C1045Y}$ (Night) | 3.824 ± 0.23 | 60 | **0.0039** |
| $Nf1^{R1861C}$ (Night) | 3.261 ± 0.087 | 158 | 0.4988 |
| Mann-Whitney (U)=3793 **p=0.0226** | | | |

| | | | |
|---|---|---|---|
| $Nf1^{C1045Y}$ (Night) | 3.824 ± 0.23 | 60 | |
| $Nf1^{R1861C}$ (Night) | 3.261 ± 0.087 | 158 | **0.0226** |
| Mann-Whitney (U)=6754 **p<0.0001** | | | |
| $w^{1118}$ (Day) | 4.492 ± 0.17 | 156 | |
| $w^{1118}$ (Night) | 3.050 ± 0.13 | 156 | **<0.0001** |
| Mann-Whitney (U)= 6189 **p=0.0014** | | | |
| $Nf1^{Null}$ (Day) | 4.272 ± 0.089 | 127 | |
| $Nf1^{Null}$ (Night) | 3.839 ± 0.10 | 127 | **0.0014** |
| Mann-Whitney (U)=1106 **p=0.0002** | | | |
| $Nf1^{C1045Y}$ (Day) | 2.867 ± 0.12 | 60 | |
| $Nf1^{C1045Y}$ (Night) | 3.824 ± 0.23 | 60 | **0.0002** |
| Mann-Whitney (U)=11652 p=0.3070 | | | |
| $Nf1^{R1861C}$ (Day) | 3.372 ± 0.088 | 158 | |
| $Nf1^{R1861C}$ (Night) | 3.261 ± 0.087 | 158 | 0.3070 |
| **Fig. 8A** ANOVA $F_{(2, 309)} = 17.47$ **p < 0.0001** | | | |
| ElavGal4/+ (Day) | 387.7 ± 11 | 92 | |
| UASdNf1; $Nf1^{C1045Y}$(Day) | 300.2 ± 11 | 83 | **<0.0001** |
| ElavGal4>UASdNf1; $Nf1^{C1045Y}$(Day) | 319.0 ± 9.3 | 137 | **<0.0001** |
| ElavGal4>UASdNf1; $Nf1^{C1045Y}$(Day) | 319.0 ± 9.3 | 137 | |
| UASdNf1; $Nf1^{C1045Y}$(Day) | 300.2 ± 11 | 83 | 0.2024 |
| ANOVA $F_{(2, 308)} = 165.1$ **p<0.0001** | | | |
| ElavGal4/+ (Night) | 506.0 ± 6.9 | 91 | |
| UASdNf1; $Nf1^{C1045Y}$ (Night) | 326.4 ± 13 | 83 | **<0.0001** |
| ElavGal4>UASdNf1; $Nf1^{C1045Y}$ (Night) | 271.2 ± 8.7 | 137 | **<0.0001** |
| ElavGal4>UASdNf1; $Nf1^{C1045Y}$ (Night) | 271.2 ± 8.7 | 137 | |
| UASdNf1; $Nf1^{C1045Y}$ (Night) | 326.4 ± 13 | 83 | **<0.0001** |
| t=9.256, df=154.4 **p<0.0001** | | | |
| ElavGal4/+ (Day) | 387.7 ± 11 | 92 | |
| ElavGal4/+ (Night) | 506.0 ± 6.9 | 91 | **<0.0001** |
| t=3.757, df=270.9 **p = 0.0002** | | | |
| ElavGal4>UASdNf1; $Nf1^{C1045Y}$ (Day) | 319.0 ± 9.3 | 137 | |
| ElavGal4>UASdNf1; $Nf1^{C1045Y}$ (Night) | 271.2 ± 8.7 | 137 | **0.0002** |

| | | | |
|---|---|---|---|
| **Fig. 8B** ANOVA $F_{(2, 309)} = 63.35$ **p<0.0001** | | | |
| ElavGal4/+ (Day) | 16.07 ± 0.50 | 92 | |
| UASdNf1; Nf1$^{C1045Y}$ (Day) | 24.50 ± 0.66 | 83 | **<0.0001** |
| ElavGal4>UASdNf1; Nf1$^{C1045Y}$ (Day) | 22.87 ± 0.46 | 137 | **<0.0001** |
| ElavGal4>UASdNf1; Nf1$^{C1045Y}$ (Day) | 22.87 ± 0.46 | 137 | |
| UASdNf1; Nf1$^{C1045Y}$ (Day) | 24.50 ± 0.66 | 83 | 0.0313 |
| ANOVA $F_{(2, 304)} = 73.65$ **p<0.0001** | | | |
| ElavGal4/+ (Night) | 16.51 ± 0.53 | 92 | |
| UASdNf1; Nf1$^{C1045Y}$ (Night) | 27.12 ± 0.61 | 78 | **<0.0001** |
| ElavGal4>UASdNf1; Nf1$^{C1045Y}$ (Night) | 21.76 ± 0.53 | 137 | **<0.0001** |
| ElavGal4>UASdNf1; Nf1$^{C1045Y}$ (Night) | 21.76 ± 0.53 | 137 | |
| UASdNf1; Nf1$^{C1045Y}$ (Night) | 27.12 ± 0.61 | 78 | **<0.0001** |
| **Fig. 8C** Kruskal Wallis $_{(2)}$ = 116.9 **p<0.0001** | | | |
| ElavGal4/+ (Day) | 25.92 ± 1.3 | 90 | |
| UASdNf1; Nf1$^{C1045Y}$ (Day) | 12.35 ± 0.46 | 83 | **<0.0001** |
| ElavGal4>UASdNf1; Nf1$^{C1045Y}$ (Day) | 14.04 ± 0.37 | 137 | **<0.0001** |
| ElavGal4>UASdNf1; Nf1$^{C1045Y}$ (Day) | 14.04 ± 0.37 | 137 | |
| UASdNf1; Nf1$^{C1045Y}$(Day) | 12.35 ± 0.46 | 83 | 0.0167 |
| Kruskal Wallis $_{(2)}$ = 179.3 **p<0.0001** | | | |
| ElavGal4/+ (Night) | 32.91 ± 1.2 | 91 | |
| UASdNf1; Nf1$^{C1045Y}$ (Night) | 13.78 ± 1.2 | 83 | **<0.0001** |
| ElavGal4>UASdNf1; Nf1$^{C1045Y}$ (Night) | 12.33 ± 0.25 | 137 | **<0.0001** |
| ElavGal4>UASdNf1; Nf1$^{C1045Y}$ (Night) | 12.33 ± 0.25 | 137 | |
| UASdNf1; Nf1$^{C1045Y}$ (Night) | 13.78 ± 1.2 | 83 | >0.9999 |
| Mann-Whitney (U)=2422 **p<0.0001** | | | |
| ElavGal4/+ (Day) | 25.92 ± 1.3 | 90 | |
| ElavGal4/+ (Night) | 32.91 ± 1.2 | 91 | **<0.0001** |
| Mann-Whitney (U)= 7359 **p=0.0020** | | | |
| ElavGal4>UASdNf1; Nf1$^{C1045Y}$ (Day) | 14.04 ± 0.37 | 137 | |
| ElavGal4>UASdNf1; Nf1$^{C1045Y}$ (Night) | 12.33 ± 0.25 | 137 | **0.0020** |
| **Fig. 8D** Kruskal Wallis $_{(3)}$ = 23.70 **p<0.0001** | | | |
| ElavGal4/+ (Day) | 394.0 ± 12 | 53 | |
| ElavGal4; Nf1$^{C1045Y}$ (Day) | 378.7 ± 16 | 58 | >0.9999 |
| ElavGal4>UASdNf1-GRD; Nf1$^{C1045Y}$ (Day) | 392.5 ± 15 | 57 | >0.9999 |
| ElavGal4>UASdNf1-GRD-RP; Nf1$^{C1045Y}$ (Day) | 315.1 ± 14 | 90 | **0.0006** |

| | | | |
|---|---|---|---|
| ElavGal4 ; $Nf1^{C1045Y}$ (Day) | 378.7 ± 16 | 58 | |
| ElavGal4>UASdNf1-GRD; $Nf1^{C1045Y}$ (Day) | 392.5 ± 15 | 57 | 0.0193 |
| ElavGal4>UASdNf1-GRD-RP; $Nf1^{C1045Y}$ (Day) | 315.1 ± 14 | 90 | >0.9999 |
| Kruskal Wallis $_{(3)}$ = 116.8 **p<0.0001** | | | |
| ElavGal4/+ (Night) | 550.0 ± 11 | 53 | |
| ElavGal4; $Nf1^{C1045Y}$ (Night) | 335.2 ± 17 | 58 | **<0.0001** |
| ElavGal4>UASdNf1-GRD; $Nf1^{C1045Y}$ (Night) | 370.4 ± 13 | 57 | **<0.0001** |
| ElavGal4>UASdNf1-GRD-RP; $Nf1^{C1045Y}$ (Night) | 264.1 ± 13 | 90 | **<0.0001** |
| ElavGal4; $Nf1^{C1045Y}$ (Night) | 335.2 ± 17 | 58 | |
| ElavGal4>UASdNf1-GRD; $Nf1^{C1045Y}$ (Night) | 370.4 ± 13 | 57 | 0.7402 |
| ElavGal4>UASdNf1-GRD-RP; $Nf1^{C1045Y}$ (Night) | 264.1 ± 13 | 90 | 0.0401 |
| Mann-Whitney (U)=267 **p<0.0001** | | | |
| ElavGal4/+ (Day) | 394.0 ± 12 | 53 | |
| ElavGal4/+ (Night) | 550.0 ± 11 | 53 | **<0.0001** |
| Mann-Whitney (U)= 1379 p=0.1643 | | | |
| ElavGal4>UASdNf1-GRD; $Nf1^{C1045Y}$ (Day) | 392.5 ± 15 | 57 | |
| ElavGal4>UASdNf1-GRD; $Nf1^{C1045Y}$ (Night) | 370.4 ± 13 | 57 | 0.1643 |
| **Fig. 8E** Kruskal Wallis $_{(3)}$ = 38.42 **p<0.0001** | | | |
| ElavGal4/+ (Day) | 18.58 ± 0.70 | 53 | |
| ElavGal4; $Nf1^{C1045Y}$ (Day) | 25.75 ± 0.81 | 58 | **<0.0001** |
| ElavGal4>UASdNf1-GRD; $Nf1^{C1045Y}$ (Day) | 22.68 ± 0.77 | 57 | **0.0053** |
| ElavGal4>UASdNf1-GRD-RP; $Nf1^{C1045Y}$ (Day) | 23.50 ± 0.73 | 90 | **<0.0001** |
| ElavGal4; $Nf1^{C1045Y}$ (Day) | 25.75 ± 0.81 | 58 | |
| ElavGal4>UASdNf1-GRD; $Nf1^{C1045Y}$ (Day) | 22.68 ± 0.77 | 57 | 0.0359 |
| ElavGal4>UASdNf1-GRD-RP; $Nf1^{C1045Y}$ (Day) | 23.50 ± 0.73 | 90 | 0.2137 |
| Kruskal Wallis $_{(3)}$ = 65.58 **p<0.0001** | | | |
| ElavGal4/+ (Night) | 15.98 ± 0.63 | 53 | |
| ElavGal4; $Nf1^{C1045Y}$ (Night) | 24.75 ± 0.90 | 58 | **<0.0001** |
| ElavGal4>UASdNf1-GRD; $Nf1^{C1045Y}$ (Night) | 25.32 ± 0.66 | 57 | **<0.0001** |
| ElavGal4>UASdNf1-GRD-RP; $Nf1^{C1045Y}$ (Night) | 22.72 ± 0.76 | 90 | **<0.0001** |
| ElavGal4; $Nf1^{C1045Y}$ (Night) | 24.75 ± 0.90 | 58 | |
| ElavGal4>UASdNf1-GRD; $Nf1^{C1045Y}$ (Night) | 25.32 ± 0.66 | 57 | >0.9999 |

| | | | |
|---|---|---|---|
| ElavGal4>UASdNf1-GRD-RP; $Nf1^{C1045Y}$ (Night) | 22.72 ± 0.76 | 90 | 0.3483 |
| **Fig. 8F** Kruskal Wallis $_{(3)}$ = 70.70 **p<0.0001** | | | |
| ElavGal4/+ (Day) | 22.45 ± 0.92 | 53 | |
| ElavGal4; $Nf1^{C1045Y}$ (Day) | 16.25 ± 1.3 | 58 | **<0.0001** |
| ElavGal4>UASdNf1-GRD; $Nf1^{C1045Y}$ (Day) | 19.07 ± 1.6 | 57 | 0.0205 |
| ElavGal4>UASdNf1-GRD-RP; $Nf1^{C1045Y}$ (Day) | 15.96 ± 2.1 | 90 | **<0.0001** |
| ElavGal4; $Nf1^{C1045Y}$ (Day) | 16.25 ± 1.3 | 58 | |
| ElavGal4>UASdNf1-GRD; $Nf1^{C1045Y}$ (Day) | 19.07 ± 1.6 | 57 | 0.0466 |
| ElavGal4>UASdNf1-GRD-RP; $Nf1^{C1045Y}$ (Day) | 15.96 ± 2.1 | 90 | 0.3284 |
| Kruskal Wallis $_{(3)}$ = 119.8 **p<0.0001** | | | |
| ElavGal4/+ (Night) | 34.51 ± 1.6 | 49 | |
| ElavGal4 $Nf1^{C1045Y}$ (Night) | 16.06 ± 2.2 | 58 | **<0.0001** |
| ElavGal4>UASdNf1-GRD; $Nf1^{C1045Y}$ (Night) | 15.51 ± 0.94 | 57 | **<0.0001** |
| ElavGal4>UASdNf1-GRD-RP; $Nf1^{C1045Y}$ (Night) | 11.58 ± 0.54 | 90 | **<0.0001** |
| ElavGal4; $Nf1^{C1045Y}$ (Night) | 16.06 ± 2.2 | 58 | |
| ElavGal4>UASdNf1-GRD; $Nf1^{C1045Y}$ (Night) | 15.51 ± 0.94 | 57 | 0.6779 |
| ElavGal4>UASdNf1-GRD-RP; $Nf1^{C1045Y}$ (Night) | 11.58 ± 0.54 | 90 | 0.0479 |
| Mann-Whitney (U)= 413 **p<0.0001** | | | |
| ElavGal4/+ (Day) | 22.45 ± 0.92 | 53 | |
| ElavGal4/+ (Night) | 34.51 ± 1.6 | 49 | **<0.0001** |
| Mann-Whitney (U)= 1172 p=0.0101 | | | |
| ElavGal4>UASdNf1-GRD; $Nf1^{C1045Y}$ (Day) | 19.07 ± 1.6 | 57 | |
| ElavGal4>UASdNf1-GRD; $Nf1^{C1045Y}$ (Night) | 15.51 ± 0.94 | 57 | 0.0101 |
| **Fig. 8G** Kruskal Wallis $_{(2)}$ = 93.60 **p<0.0001** | | | |
| ChaGal4/+ (Day) | 368.7 ± 10 | 95 | |
| UASdNf1; $Nf1^{C1045Y}$ (Day) | 234.9 ± 8.9 | 97 | **<0.0001** |
| ChaGal4/+ >UASdNf1; $Nf1^{C1045Y}$ (Day) | 306.4 ± 5.9 | 148 | **<0.0001** |
| ChaGal4/+ >UASdNf1; $Nf1^{C1045Y}$ (Day) | 306.4 ± 5.9 | 148 | |
| UASdNf1; $Nf1^{C1045Y}$ (Day) | 234.9 ± 8.9 | 97 | **<0.0001** |
| Kruskal Wallis $_{(2)}$ = 210.8 **p<0.0001** | | | |
| ChaGal4/+ (Night) | 561.8 ± 8.1 | 95 | |

| | | | |
|---|---|---|---|
| UASdNf1; Nf1$^{C1045Y}$ (Night) | 294.2 ± 9.6 | 97 | **<0.0001** |
| ChaGal4>UASdNf1; Nf1$^{C1045Y}$ (Night) | 470.9 ± 5.6 | 148 | **<0.0001** |
| ChaGal4>UASdNf1; Nf1$^{C1045Y}$ (Night) | 470.9 ± 5.6 | 148 | |
| UASdNf1; Nf1$^{C1045Y}$ (Night) | 294.2 ± 9.6 | 97 | **<0.0001** |
| Mann-Whitney (U)= 651 **p<0.0001** | | | |
| ChaGal4/+ (Day) | 368.7 ± 10 | 95 | |
| ChaGal4/+ (Night) | 561.8 ± 8.1 | 95 | **<0.0001** |
| Mann-Whitney (U)= 1206 **p<0.0001** | | | |
| ChaGal4/+ >UASdNf1; Nf1$^{C1045Y}$ (Day) | 306.4 ± 5.9 | 148 | |
| ChaGal4>UASdNf1; Nf1$^{C1045Y}$ (Night) | 470.9 ± 5.6 | 148 | **<0.0001** |
| **Fig. 8H** Kruskal Wallis $_{(2)}$ = 12.35 **p=0.0021** | | | |
| ChaGal4/+ (Day) | 21.71 ± 0.49 | 95 | |
| UASdNf1; Nf1$^{C1045Y}$ (Day) | 23.75 ± 0.56 | 97 | **0.0031** |
| ChaGal4>UASdNf1; Nf1$^{C1045Y}$ (Day) | 22.29 ± 0.39 | 148 | >0.9999 |
| ChaGal4>UASdNf1; Nf1$^{C1045Y}$ (Day) | 22.29 ± 0.39 | 148 | |
| UASdNf1; Nf1$^{C1045Y}$ (Day) | 23.75 ± 0.56 | 97 | 0.0132 |
| Kruskal Wallis $_{(2)}$ = 145.8 **p<0.0001** | | | |
| ChaGal4/+ (Night) | 15.20 ± 0.54 | 95 | |
| UASdNf1; Nf1$^{C1045Y}$ (Night) | 27.78 ± 0.66 | 97 | **<0.0001** |
| ChaGal4>UASdNf1; Nf1$^{C1045Y}$ (Night) | 21.98 ± 0.42 | 148 | **<0.0001** |
| ChaGal4>UASdNf1; Nf1$^{C1045Y}$ (Night) | 21.98 ± 0.42 | 148 | |
| UASdNf1; Nf1$^{C1045Y}$ (Night) | 27.78 ± 0.66 | 97 | **<0.0001** |
| **Fig. 8I** Kruskal Wallis $_{(2)}$ = 153.0 **p<0.0001** | | | |
| ChaGal4/+ (Day) | 17.47 ± 0.58 | 95 | |
| UASdNf1; Nf1$^{C1045Y}$ (Day) | 10.27 ± 0.71 | 97 | **<0.0001** |
| ChaGal4>UASdNf1; Nf1$^{C1045Y}$ (Day) | 14.11 ± 0.31 | 148 | **<0.0001** |
| ChaGal4>UASdNf1; Nf1$^{C1045Y}$ (Day) | 14.11 ± 0.31 | 148 | |
| UASdNf1; Nf1$^{C1045Y}$ (Day) | 10.27 ± 0.71 | 97 | **<0.0001** |
| Kruskal Wallis $_{(2)}$ = 235.5 **p<0.0001** | | | |
| ChaGal4/+ (Night) | 41.06 ± 1.7 | 93 | |
| UASdNf1; Nf1$^{C1045Y}$ (Night) | 10.95 ± 0.55 | 97 | **<0.0001** |
| ChaGal4>UASdNf1; Nf1$^{C1045Y}$ (Night) | 22.76 ± 0.57 | 148 | **<0.0001** |
| ChaGal4>UASdNf1; Nf1$^{C1045Y}$ (Night) | 22.76 ± 0.57 | 148 | |
| UASdNf1; Nf1$^{C1045Y}$ (Night) | 10.95 ± 0.55 | 97 | **<0.0001** |
| Mann-Whitney (U)= 439 **p<0.0001** | | | |

| | | | |
|---|---|---|---|
| ChaGal4/+ (Day) | 17.47 ± 0.58 | 96 | |
| ChaGal4/+ (Night) | 41.06 ± 1.7 | 93 | **<0.0001** |
| Mann-Whitney (U)= 2246 **p<0.0001** | | | |
| ChaGal4>UASdNf1; $Nf1^{C1045Y}$ (Day) | 14.11 ± 0.31 | 148 | |
| ChaGal4>UASdNf1; $Nf1^{C1045Y}$ (Night) | 22.76 ± 0.57 | 148 | **<0.0001** |
| **Fig. 8J** Kruskal Wallis $_{(3)}$ = 147.6 **p<0.0001** | | | |
| ChaGal4/+ (Day) | 404.8 ± 11 | 71 | |
| ChaGal4; $Nf1^{C1045Y}$ (Day) | 234.8 ± 15 | 33 | **<0.0001** |
| ChaGal4>UASdNf1-GRD; $Nf1^{C1045Y}$ (Day) | 362.7 ± 14 | 64 | 0.4120 |
| ChaGal4>UASdNf1-GRD-RP; $Nf1^{C1045Y}$ (Day) | 125.8 ± 12 | 79 | **<0.0001** |
| ChaGal4; $Nf1^{C1045Y}$ (Day) | 234.8 ± 15 | 33 | |
| ChaGal4>UASdNf1-GRD; $Nf1^{C1045Y}$ (Day) | 362.7 ± 14 | 64 | **0.0056** |
| ChaGal4>UASdNf1-GRD-RP; $Nf1^{C1045Y}$ (Day) | 125.8 ± 12 | 79 | **0.0007** |
| Kruskal Wallis $_{(3)}$ = 179.0 **p<0.0001** | | | |
| ChaGal4/+ (Night) | 506.2 ± 6.7 | 71 | |
| ChaGal4; $Nf1^{C1045Y}$ (Night) | 267.3 ± 18 | 33 | **<0.0001** |
| ChaGal4>UASdNf1-GRD; $Nf1^{C1045Y}$ (Night) | 428.1 ± 9.3 | 64 | **0.0006** |
| ChaGal4>UASdNf1-GRD-RP; $Nf1^{C1045Y}$ (Night) | 180.1 ± 13 | 79 | **<0.0001** |
| ChaGal4; $Nf1^{C1045Y}$ (Night) | 267.3 ± 18 | 33 | |
| ChaGal4>UASdNf1-GRD; $Nf1^{C1045Y}$ (Night) | 428.1 ± 9.3 | 64 | **<0.0001** |
| ChaGal4>UASdNf1-GRD-RP; $Nf1^{C1045Y}$ (Night) | 180.1 ± 13 | 79 | 0.2704 |
| Mann-Whitney (U)= 902 **p<0.0001** | | | |
| ChaGal4/+ (Day) | 404.8 ± 11 | 71 | |
| ChaGal4/+ (Night) | 506.2 ± 6.7 | 71 | **<0.0001** |
| Mann-Whitney (U)= 1218 **p<0.0001** | | | |
| ChaGal4>UASdNf1-GRD; $Nf1^{C1045Y}$ (Day) | 362.7 ± 14 | 64 | |
| ChaGal4>UASdNf1-GRD; $Nf1^{C1045Y}$ (Night) | 428.1 ± 9.3 | 64 | **<0.0001** |
| **Fig. 8K** Kruskal Wallis $_{(3)}$ = 120.2 **p<0.0001** | | | |
| ChaGal4/+ (Day) | 21.32 ± 0.64 | 71 | |
| ChaGal4; $Nf1^{C1045Y}$ (Day) | 22.76 ± 1.2 | 33 | >0.9999 |
| ChaGal4>UASdNf1-GRD; $Nf1^{C1045Y}$ (Day) | 26.25 ± 0.78 | 64 | **0.0036** |

| | | | |
|---|---|---|---|
| ChaGal4>UASdNf1-GRD-RP; Nf1$^{C1045Y}$ (Day) | 11.78 ± 0.68 | 79 | **<0.0001** |
| ChaGal4; Nf1$^{C1045Y}$ (Day) | 22.76 ± 1.2 | 33 | |
| ChaGal4>UASdNf1-GRD; Nf1$^{C1045Y}$ (Day) | 26.25 ± 0.78 | 64 | 0.3498 |
| ChaGal4>UASdNf1-GRD-RP; Nf1$^{C1045Y}$ (Day) | 11.78 ± 0.68 | 79 | **<0.0001** |
| Kruskal Wallis $_{(3)}$ = 44.72 **p<0.0001** | | | |
| ChaGal4/+ (Night) | 14.87 ± 0.55 | 71 | |
| ChaGal4; Nf1$^{C1045Y}$ (Night) | 23.12 ± 1.1 | 33 | **<0.0001** |
| ChaGal4>UASdNf1-GRD; Nf1$^{C1045Y}$ (Night) | 19.25 ± 0.57 | 64 | **<0.0001** |
| ChaGal4>UASdNf1-GRD-RP; Nf1$^{C1045Y}$ (Night) | 16.47 ± 0.90 | 79 | 0.1935 |
| ChaGal4; Nf1$^{C1045Y}$ (Night) | 23.12 ± 1.1 | 33 | |
| ChaGal4>UASdNf1-GRD; Nf1$^{C1045Y}$ (Night) | 19.25 ± 0.57 | 64 | 0.0785 |
| ChaGal4>UASdNf1-GRD-RP; Nf1$^{C1045Y}$ (Night) | 16.47 ± 0.90 | 79 | **<0.0001** |
| **Fig.8L** Kruskal Wallis $_{(3)}$ =118.2 **p<0.0001** | | | |
| ChaGal4/+ (Day) | 19.27 ± 0.60 | 70 | |
| ChaGal4; Nf1$^{C1045Y}$ (Day) | 10.30 ± 0.47 | 33 | **<0.0001** |
| ChaGal4>UASdNf1-GRD; Nf1$^{C1045Y}$ (Day) | 14.81 ± 0.90 | 64 | **0.0001** |
| ChaGal4>UASdNf1-GRD-RP; Nf1$^{C1045Y}$ (Day) | 10.05 ± 0.60 | 79 | **<0.0001** |
| ChaGal4; Nf1$^{C1045Y}$ (Day) | 10.30 ± 0.47 | 33 | |
| ChaGal4>UASdNf1-GRD; Nf1$^{C1045Y}$ (Day) | 14.81 ± 0.90 | 64 | **0.0056** |
| ChaGal4>UASdNf1-GRD-RP; Nf1$^{C1045Y}$ (Day) | 10.05 ± 0.60 | 79 | >0.9999 |
| Kruskal Wallis $_{(3)}$ =183.2 **p<0.0001** | | | |
| ChaGal4/+ (Night) | 34.06 ± 1.1 | 66 | |
| ChaGal4; Nf1$^{C1045Y}$ (Night) | 11.42 ± 0.57 | 33 | **<0.0001** |
| ChaGal4>UASdNf1-GRD; Nf1$^{C1045Y}$ (Night) | 23.96 ± 1.2 | 64 | **0.0014** |
| ChaGal4>UASdNf1-GRD-RP; Nf1$^{C1045Y}$ (Night) | 10.51 ± 0.37 | 78 | **<0.0001** |
| ChaGal4; Nf1$^{C1045Y}$ (Night) | 11.42 ± 0.57 | 33 | |
| ChaGal4>UASdNf1-GRD; Nf1$^{C1045Y}$ (Night) | 23.96 ± 1.2 | 64 | **<0.0001** |
| ChaGal4>UASdNf1-GRD-RP; Nf1$^{C1045Y}$ (Night) | 10.51 ± 0.37 | 78 | >0.9999 |
| Mann-Whitney (U)= 320 **p<0.0001** | | | |
| ChaGal4/+ (Day) | 19.27 ± 0.60 | 70 | |
| ChaGal4/+ (Night) | 34.06 ± 1.1 | 66 | **<0.0001** |
| Mann-Whitney (U)= 583 **p<0.0001** | | | |

| | | | |
|---|---|---|---|
| ChaGal4>UASdNf1-GRD; $Nf1^{C1045Y}$ (Day) | 14.81 ± 0.90 | 64 | |
| ChaGal4>UASdNf1-GRD; $Nf1^{C1045Y}$ (Night) | 23.96 ± 1.2 | 64 | **<0.0001** |
| **Fig. 9A** ANOVA $F_{(2, 169)}$ = 7.810 **p=0.0006** | | | |
| ElavGal4/+ (Day) | 325.5 ± 14 | 62 | |
| UASdNf1; $Nf1^{R1861C}$ (Day) | 264.5 ± 10 | 48 | **0.0022** |
| ElavGal4>UASdNf1; $Nf1^{R1861C}$ (Day) | 266.5 ± 12 | 62 | **0.0022** |
| ElavGal4>UASdNf1; $Nf1^{R1861C}$ (Day) | 266.5 ± 12 | 62 | |
| UASdNf1; $Nf1^{R1861C}$ (Day) | 264.5 ± 10 | 48 | 0.9129 |
| ANOVA $F_{(2, 169)}$ = 124.5 **p<0.0001** | | | |
| ElavGal4/+ (Night) | 532.3 ± 8.5 | 62 | |
| UASdNf1; $Nf1^{R1861C}$ (Night) | 284.3 ± 12 | 48 | **<0.0001** |
| ElavGal4>UASdNf1; $Nf1^{R1861C}$ (Night) | 440.5 ± 12 | 62 | **<0.0001** |
| ElavGal4>UASdNf1; $Nf1^{R1861C}$ (Night) | 440.5 ± 12 | 62 | |
| UASdNf1; $Nf1^{R1861C}$ (Night) | 284.3 ± 12 | 48 | **<0.0001** |
| t=12.39, df=98.96 **p<0.0001** | | | |
| ElavGal4/+ (Day) | 325.5 ± 14 | 62 | |
| ElavGal4/+ (Night) | 532.3 ± 8.5 | 62 | **<0.0001** |
| t=10.30, df= 121.9 **p <0.0001** | | | |
| ElavGal4>UASdNf1; $Nf1^{R1861C}$ (Day) | 266.5 ± 12 | 62 | |
| ElavGal4>UASdNf1; $Nf1^{R1861C}$ (Night) | 440.5 ± 12 | 62 | **0.0002** |
| **Fig. 9B** ANOVA $F_{(2, 169)}$ = 43.10 **p<0.0001** | | | |
| ElavGal4/+ (Day) | 20.85 ± 0.58 | 62 | |
| UASdNf1; $Nf1^{R1861C}$ (Day) | 22.10 ± 0.60 | 48 | 0.1612 |
| ElavGal4>UASdNf1; $Nf1^{R1861C}$ (Day) | 14.67 ± 0.63 | 62 | **<0.0001** |
| ElavGal4>UASdNf1; $Nf1^{R1861C}$ (Day) | 14.67 ± 0.63 | 62 | |
| UASdNf1; $Nf1^{R1861C}$ (Day) | 22.10 ± 0.60 | 48 | **<0.0001** |
| ANOVA $F_{(2, 169)}$ = 8.208 **p=0.0004** | | | |
| ElavGal4/+ (Night) | 18.47 ± 0.69 | 62 | |
| UASdNf1; $Nf1^{R1861C}$ (Night) | 21.97 ± 0.70 | 48 | **0.0008** |
| ElavGal4>UASdNf1; $Nf1^{R1861C}$ (Night) | 18.73 ± 0.55 | 62 | 0.7629 |
| ElavGal4>UASdNf1; $Nf1^{R1861C}$ (Night) | 18.73 ± 0.55 | 62 | |
| UASdNf1; $Nf1^{R1861C}$ (Night) | 21.97 ± 0.70 | 48 | **0.0015** |
| **Fig. 9C** Kruskal Wallis $_{(2)}$ = 38.02 **p<0.0001** | | | |
| ElavGal4/+ (Day) | 16.14 ± 0.92 | 62 | |

| | | | |
|---|---|---|---|
| UASdNf1; Nf1$^{R1861C}$ (Day) | 11.93 ± 0.33 | 48 | **0.0004** |
| ElavGal4>UASdNf1; Nf1$^{R1861C}$ (Day) | 19.55 ± 1.2 | 62 | 0.0359 |
| ElavGal4>UASdNf1; Nf1$^{R1861C}$ (Day) | 19.55 ± 1.2 | 62 | |
| UASdNf1; Nf1$^{R1861C}$ (Day) | 11.93 ± 0.33 | 48 | **<0.0001** |
| Kruskal Wallis $_{(2)}$ = 95.13 **p<0.0001** | | | |
| ElavGal4/+ (Night) | 30.73 ± 1.3 | 61 | |
| UASdNf1; Nf1$^{R1861C}$ (Night) | 12.97 ± 0.41 | 48 | **<0.0001** |
| ElavGal4>UASdNf1; Nf1$^{R1861C}$ (Night) | 24.74 ± 0.99 | 62 | 0.0273 |
| ElavGal4>UASdNf1; Nf1$^{R1861C}$ (Night) | 24.74 ± 0.99 | 62 | |
| UASdNf1; Nf1$^{R1861C}$ (Night) | 12.97 ± 0.41 | 48 | **<0.0001** |
| Mann-Whitney (U)=308 **p<0.0001** | | | |
| ElavGal4/+ (Day) | 16.14 ± 0.92 | 61 | |
| ElavGal4/+ (Night) | 30.73 ± 1.3 | 61 | **<0.0001** |
| Mann-Whitney (U)= 1087 **p<0.0001** | | | |
| ElavGal4>UASdNf1; Nf1$^{R1861C}$ (Day) | 19.55 ± 1.2 | 62 | |
| ElavGal4>UASdNf1; Nf1$^{R1861C}$ (Night) | 24.74 ± 0.99 | 62 | **<0.0001** |
| **Fig. 9D** ANOVA F $_{(3,316)}$ = 35.56 **p<0.0001** | | | |
| ElavGal4/+ (Day) | 393.5 ± 13 | 88 | |
| UASdNf1-GRD; Nf1$^{R1861C}$ (Day) | 329.3 ± 12 | 59 | **0.0015** |
| ElavGal4>UASdNf1-GRD; Nf1$^{R1861C}$ (Day) | 382.0 ± 11 | 74 | 0.5021 |
| ElavGal4>UASdNf1-GRD-RP; Nf1$^{R1861C}$ (Day) | 245.3 ± 12 | 99 | **<0.0001** |
| UASdNf1-GRD; Nf1$^{R1861C}$ (Day) | 329.3 ± 12 | 59 | |
| ElavGal4>UASdNf1-GRD; Nf1$^{R1861C}$ (Day) | 382.0 ± 11 | 74 | 0.0116 |
| ElavGal4>UASdNf1-GRD-RP; Nf1$^{R1861C}$ (Day) | 245.3 ± 12 | 99 | **<0.0001** |
| Kruskal Wallis $_{(3)}$ = 177.5 **p<0.0001** | | | |
| ElavGal4/+ (Night) | 567.6 ± 8.5 | 88 | |
| UASdNf1-GRD; Nf1$^{R1861C}$ (Night) | 296.0 ± 11 | 59 | **<0.0001** |
| ElavGal4>UASdNf1-GRD; Nf1$^{R1861C}$ (Night) | 399.0 ± 13 | 74 | **<0.0001** |
| ElavGal4>UASdNf1-GRD-RP; Nf1$^{R1861C}$ (Night) | 361.1 ± 9.7 | 99 | **<0.0001** |
| UASdNf1-GRD; Nf1$^{R1861C}$ (Night) | 296.0 ± 11 | 59 | |

| | | | |
|---|---|---|---|
| ElavGal4>UASdNf1- GRD; Nf1$^{R1861C}$ (Night) | 399.0 ± 13 | 74 | **<0.0001** |
| ElavGal4>UASdNf1-GRD-RP; Nf1$^{R1861C}$ (Night) | 361.1 ± 9.7 | 99 | 0.0141 |
| Mann-Whitney (U)= 958.8 **p<0.0001** | | | |
| ElavGal4/+ (Day) | 393.5 ± 13 | 88 | |
| ElavGal4/+ (Night) | 567.6 ± 8.5 | 88 | **<0.0001** |
| Mann-Whitney (U)= 2365 p=0.1527 | | | |
| ElavGal4>UASdNf1-GRD; Nf1$^{R1861C}$ (Day) | 382.0 ± 11 | 74 | |
| ElavGal4>UASdNf1-GRD; Nf1$^{R1861C}$ (Night) | 399.0 ± 13 | 74 | 0.1527 |
| **Fig. 9E** ANOVA $F_{(3,316)}$ = 22.21 **p<0.0001** | | | |
| ElavGal4/+ (Day) | 18.38 ± 0.56 | 88 | |
| UASdNf1-GRD; Nf1$^{R1861C}$ (Day) | 23.87 ± 0.52 | 59 | **<0.0001** |
| ElavGal4>UASdNf1-GRD; Nf1$^{R1861C}$ (Day) | 16.36 ± 0.64 | 74 | 0.0389 |
| ElavGal4>UASdNf1-GRD-RP; Nf1$^{R1861C}$ (Day) | 18.45 ± 0.63 | 99 | 0.9258 |
| UASdNf1-GRD; Nf1$^{R1861C}$ (Day) | 23.87 ± 0.52 | 89 | |
| ElavGal4>UASdNf1-GRD; Nf1$^{R1861C}$ (Day) | 16.36 ± 0.64 | 74 | **<0.0001** |
| ElavGal4>UASdNf1-GRD-RP; Nf1$^{R1861C}$ (Day) | 18.45 ± 0.63 | 99 | **<0.0001** |
| ANOVA $F_{(3,316)}$ =31.94 **p<0.0001** | | | |
| ElavGal4/+ (Night) | 14.76 ± 0.51 | 88 | |
| UASdNf1-GRD; Nf1$^{R1861C}$ (Night) | 21.22 ± 0.64 | 59 | **<0.0001** |
| ElavGal4>UASdNf1-GRD; Nf1$^{R1861C}$ (Night) | 19.70 ± 0.64 | 74 | **<0.0001** |
| ElavGal4>UASdNf1-GRD-RP; Nf1$^{R1861C}$ (Night) | 21.60 ± 0.54 | 99 | **<0.0001** |
| UASdNf1-GRD; Nf1$^{R1861C}$ (Night) | 21.22 ± 0.64 | 59 | |
| ElavGal4>UASdNf1-GRD; Nf1$^{R1861C}$ (Night) | 19.70 ± 0.64 | 74 | 0.1790 |
| ElavGal4>UASdNf1-GRD-RP; Nf1$^{R1861C}$ (Night) | 21.60 ± 0.54 | 99 | 0.6593 |
| **Fig. 9F** Kruskal Wallis $_{(3)}$ = 122.6 **p<0.0001** | | | |
| ElavGal4/+ (Day) | 23.44 ± 1.6 | 88 | |
| UASdNf1-GRD; Nf1$^{R1861C}$ (Day) | 13.77 ± 0.45 | 59 | **<0.0001** |
| ElavGal4>UASdNf1-GRD; Nf1$^{R1861C}$ (Day) | 27.69 ± 2.2 | 74 | >0.9999 |
| ElavGal4>UASdNf1-GRD-RP; Nf1$^{R1861C}$ (Day) | 13.49 ± 0.67 | 98 | **<0.0001** |
| UASdNf1-GRD; Nf1$^{R1861C}$ (Day) | 13.77 ± 0.45 | 59 | |
| ElavGal4>UASdNf1-GRD; Nf1$^{R1861C}$ (Day) | 27.69 ± 2.2 | 74 | **<0.0001** |
| ElavGal4>UASdNf1-GRD-RP; Nf1$^{R1861C}$ (Day) | 13.49 ± 0.67 | 98 | >0.9999 |
| Kruskal Wallis $_{(3)}$ = 168.7 **p<0.0001** | | | |
| ElavGal4/+ (Night) | 39.10 ± 1.5 | 82 | |

| | | | |
|---|---|---|---|
| UASdNf1-GRD; $Nf1^{R1861C}$ (Night) | 14.14 ± 0.51 | 59 | **<0.0001** |
| ElavGal4>UASdNf1-GRD; $Nf1^{R1861C}$ Night) | 21.58 ± 0.99 | 74 | **<0.0001** |
| ElavGal4>UASdNf1-GRD-RP; $Nf1^{R1861C}$ (Night) | 17.57 ± 0.71 | 99 | **<0.0001** |
| UASdNf1-GRD; $Nf1^{R1861C}$ (Night) | 14.14 ± 0.51 | 59 | |
| ElavGal4>UASdNf1-GRD; $Nf1^{R1861C}$ (Night) | 21.58 ± 0.99 | 74 | **<0.0001** |
| ElavGal4>UASdNf1-GRD-RP; $Nf1^{R1861C}$ (Night) | 17.57 ± 0.71 | 99 | 0.0360 |
| Mann-Whitney (U)=949 **p<0.0001** | | | |
| ElavGal4/+ (Day) | 23.44 ± 1.6 | 88 | |
| ElavGal4/+ (Night) | 39.10 ± 1.5 | 88 | **<0.0001** |
| Mann-Whitney (U)=2240 p=0.0559 | | | |
| ElavGal4>UASdNf1-GRD; $Nf1^{R1861C}$ (Day) | 27.69 ± 2.2 | 74 | |
| ElavGal4>UASdNf1-GRD; $Nf1^{R1861C}$ (Night) | 21.58 ± 0.99 | 74 | 0.0559 |
| **Fig. 9G** ANOVA $F_{(2, 399)}$ =13.21 **p < 0.0001** | | | |
| ChaGal4/+ (Day) | 401.3 ± 8.2 | 105 | |
| UASdNf1; $Nf1^{R1861C}$ (Day) | 343.9 ± 6.5 | 118 | **<0.0001** |
| ChaGal4>UASdNf1;$Nf1^{R1861C}$ (Day) | 377.9 ± 6.9 | 179 | 0.0241 |
| ChaGal4>UASdNf1; $Nf1^{R1861C}$ (Day) | 377.9 ± 6.9 | 179 | |
| UASdNf1; $Nf1^{R1861C}$ (Day) | 343.9 ± 6.5 | 118 | **0.0015** |
| Kruskal Wallis $_{(2)}$ = 93.88 **p<0.0001** | | | |
| ChaGal4/+ (Night) | 504.8 ± 5.5 | 105 | |
| UASdNf1; $Nf1^{R1861C}$ Night) | 388.1 ± 9.0 | 118 | **<0.0001** |
| ChaGal4>UASdNf1; $Nf1^{R1861C}$ (Night) | 483.2 ± 5.7 | 179 | **<0.0001** |
| ChaGal4>UASdNf1; $Nf1^{R1861C}$ (Night) | 483.2 ± 5.7 | 179 | |
| UASdNf1; $Nf1^{R1861C}$ (Night) | 388.1 ± 9.0 | 118 | **<0.0001** |
| Mann-Whitney (U) = 1674 **p<0.0001** | | | |
| ChaGal4/+ (Day) | 401.3 ± 8.2 | 105 | |
| ChaGal4/+ (Night) | 504.8 ± 5.5 | 105 | **<0.0001** |
| Mann-Whitney (U) = 5917 **p<0.0001** | | | |
| ChaGal4>UASdNf1; $Nf1^{R1861C}$ (Day) | 377.9 ± 6.9 | 179 | |
| ChaGal4>UASdNf1; $Nf1^{R1861C}$ (Night) | 483.2 ± 5.7 | 179 | **<0.0001** |
| **Fig. 9H** Kruskal Wallis $_{(2)}$ = 108.3 **p<0.0001** | | | |

| | | | |
|---|---|---|---|
| ChaGal4/+ (Day) | 21.54 ± 0.49 | 105 | |
| UASdNf1; $Nf1^{R1861C}$ (Day) | 24.83 ± 0.41 | 118 | **<0.0001** |
| ChaGal4>UASdNf1; $Nf1^{R1861C}$ (Day) | 18.38 ± 0.36 | 179 | **<0.0001** |
| ChaGal4>UASdNf1; $Nf1^{R1861C}$ (Day) | 18.38 ± 0.36 | 179 | |
| UASdNf1; $Nf1^{R1861C}$ (Day) | 24.83 ± 0.41 | 118 | **<0.0001** |
| Kruskal Wallis $_{(2)}$ = 127.8 **p<0.0001** | | | |
| ChaGal4/+ (Night) | 14.41 ± 0.46 | 105 | |
| UASdNf1; $Nf1^{R1861C}$ (Night) | 22.56 ± 0.49 | 118 | **<0.0001** |
| ChaGal4>UASdNf1; $Nf1^{R1861C}$ (Night) | 15.95 ± 0.32 | 179 | 0.0658 |
| ChaGal4>UASdNf1; $Nf1^{R1861C}$ (Night) | 15.95 ± 0.32 | 179 | |
| UASdNf1; $Nf1^{R1861C}$ (Night) | 22.56 ± 0.49 | 118 | **<0.0001** |
| **Fig. 9I** Kruskal Wallis $_{(2)}$ =91.84 **p<0.0001** | | | |
| ChaGal4/+ (Day) | 20.65 ± 1.6 | 105 | |
| UASdNf1; $Nf1^{R1861C}$ (Day) | 14.30 ± 0.38 | 118 | **<0.0001** |
| ChaGal4>UASdNf1; $Nf1^{R1861C}$ (Day) | 22.32 ± 0.79 | 179 | 0.4667 |
| ChaGal4>UASdNf1; $Nf1^{R1861C}$ (Day) | 22.32 ± 0.79 | 179 | |
| UASdNf1; $Nf1^{R1861C}$ (Day) | 14.30 ± 0.38 | 118 | **<0.0001** |
| Kruskal Wallis $_{(2)}$ =149.7 **p<0.0001** | | | |
| ChaGal4/+ (Night) | 38.39 ± 1.4 | 104 | |
| UASdNf1; $Nf1^{R1861C}$ (Night) | 18.67 ± 0.79 | 118 | **<0.0001** |
| ChaGal4>UASdNf1; $Nf1^{R1861C}$ (Night) | 33.09 ± 1.1 | 179 | 0.0127 |
| ChaGal4>UASdNf1; $Nf1^{R1861C}$ (Night) | 33.09 ± 1.1 | 179 | |
| UASdNf1; $Nf1^{R1861C}$ (Night) | 18.67 ± 0.79 | 118 | **<0.0001** |
| Mann-Whitney (U)=789 **p<0.0001** | | | |
| ChaGal4/+ (Day) | 20.65 ± 1.6 | 105 | |
| ChaGal4/+ (Night) | 38.39 ± 1.4 | 104 | **<0.0001** |
| Mann-Whitney (U)=6603 **p<0.0001** | | | |
| ChaGal4>UASdNf1; $Nf1^{R1861C}$ (Day) | 22.32 ± 0.79 | 179 | |
| ChaGal4>UASdNf1; $Nf1^{R1861C}$ (Night) | 33.09 ± 1.1 | 179 | **<0.0001** |
| **Fig. 9J** Kruskal Wallis $_{(3)}$ = 100.7 **p<0.0001** | | | |
| ChaGal4/+ (Day) | 353.5 ± 9.7 | 102 | |
| ChaGal4; $Nf1^{R1861C}$ (Day) | 306.0 ± 8.2 | 109 | **0.0034** |
| ChaGal4>UASdNf1-GRD; $Nf1^{R1861C}$ (Day) | 311.5 ± 7.1 | 214 | **0.0026** |
| ChaGal4>UASdNf1-GRD-RP; $Nf1^{R1861C}$ (Day) | 188.3 ± 9.1 | 62 | **<0.0001** |

| | | | |
|---|---|---|---|
| ChaGal4; Nf1$^{R1861C}$ (Day) | 306.0 ± 8.2 | 109 | |
| ChaGal4>UASdNf1-GRD; Nf1$^{R1861C}$ (Day) | 311.5 ± 7.1 | 214 | >0.9999 |
| ChaGal4>UASdNf1-GRD-RP; Nf1$^{R1861C}$ (Day) | 188.3 ± 9.1 | 62 | **<0.0001** |
| Kruskal Wallis $_{(3)}$ = 174.5 **p<0.0001** | | | |
| ChaGal4/+ (Night) | 557.6 ± 7.2 | 102 | |
| ChaGal4; Nf1$^{R1861C}$ (Night) | 350.2 ± 9.8 | 109 | **<0.0001** |
| ChaGal4>UASdNf1-GRD; Nf1$^{R1861C}$ (Night) | 457.1 ± 7.5 | 214 | **<0.0001** |
| ChaGal4>UASdNf1-GRD-RP; Nf1$^{R1861C}$ (Night) | 409.7 ± 13 | 62 | **<0.0001** |
| ChaGal4; Nf1$^{R1861C}$ (Night) | 350.2 ± 9.8 | 109 | |
| ChaGal4>UASdNf1-GRD; Nf1$^{R1861C}$ (Night) | 457.1 ± 7.5 | 214 | **<0.0001** |
| ChaGal4>UASdNf1-GRD-RP; Nf1$^{R1861C}$ (Night) | 409.7 ± 13 | 62 | 0.0194 |
| Mann-Whitney (U) = 433 **p<0.0001** | | | |
| ChaGal4/+ (Day) | 353.5 ± 9.7 | 102 | |
| ChaGal4/+ (Night) | 557.6 ± 7.2 | 102 | **<0.0001** |
| Mann-Whitney (U) = 7754 **p<0.0001** | | | |
| ChaGal4>UASdNf1-GRD; Nf1$^{R1861C}$ (Day) | 311.5 ± 7.1 | 214 | |
| ChaGal4>UASdNf1-GRD; Nf1$^{R1861C}$ (Night) | 457.1 ± 7.5 | 214 | **<0.0001** |
| **Fig. 9K** Kruskal Wallis $_{(3)}$ = 132.9 **p<0.0001** | | | |
| ChaGal4/+ (Day) | 20.26 ± 0.49 | 102 | |
| ChaGal4; Nf1$^{R1861C}$ (Day) | 23.12 ± 0.42 | 109 | **0.0008** |
| ChaGal4>UASdNf1-GRD; Nf1$^{R1861C}$ (Day) | 16.79 ± 0.33 | 214 | **<0.0001** |
| ChaGal4>UASdNf1-GRD-RP; Nf1$^{R1861C}$ (Day) | 15.43 ± 0.62 | 62 | **<0.0001** |
| ChaGal4; Nf1$^{R1861C}$ (Day) | 23.12 ± 0.42 | 109 | |
| ChaGal4>UASdNf1-GRD; Nf1$^{R1861C}$ (Day) | 16.79 ± 0.33 | 214 | **<0.0001** |
| ChaGal4>UASdNf1-GRD-RP; Nf1$^{R1861C}$ (Day) | 15.43 ± 0.62 | 62 | **<0.0001** |
| Kruskal Wallis $_{(3)}$ = 108.7 **p<0.0001** | | | |
| ChaGal4/+ (Night) | 15.62 ± 0.54 | 102 | |
| ChaGal4; Nf1$^{R1861C}$ (Night) | 24.13 ± 0.55 | 109 | **<0.0001** |
| ChaGal4>UASdNf1-GRD; Nf1$^{R1861C}$ (Night) | 20.17 ± 0.37 | 214 | **<0.0001** |
| ChaGal4>UASdNf1-GRD-RP; Nf1$^{R1861C}$ (Night) | 22.15 ± 0.61 | 62 | **<0.0001** |
| ChaGal4; Nf1$^{R1861C}$ (Night) | 24.13 ± 0.55 | 109 | |
| ChaGal4>UASdNf1-GRD; Nf1$^{R1861C}$ (Night) | 20.17 ± 0.37 | 214 | **<0.0001** |

| | | | |
|---|---|---|---|
| ChaGal4>UASdNf1-GRD-RP; $Nf1^{R1861C}$ (Night) | 22.15 ± 0.61 | 62 | 0.2889 |
| **Fig. 9L** Kruskal Wallis $_{(3)}$ =108.8 **p<0.0001** | | | |
| ChaGal4/+ (Day) | 18.64 ± 0.89 | 102 | |
| ChaGal4; $Nf1^{R1861C}$ (Day) | 13.81 ± 0.61 | 109 | **<0.0001** |
| ChaGal4>UASdNf1-GRD; $Nf1^{R1861C}$ (Day) | 20.36 ± 0.80 | 214 | >0.9999 |
| ChaGal4>UASdNf1-GRD-RP; $Nf1^{R1861C}$ (Day) | 12.24 ± 0.42 | 62 | **<0.0001** |
| ChaGal4; $Nf1^{R1861C}$ (Day) | 13.81 ± 0.61 | 109 | |
| ChaGal4>UASdNf1-GRD; $Nf1^{R1861C}$ (Day) | 20.36 ± 0.80 | 214 | **<0.0001** |
| ChaGal4>UASdNf1-GRD-RP; $Nf1^{R1861C}$ (Day) | 12.24 ± 0.42 | 62 | 0.8141 |
| Kruskal Wallis $_{(3)}$ =218.5 **p<0.0001** | | | |
| ChaGal4/+ (Night) | 42.38 ± 2.5 | 102 | |
| ChaGal4; $Nf1^{R1861C}$ (Night) | 15.24 ± 0.65 | 109 | **<0.0001** |
| ChaGal4>UASdNf1-GRD; $Nf1^{R1861C}$ (Night) | 28.25 ± 3.3 | 214 | **<0.0001** |
| ChaGal4>UASdNf1-GRD-RP; $Nf1^{R1861C}$ (Night) | 19.55 ± 1.0 | 62 | **<0.0001** |
| ChaGal4; $Nf1^{R1861C}$ (Night) | 15.24 ± 0.65 | 109 | |
| ChaGal4>UASdNf1-GRD; $Nf1^{R1861C}$ (Night) | 28.25 ± 3.3 | 214 | **<0.0001** |
| ChaGal4>UASdNf1-GRD-RP; $Nf1^{R1861C}$ (Night) | 19.55 ± 1.0 | 62 | **0.0013** |
| Mann-Whitney (U)= 778.5 **p<0.0001** | | | |
| ChaGal4/+ (Day) | 18.64 ± 0.89 | 102 | |
| ChaGal4/+ (Night) | 42.38 ± 2.5 | 102 | **<0.0001** |
| Mann-Whitney (U)= 15151 **p<0.0001** | | | |
| ChaGal4>UASdNf1-GRD; $Nf1^{R1861C}$ (Day) | 20.36 ± 0.80 | 214 | |
| ChaGal4>UASdNf1-GRD; $Nf1^{R1861C}$ Night) | 28.25 ± 3.3 | 214 | **<0.0001** |
| **SUPPLEMENTAL FIGURES** | | | |
| **Fig. S1 B** Kruskal Wallis $H_{(4)}$ = 131.0 **p<0.0001** | | | |
| $w^{1118}$ | 799.9 ± 26 | 57 | |
| $Nf1^{Null}$ | 208.9 ± 17 | 37 | **<0.0001** |
| $Nf1^{\Delta 216}$ | 206.9 ± 15 | 54 | **<0.0001** |
| $Nf1^{Null/+}$ | 904.0 ± 44 | 10 | >0.9999 |
| $Nf1^{Null/\Delta 216}$ | 232.3 ± 19 | 37 | **<0.0001** |
| **Fig. S1 C** Kruskal Wallis $H_{(4)}$ = 92.15 **p<0.0001** | | | |

| | | | |
|---|---|---|---|
| $w^{1118}$ (Day) | 291.2 ± 16 | 57 | |
| $Nf1^{Null}$ (Day) | 109.3 ± 10 | 37 | **<0.0001** |
| $Nf1^{\Delta216}$ (Day) | 112.3 ± 12 | 56 | **<0.0001** |
| $Nf1^{Null/+}$ (Day) | 339.7 ± 31 | 11 | >0.9999 |
| $Nf1^{Null/\Delta216}$ (Day) | 115.4 ± 14 | 37 | **<0.0001** |
| Kruskal Wallis $H_{(4)}$ = 127.2 **p<0.0001** | | | |
| $w^{1118}$ (Night) | 508.5 ± 12 | 57 | |
| $Nf1^{Null}$ (Night) | 99.46 ± 10 | 37 | **<0.0001** |
| $Nf1^{\Delta216}$ (Night) | 124.3 ± 15 | 56 | **<0.0001** |
| $Nf1^{Null/+}$ (Night) | 563.8 ± 19 | 11 | **>0.9999** |
| $Nf1^{Null/\Delta216}$ (Night) | 116.8 ± 11 | 37 | **<0.0001** |
| Mann-Whitney (U)= 296.5 **p<0.0001** | | | |
| $w^{1118}$ (Day) | 291.2 ± 16 | 57 | |
| $w^{1118}$ (Night) | 508.5 ± 12 | 57 | **<0.0001** |
| Mann-Whitney (U)= 1 **p<0.0001** | | | |
| $Nf1^{Null/+}$ (Day) | 339.7 ± 31 | 11 | |
| $Nf1^{Null/+}$ (Night) | 563.8 ± 19 | 11 | **<0.0001** |
| Mann-Whitney (U)= 1501 p=0.6966 | | | |
| $Nf1^{\Delta216}$ (Day) | 112.3 ± 12 | 56 | |
| $Nf1^{\Delta216}$ (Night) | 124.3 ± 15 | 56 | 0.6966 |
| Mann-Whitney (U)= 643 p=0.6574 | | | |
| $Nf1^{Null/\Delta216}$ (Day) | 115.4 ± 14 | 37 | |
| $Nf1^{Null/\Delta216}$ (Night) | 116.8 ± 11 | 37 | 0.6574 |
| **Fig.S1 D** ANOVA $F_{(4,193)}$ = 27.87 **p<0.0001** | | | |
| $w^{1118}$ (Day) | 20.79 ± 0.64 | 57 | |
| $Nf1^{Null}$ (Day) | 13.45 ± 0.97 | 37 | **<0.0001** |
| $Nf1^{\Delta216}$ (Day) | 11.16 ± 0.77 | 56 | **<0.0001** |
| $Nf1^{Null/+}$ (Day) | 23.85 ± 1.0 | 11 | 0.1086 |
| $Nf1^{Null/\Delta216}$ (Day) | 13.59 ± 1.2 | 37 | **<0.0001** |
| ANOVA $F_{(4,193)}$ = 21.41 **p<0.0001** | | | |
| $w^{1118}$ (Night) | 21.91 ± 0.76 | 57 | |
| $Nf1^{Null}$ (Night) | 13.24 ± 1.1 | 37 | **<0.0001** |
| $Nf1^{\Delta216}$ (Night) | 12.18 ± 0.86 | 56 | **<0.0001** |
| $Nf1^{Null/+}$ (Night) | 19.35 ± 1.8 | 11 | 0.2193 |
| $Nf1^{Null/\Delta216}$ (Night) | 13.66 ± 1.1 | 37 | **<0.0001** |
| **Fig. S1 E** Kruskal Wallis $H_{(4)}$ = 90.54 **p<0.0001** | | | |
| $w^{1118}$ (Day) | 14.03 ± 0.77 | 57 | |
| $Nf1^{Null}$ (Day) | 7.858 ± 0.32 | 37 | **<0.0001** |
| $Nf1^{\Delta216}$ (Day) | 10.11 ± 1.4 | 55 | **<0.0001** |
| $Nf1^{Null/+}$ (Day) | 14.15 ± 1.2 | 11 | >0.9999 |
| $Nf1^{Null/\Delta216}$ (Day) | 7.843 ± 0.33 | 36 | **<0.0001** |

| Kruskal Wallis $H_{(4)}$ = 135.1 **p<0.0001** | | | |
|---|---|---|---|
| $w^{1118}$ (Night) | 25.93 ± 1.6 | 57 | |
| $Nf1^{Null}$ (Night) | 7.420 ± 0.58 | 37 | **<0.0001** |
| $Nf1^{\Delta216}$ (Night) | 9.444 ± 0.67 | 56 | **<0.0001** |
| $Nf1^{Null/+}$ (Night) | 33.58 ± 4.7 | 11 | >0.9999 |
| $Nf1^{Null/\Delta216}$ (Night) | 8.196 ± 0.23 | 37 | **<0.0001** |
| Mann-Whitney (U)= 435 **p<0.0001** | | | |
| $w^{1118}$ (Day) | 14.03 ± 0.77 | 57 | |
| $w^{1118}$ (Night) | 25.93 ± 1.6 | 57 | **<0.0001** |
| Mann-Whitney (U)= 4 **p<0.0001** | | | |
| $Nf1^{Null/+}$ (Day) | 14.15 ± 1.2 | 11 | |
| $Nf1^{Null/+}$ (Night) | 33.58 ± 4.7 | 11 | **<0.0001** |
| Mann-Whitney (U)= 1487 p=0.7544 | | | |
| $Nf1^{\Delta216}$ (Day) | 10.11 ± 1.4 | 55 | |
| $Nf1^{\Delta216}$ (Night) | 9.444 ± 0.67 | 56 | 0.7544 |
| Mann-Whitney (U)=515.5 p=0.0975 | | | |
| $Nf1^{Null/\Delta216}$ (Day) | 7.843 ± 0.33 | 36 | |
| $Nf1^{Null/\Delta216}$ (Night) | 8.196 ± 0.23 | 37 | 0.0975 |
| **Fig. S1 F** Kruskal Wallis $H_{(4)}$ = 72.65 **p<0.0001** | | | |
| $w^{1118}$ (Day) | 7504 ± 390 | 57 | |
| $Nf1^{Null}$ (Day) | 4987 ± 140 | 37 | **<0.0001** |
| $Nf1^{\Delta216}$ (Day) | 9682 ± 370 | 56 | **0.0016** |
| $Nf1^{Null/+}$ (Day) | 6071 ± 960 | 10 | 0.3905 |
| $Nf1^{Null/\Delta216}$ (Day) | 6422 ± 270 | 37 | 0.1823 |
| Kruskal Wallis $H_{(4)}$ =85.93 **p<0.0001** | | | |
| $w^{1118}$ (Night) | 3400 ± 270 | 57 | |
| $Nf1^{Null}$ (Night) | 5255 ± 230 | 37 | **0.0030** |
| $Nf1^{\Delta216}$ (Night) | 8042 ± 400 | 55 | **<0.0001** |
| $Nf1^{Null/+}$ (Night) | 2760 ± 510 | 10 | >0.9999 |
| $Nf1^{Null/\Delta216}$ (Night) | 6078 ± 340 | 37 | **<0.0001** |
| Mann-Whitney (U)= 462 **p<0.0001** | | | |
| $w^{1118}$ (Day) | 7504 ± 390 | 57 | |
| $w^{1118}$ (Night) | 3400 ± 270 | 57 | **<0.0001** |
| Mann-Whitney (U)= 16 **p=0.0089** | | | |
| $Nf1^{Null/+}$ (Day) | 6071 ± 960 | 10 | |
| $Nf1^{Null/+}$ (Night) | 2760 ± 510 | 10 | **0.0089** |

| | | | |
|---|---|---|---|
| Mann-Whitney (U)= 998 **p=0.0013** | | | |
| $Nf1^{\Delta216}$ (Day) | 9682 ± 370 | 56 | |
| $Nf1^{\Delta216}$ (Night) | 8042 ± 400 | 55 | **0.0013** |
| Mann-Whitney (U)= 601 p=0.3717 | | | |
| $Nf1^{Null/\Delta216}$ (Day) | 6422 ± 270 | 37 | |
| $Nf1^{Null/\Delta216}$ (Night) | 6078 ± 340 | 37 | 0.3717 |
| **Fig. S1G** Kruskal Wallis $H_{(4)}$ = 77.88 **p<0.0001** | | | |
| $w^{1118}$ (Day) | 6.364 ± 0.29 | 57 | |
| $Nf1^{Null}$ (Day) | 3.741 ± 0.096 | 37 | **<0.0001** |
| $Nf1^{\Delta216}$ (Day) | 7.230 ± 0.25 | 56 | 0.1603 |
| $Nf1^{Null/+}$ (Day) | 5.361 ± 0.70 | 10 | 0.4788 |
| $Nf1^{Null/\Delta216}$ (Day) | 4.819 ± 0.18 | 37 | **0.0021** |
| Kruskal Wallis $H_{(4)}$ = 54.25 **p<0.0001** | | | |
| $w^{1118}$ (Night) | 3.500 ± 0.24 | 57 | |
| $Nf1^{Null}$ (Night) | 3.904 ± 0.16 | 37 | >0.9999 |
| $Nf1^{\Delta216}$ (Night) | 5.984 ± 0.27 | 55 | **<0.0001** |
| $Nf1^{Null/+}$ (Night) | 3.058 ± 0.51 | 10 | >0.9999 |
| $Nf1^{Null/\Delta216}$ (Night) | 4.547 ± 0.23 | 37 | 0.0945 |
| Mann-Whitney (U)= 515 **p<0.0001** | | | |
| $w^{1118}$ (Day) | 6.364 ± 0.29 | 57 | |
| $w^{1118}$ (Night) | 3.500 ± 0.24 | 57 | **<0.0001** |
| Mann-Whitney (U)= 19 p=0.0185 | | | |
| $Nf1^{Null/+}$ (Day) | 5.361 ± 0.70 | 10 | |
| $Nf1^{Null/+}$ (Night) | 3.058 ± 0.51 | 10 | 0.0185 |
| Mann-Whitney (U)= 938 **p=0.0003** | | | |
| $Nf1^{\Delta216}$ (Day) | 7.230 ± 0.25 | 56 | |
| $Nf1^{\Delta216}$ (Night) | 5.984 ± 0.27 | 55 | **0.0003** |
| Mann-Whitney (U)=590 p=0.3115 | | | |
| $Nf1^{Null/\Delta216}$ (Day) | 4.819 ± 0.18 | 37 | |
| $Nf1^{Null/\Delta216}$ (Night) | 4.547 ± 0.23 | 37 | 0.3115 |
| Genotype | Mean ± SEM | n | p |
| **Fig. S2A** ANOVA $F_{(2,187)}$ = 32.65 **p<0.0001** | | | |
| ElavGal4>Dcr-2 | 749.7 ± 23 | 63 | |
| ElavGal4/+ | 897.5 ± 18 | 63 | **<0.0001** |
| Dcr-2/+ | 949.2 ± 12 | 64 | **<0.0001** |
| **Fig. S2B** Kruskal Wallis $_{(2)}$ =100.5 **p<0.0001** | | | |

| | | | |
|---|---|---|---|
| ElavGal4>Dcr-2 (Day) | 350.4 ± 10 | 63 | |
| ElavGal4/+ (Day) | 378.3 ± 11 | 63 | 0.2316 |
| Dcr-2/+ (Day) | 517.0 ± 6.1 | 64 | **<0.0001** |
| Kruskal Wallis $_{(2)}$ =43.05 **p<0.0001** | | | |
| ElavGal4>Dcr-2 (Night) | 413.5 ± 14 | 63 | |
| ElavGal4/+ (Night) | 516.7 ± 8.9 | 63 | **<0.0001** |
| Dcr-2/+ (Night) | 431.7 ± 9.7 | 64 | >0.9999 |
| **Fig. S2C** Kruskal Wallis $_{(2)}$ =21.49 **p<0.0001** | | | |
| ElavGal4>Dcr-2 (Day) | 19.65 ± 0.63 | 63 | |
| ElavGal4/+ (Day) | 22.66 ± 0.65 | 63 | **0.0069** |
| Dcr-2/+ (Day) | 23.79 ± 0.56 | 64 | **<0.0001** |
| Kruskal Wallis $_{(2)}$ =32.14 **p<0.0001** | | | |
| ElavGal4>Dcr-2 (Night) | 14.79 ± 0.57 | 63 | |
| ElavGal4/+ (Night) | 19.78 ± 0.61 | 63 | **<0.0001** |
| Dcr-2/+ (Night) | 16.91 ± 0.66 | 64 | 0.0461 |
| **Fig. S2D** Kruskal Wallis $_{(2)}$ = 35.36 **p<0.0001** | | | |
| ElavGal4>Dcr-2 (Day) | 17.25 ± 0.69 | 63 | |
| ElavGal4/+ (Day) | 17.44 ± 0.71 | 63 | >0.9999 |
| Dcr-2/+ (Day) | 22.84 ± 0.83 | 64 | **<0.0001** |
| Kruskal Wallis $_{(2)}$ = 1.996 **p=0.3686** | | | |
| ElavGal4>Dcr-2 (Night) | 29.74 ± 1.6 | 63 | |
| ElavGal4/+ (Night) | 27.04 ± 1.1 | 63 | 0.3159 |
| Dcr-2/+ (Night) | 27.72 ± 1.1 | 64 | 0.9023 |
| **Fig. S2E** Kruskal Wallis $_{(2)}$ = 80.09 **p<0.0001** | | | |
| ElavGal4>Dcr-2 (Day) | 7022 ± 350 | 63 | |
| ElavGal4/+ (Day) | 4037 ± 190 | 63 | **<0.0001** |
| Dcr-2/+ (Day) | 3074 ± 160 | 64 | **<0.0001** |
| Kruskal Wallis $_{(2)}$ = 57.50 **p<0.0001** | | | |
| ElavGal4>Dcr-2 (Night) | 8275 ± 530 | 63 | |
| ElavGal4/+ (Night) | 3674 ± 290 | 63 | **<0.0001** |
| Dcr-2/+ (Night) | 6238 ± 250 | 64 | 0.0850 |
| **Fig. S2F** Kruskal Wallis $_{(2)}$ = 73.66 **p<0.0001** | | | |
| ElavGal4>Dcr-2 (Day) | 6.240 ± 0.26 | 63 | |
| ElavGal4/+ (Day) | 3.734 ± 0.15 | 63 | **<0.0001** |
| Dcr-2/+ (Day) | 3.284 ± 0.16 | 64 | **<0.0001** |
| Kruskal Wallis $_{(2)}$ = 59.20 **p<0.0001** | | | |
| ElavGal4>Dcr-2 (Night) | 7.776 ± 0.41 | 63 | |
| ElavGal4/+ (Night) | 3.789 ± 0.26 | 63 | **<0.0001** |
| Dcr-2/+ (Night) | 6.095 ± 0.21 | 64 | 0.0402 |

| | | | |
|---|---|---|---|
| **Fig. S3A** Kruskal Wallis $_{(2)}$ = 33.31 **p<0.0001** | | | |
| ElavGal4>Nf1$^{RNAi}$ (Day) | 252.2 ± 17 | 30 | |
| ElavGal4/+ (Day) | 378.3 ± 11 | 64 | **0.0001** |
| Nf1$^{RNAi}$/+ (Day) | 387.5 ± 14 | 32 | **0.0001** |
| Kruskal Wallis $_{(2)}$ = 58.96 **p<0.0001** | | | |
| ElavGal4>Nf1$^{RNAi}$ (Night) | 347.9 ± 15 | 30 | |
| ElavGal4/+ (Night) | 509.2 ± 12 | 64 | **0.0001** |
| Nf1$^{RNAi}$/+ (Night) | 546.9 ± 5.6 | 32 | **0.0001** |
| **Fig. S3B** Kruskal Wallis $_{(2)}$ = 11.89 p=0.0026 | | | |
| ElavGal4>Nf1$^{RNAi}$ (Day) | 19.25 ± 0.98 | 30 | |
| ElavGal4/+ (Day) | 22.66 ± 0.65 | 64 | **0.0086** |
| Nf1$^{RNAi}$/+ (Day) | 19.81 ± 0.77 | 32 | 0.9999 |
| Kruskal Wallis $_{(2)}$ = 29.30 **p<0.0001** | | | |
| ElavGal4>Nf1$^{RNAi}$ (Night) | 20.60 ± 0.63 | 30 | |
| ElavGal4/+ (Night) | 19.78 ± 0.61 | 64 | 0.6199 |
| Nf1$^{RNAi}$ /+ (Night) | 14.72 ± 0.74 | 32 | **<0.0001** |
| **Fig. S3C** Kruskal Wallis $_{(2)}$ = 31.58 **p<0.0001** | | | |
| ElavGal4>Nf1$^{RNAi}$ (Day) | 13.84 ± 1.4 | 31 | |
| ElavGal4/+ (Day) | 17.44 ± 0.71 | 64 | **0.0002** |
| Nf1$^{RNAi}$ /+ (Day) | 20.03 ± 0.86 | 32 | **<0.0001** |
| Kruskal Wallis $_{(2)}$ = 69.14 **p<0.0001** | | | |
| ElavGal4>Nf1$^{RNAi}$ (Night) | 16.96 ± 0.68 | 31 | |
| ElavGal4/+ (Night) | 27.04 ± 1.1 | 64 | **<0.0001** |
| Nf1$^{RNAi}$ /+ (Night) | 41.22 ± 3.0 | 32 | **<0.0001** |
| **Fig. S3D** Kruskal Wallis $H_{(2)}$=19.39 **p<0.0001** | | | |
| LeoGal4>Nf1RNAi; Dcr-2 (Day) | 408.9 ± 27 | 38 | |
| LeoGal4/+ (Day) | 287.8 ± 29 | 24 | **0.0010** |
| Nf1RNAi; Dcr-2/+ (Day) | 433.1 ± 38 | 22 | 0.4807 |
| Kruskal Wallis $H_{(2)}$= 3.660 p=0.1604 | | | |
| LeoGal4>Nf1RNAi; Dcr-2(Night) | 550.4 ± 11 | 38 | |
| LeoGal4/+ (Night) | 561.1 ± 8.2 | 24 | >0.9999 |
| Nf1RNAi; Dcr-2/+ (Night) | 534.7 ± 13 | 22 | 0.1579 |
| **Fig. S3E** Kruskal Wallis $H_{(2)}$= 3.561 p=0.1685 | | | |
| LeoGal4>Nf1RNAi; Dcr-2 (Day) | 20.32 ± 1.4 | 38 | |

| | | | |
|---|---|---|---|
| LeoGal4/+ (Day) | 19.04 ± 2.0 | 24 | >0.9999 |
| Nf1RNAi; Dcr-2/+ (Day) | 23.05 ± 2.1 | 22 | 0.2593 |
| Kruskal Wallis $H_{(2)}$= 7.672 p=0.0216 | | | |
| LeoGal4>Nf1RNAi; Dcr-2(Night) | 18.07 ± 0.85 | 38 | |
| LeoGal4/+ (Night) | 19.63 ± 1.2 | 24 | 0.7128 |
| Nf1RNAi; Dcr-2/+ (Night) | 15.45 ± 0.81 | 22 | 0.0766 |
| **Fig. S3F** Kruskal Wallis $H_{(2)}$= 9.097 p=0.0106 | | | |
| LeoGal4>Nf1RNAi; Dcr-2 (Day) | 24.74 ± 4.5 | 35 | |
| LeoGal4/+ (Day) | 15.15 ± 0.95 | 21 | **0.0051** |
| Nf1RNAi; Dcr-2/+ (Day) | 18.61 ± 1.9 | 22 | 0.5739 |
| Kruskal Wallis $H_{(2)}$= 3.254 p=0.1966 | | | |
| LeoGal4>Nf1RNAi; Dcr-2(Night) | 33.90 ± 2.1 | 38 | |
| LeoGal4/+ (Night) | 32.04 ± 2.6 | 24 | 0.8999 |
| Nf1RNAi; Dcr-2/+ (Night) | 37.39 ± 3.0 | 22 | 0.4354 |
| **Fig. S3G** Kruskal Wallis $H_{(2)}$ =57.36 **p<0.0001** | | | |
| ElavGal4;MBG80>Nf1RNAi; Dcr-2(Day) | 161.6 ± 20 | 60 | |
| ElavGal4;MBG80/+ (Day) | 372.9 ± 19 | 43 | **<0.0001** |
| Nf1RNAi; Dcr-2/+ (Day) | 417.3 ± 11 | 45 | **<0.0001** |
| Kruskal Wallis $H_{(2)}$ =76.20 **p<0.0001** | | | |
| ElavGal4;MBG80>Nf1RNAi; Dcr-2(Night) | 92.33 ± 24 | 60 | |
| ElavGal4;MBG80/+ (Night) | 520.4 ± 12 | 43 | **<0.0001** |
| Nf1RNAi; Dcr-2/+ (Night) | 524.5 ± 9.8 | 45 | **<0.0001** |
| Mann-Whitney (U)= 275 **p<0.0001** | | | |
| ElavGal4;MBG80/+ (Day) | 372.9 ± 19 | 43 | |
| ElavGal4;MBG80/+ (Night) | 520.4 ± 12 | 43 | **<0.0001** |
| Mann-Whitney (U)= 283.5 **p<0.0001** | | | |
| Nf1RNAi; Dcr-2/+ (Day) | 417.3 ± 11 | 45 | |
| Nf1RNAi; Dcr-2/+ (Night) | 524.5 ± 9.8 | 45 | **<0.0001** |
| Mann-Whitney (U)= 784 **p<0.0001** | | | |
| ElavGal4;MBG80>Nf1RNAi Dcr-2(Day) | 161.6 ± 20 | 60 | |
| ElavGal4;MBG80>Nf1RNAi Dcr-2(Night) | 92.33 ± 24 | 60 | **<0.0001** |
| **Fig. S3H** Kruskal Wallis $H_{(2)}$ =53.03 **p<0.0001** | | | |
| ElavGal4;MBG80>Nf1RNAi Dcr-2(Day) | 13.88 ± 1.1 | 60 | |
| ElavGal4;MBG80/+ (Day) | 18.19 ± 0.93 | 43 | **<0.0001** |
| Nf1RNAi; Dcr-2/+ (Day) | 26.00 ± 0.80 | 45 | 0.0369 |
| Kruskal Wallis $H_{(2)}$ =88.91 **p<0.0001** | | | |
| ElavGal4; MBG80>Nf1RNAi; Dcr-2(Night) | 3.808 ± 0.65 | 60 | |
| ElavGal4; MBG80/+ (Night) | 18.38 ± 0.93 | 43 | **<0.0001** |
| Nf1RNAi; Dcr-2/+ (Night) | 17.43 ± 0.63 | 45 | **<0.0001** |
| **Fig. S3I** Kruskal Wallis $H_{(2)}$ =61.56 **p<0.0001** | | | |
| ElavGal4;MBG80>Nf1RNAi; Dcr-2(Day) | 10.11 ± 0.76 | 60 | |
| ElavGal4;MBG80/+ (Day) | 22.67 ± 1.9 | 43 | **<0.0001** |

| | | | |
|---|---|---|---|
| Nf1RNAi; Dcr-2/+ (Day) | 16.58 ± 0.62 | 45 | **<0.0001** |
| Kruskal Wallis $H_{(2)}$ =53.97 **p<0.0001** | | | |
| ElavGal4;MBG80>Nf1RNAi Dcr-2 (Night) | 12.47 ± 2.0 | 60 | |
| ElavGal4;MBG80/+ (Night) | 36.51 ± 4.9 | 43 | **<0.0001** |
| Nf1RNAi; Dcr-2/+ (Night) | 32.95 ± 2.1 | 45 | **<0.0001** |
| Mann-Whitney (U)=509 **p=0.0003** | | | |
| ElavGal4;MBG80/+ (Day) | 22.67 ± 1.9 | 43 | |
| ElavGal4;MBG80/+ (Night) | 36.51 ± 4.9 | 43 | **0.0003** |
| Mann-Whitney (U)=82 **p<0.0001** | | | |
| Nf1RNAi; Dcr-2/+ (Day) | 16.58 ± 0.62 | 45 | |
| Nf1RNAi; Dcr-2/+ (Night) | 32.95 ± 2.1 | 45 | **<0.0001** |
| Mann-Whitney (U)=1094 p=0.0143 | | | |
| ElavGal4;MBG80>Nf1RNAi ;UASDcr-2(Day) | 10.11 ± 0.76 | 60 | |
| ElavGal4;MBG80>Nf1RNAi ;UASDcr-2 (Night) | 12.47 ± 2.0 | 60 | 0.0143 |
| **Fig. S4B** Kruskal Wallis $H_{(2)}$ =165.1 **p<0.0001** | | | |
| ElavGal4/+ | 859.3 ± 16 | 86 | |
| UASdNf1; Nf1Null | 260.0 ± 13 | 83 | **<0.0001** |
| ElavGal4>UASdNf1; $Nf1^{Null}$ | 771.5 ± 23 | 96 | 0.1100 |
| ElavGal4>UASdNf1; Nf1Null | 771.5 ± 23 | 96 | |
| UASdNf1; Nf1Null | 260.0 ± 13 | 83 | **<0.0001** |
| **Fig. S4C** Kruskal Wallis $H_{(2)}$ =132.3 **p<0.0001** | | | |
| ElavGal4/+(Day) | 349.1 ± 11 | 86 | |
| UASdNf1; Nf1Null (Day) | 123.8 ± 8.2 | 85 | **<0.0001** |
| ElavGal4>UASdNf1; Nf1Null (Day) | 354.4 ± 13 | 96 | >0.9999 |
| ElavGal4>UASdNf1; Nf1Null (Day) | 354.4 ± 13 | 96 | |
| UASdNf1; Nf1Null (Day) | 123.8 ± 8.2 | 85 | **<0.0001** |
| Kruskal Wallis $H_{(2)}$ = 160.9 **p<0.0001** | | | |
| ElavGal4/+(Night) | 509.8 ± 7.3 | 86 | |
| UASdNf1; Nf1Null (Night) | 148.3 ± 9.1 | 85 | **<0.0001** |
| ElavGal4>UASdNf1; Nf1Null (Night) | 416.6 ± 15 | 96 | **0.0010** |
| ElavGal4>UASdNf1; Nf1Null (Night) | 416.6 ± 15 | 96 | |
| UASdNf1; Nf1Null (Night) | 148.3 ± 9.1 | 85 | **<0.0001** |
| **Fig. S4D** Kruskal Wallis $H_{(2)}$ =55.73 **p<0.0001** | | | |
| ElavGal4/+(Day) | 20.93 ± 0.57 | 86 | |
| UASdNf1; Nf1Null (Day) | 15.31 ± 0.67 | 85 | **<0.0001** |

| | | | |
|---|---|---|---|
| ElavGal4>UASdNf1; Nf1Null (Day) | 22.38 ± 0.59 | 92 | 0.4449 |
| ElavGal4>UASdNf1; Nf1Null (Day) | 22.38 ± 0.59 | 92 | |
| UASdNf1; Nf1Null (Day) | 15.31 ± 0.67 | 85 | **<0.0001** |
| Kruskal Wallis $H_{(2)}$ =80.60 **p<0.0001** | | | |
| ElavGal4/+(Night) | 18.98 ± 0.52 | 86 | |
| UASdNf1; Nf1Null (Night) | 17.25 ± 0.80 | 85 | 0.4589 |
| ElavGal4>UASdNf1; Nf1Null (Night) | 27.62 ± 0.76 | 94 | **<0.0001** |
| ElavGal4>UASdNf1; Nf1Null (Night) | 27.62 ± 0.76 | 94 | |
| UASdNf1; Nf1Null (Night) | 17.25 ± 0.80 | 85 | **<0.0001** |
| **Fig. S4E** Kruskal Wallis $H_{(2)}$ = 144.9 **p<0.0001** | | | |
| ElavGal4/+(Day) | 18.25 ± 1.1 | 86 | |
| UASdNf1; Nf1Null (Day) | 7.839 ± 0.38 | 85 | **<0.0001** |
| ElavGal4>UASdNf1; Nf1Null (Day) | 17.14 ± 0.87 | 96 | >0.9999 |
| ElavGal4>UASdNf1; Nf1Null (Day) | 17.14 ± 0.87 | 96 | |
| UASdNf1; Nf1Null (Day) | 7.839 ± 0.38 | 85 | **<0.0001** |
| Kruskal Wallis $H_{(2)}$ = 179.0 **p<0.0001** | | | |
| ElavGal4/+(Night) | 28.27 ± 0.97 | 86 | |
| UASdNf1; Nf1Null (Night) | 8.986 ± 0.68 | 85 | **<0.0001** |
| ElavGal4>UASdNf1; Nf1Null (Night) | 16.14 ± 0.87 | 96 | **<0.0001** |
| ElavGal4>UASdNf1; Nf1Null (Night) | 16.14 ± 0.87 | 96 | |
| UASdNf1; Nf1Null (Night) | 8.986 ± 0.68 | 85 | **<0.0001** |
| **Fig. S5** ANOVA $F_{(3,32)}$ = 1.061 p=0.3794 | | | |
| w1118 | 1.00 ± 0 | 9 | |
| $Nf1^{Null}$ | 0.8973 ± 0.057 | 9 | 0.6368 |
| $Nf1^{C1045Y}$ | 1.007 ± 0.058 | 9 | 0.9778 |
| $Nf1^{R1861C}$ | 1.029 ± 0.080 | 9 | 0.9778 |
| **Fig. S6A** Kruskal Wallis $H_{(2)}$ =251.7 **p<0.0001** | | | |
| ElavGal4/+ | 835.5 ± 11 | 157 | |
| UASdNf1-GRD; Nf1Null | 315.1 ± 14 | 164 | **<0.0001** |
| ElavGal4>UASdNf1-GRD; $Nf1^{Null}$ | 624.7 ± 25 | 109 | **<0.0001** |
| ElavGal4>UASdNf1-GRD; $Nf1^{Null}$ | 624.7 ± 25 | 109 | |
| UASdNf1-GRD; Nf1Null | 315.1 ± 14 | 164 | **<0.0001** |

| | | | |
|---|---|---|---|
| **Fig. S6B** Kruskal Wallis $H_{(2)} = 255.6$ **p<0.0001** | | | |
| ElavGal4/+ (Day) | 376.6 ± 6.3 | 157 | |
| UASdNf1-GRD; Nf1Null (Day) | 111.5 ± 6.6 | 164 | **<0.0001** |
| ElavGal4>UASdNf1-GRD; Nf1Null (Day) | 316.6 ± 13 | 109 | **0.0030** |
| ElavGal4>UASdNf1-GRD; Nf1Null (Day) | 316.6 ± 13 | 109 | |
| UASdNf1-GRD; Nf1Null (Day) | 111.5 ± 6.6 | 164 | **<0.0001** |
| Kruskal Wallis $H_{(2)} = 222.4$ **p<0.0001** | | | |
| ElavGal4/+ (Night) | 458.4 ± 6.2 | 157 | |
| UASdNf1-GRD; Nf1Null (Night) | 203.4 ± 8.5 | 164 | **<0.0001** |
| ElavGal4>UASdNf1-GRD; Nf1Null (Night) | 308.1 ± 14 | 109 | **<0.0001** |
| ElavGal4>UASdNf1-GRD; Nf1Null (Night) | 308.1 ± 14 | 109 | |
| UASdNf1-GRD; Nf1Null (Night) | 203.4 ± 8.5 | 164 | **<0.0001** |
| **Fig. S6C** ANOVA $F_{(2,427)} = 88.76$ **p<0.0001** | | | |
| ElavGal4/+ (Day) | 21.60 ± 0.37 | 157 | |
| UASdNf1-GRD; Nf1Null (Day) | 13.34 ± 0.48 | 164 | **<0.0001** |
| ElavGal4>UASdNf1-GRD; $Nf1^{Null}$ (Day) | 17.05 ± 0.55 | 109 | **<0.0001** |
| ElavGal4>UASdNf1-GRD; $Nf1^{Null}$ (Day) | 17.05 ± 0.55 | 109 | |
| UASdNf1-GRD; Nf1Null (Day) | 13.34 ± 0.48 | 164 | **<0.0001** |
| ANOVA $F_{(2,427)} = 1.808$ p=0.1652 | | | |
| ElavGal4/+ (Night) | 20.82 ± 0.39 | 157 | |
| UASdNf1-GRD; Nf1Null (Night) | 19.87 ± 0.56 | 164 | 0.3667 |
| ElavGal4>UASdNf1-GRD; $Nf1^{Null}$ (Night) | 19.40 ± 0.63 | 109 | 0.1684 |
| ElavGal4>UASdNf1-GRD; $Nf1^{Null}$ (Night) | 19.40 ± 0.63 | 109 | |
| UASdNf1-GRD; Nf1Null (Night) | 19.87 ± 0.56 | 164 | 0.8198 |
| **Fig. S6D** Kruskal Wallis $H_{(2)} = 264.5$ **p<0.0001** | | | |
| ElavGal4/+ (Day) | 19.22 ± 1.2 | 157 | |
| UASdNf1-GRD; $Nf1^{Null}$ (Day) | 8.442 ± 0.85 | 163 | **<0.0001** |
| ElavGal4>UASdNf1-GRD; Nf1Null (Day) | 21.06 ± 1.4 | 109 | >0.9999 |
| ElavGal4>UASdNf1-GRD; $Nf1^{Null}$ (Day) | 21.06 ± 1.4 | 109 | |
| UASdNf1-GRD; $Nf1^{Null}$ (Day) | 8.442 ± 0.85 | 163 | **<0.0001** |
| Kruskal Wallis $H_{(2)} = 222.0$ **p<0.0001** | | | |

| | | | |
|---|---|---|---|
| ElavGal4/+ (Night) | 24.02 ± 0.98 | 157 | |
| UASdNf1-GRD; Nf1Null (Night) | 9.985 ± 0.37 | 163 | **<0.0001** |
| ElavGal4>UASdNf1-GRD; $Nf1^{Null}$ (Night) | 18.34 ± 1.7 | 109 | **<0.0001** |
| ElavGal4>UASdNf1-GRD; $Nf1^{Null}$ (Night) | 18.34 ± 1.7 | 109 | |
| UASdNf1-GRD; $Nf1^{Null}$ (Night) | 9.985 ± 0.37 | 163 | **<0.0001** |
| **Fig. S7B** Kruskal Wallis $H_{(3)}$ = 32.62 **p<0.0001** | | | |
| $w^{1118}$ | 806.0 ± 20 | 35 | |
| $Nf1^{Null/+}$ | 924.0 ± 14 | 49 | **<0.0001** |
| $Nf1^{C1045Y/+}$ | 921.2 ± 17 | 49 | **<0.0001** |
| $Nf1^{R1861C/+}$ | 873.3 ± 17 | 104 | 0.0661 |
| $Nf1^{C1045Y/+}$ | 921.2 ± 17 | 49 | |
| $Nf1^{R1861C/+}$ | 873.3 ± 17 | 104 | **0.0073** |
| $Nf1^{Null/+}$ | 924.0 ± 14 | 49 | |
| $Nf1^{C1045Y/+}$ | 921.2 ± 17 | 49 | >0.9999 |
| $Nf1^{R1861C/+}$ | 873.3 ± 17 | 104 | 0.0114 |
| **Fig. S7C** Kruskal Wallis $H_{(3)}$ = 136.7 **p<0.0001** | | | |
| $w^{1118}$ (Day) | 288.0 ± 15 | 35 | |
| $Nf1^{Null/+}$ (Day) | 369.3 ± 11 | 49 | 0.0621 |
| $Nf1^{C1045Y/+}$ (Day) | 367.7 ± 9.4 | 49 | 0.0789 |
| $Nf1^{R1861C/+}$ (Day) | 493.5 ± 7.2 | 104 | **<0.0001** |
| $Nf1^{C1045Y/+}$ (Day) | 367.7 ± 9.4 | 49 | |
| $Nf1^{R1861C/+}$ (Day) | 493.5 ± 7.2 | 104 | **<0.0001** |
| $Nf1^{Null/+}$ (Day) | 369.3 ± 11 | 49 | |
| $Nf1^{C1045Y/+}$ (Day) | 367.7 ± 9.4 | 49 | >0.9999 |
| $Nf1^{R1861C/+}$ (Day) | 493.5 ± 7.2 | 104 | **<0.0001** |
| Kruskal Wallis $H_{(3)}$ = 104.7 **p<0.0001** | | | |
| $w^{1118}$ (Night) | 517.6 ± 9.9 | 35 | |
| $Nf1^{Null/+}$ (Night) | 554.4 ± 6.6 | 49 | 0.1486 |

| | | | |
|---|---|---|---|
| Nf1$^{C1045Y/+}$ (Night) | 553.0 ± 10 | 49 | 0.1552 |
| Nf1$^{R1861C/+}$ (Night) | 379.0 ± 12 | 104 | **<0.0001** |
| Nf1$^{C1045Y/+}$ (Night) | 553.0 ± 10 | 49 | |
| Nf1$^{R1861C/+}$ (Night) | 379.0 ± 12 | 104 | **<0.0001** |
| Nf1$^{Null/+}$ (Night) | 554.4 ± 6.6 | 49 | |
| Nf1$^{C1045Y/+}$ (Night) | 553.0 ± 10 | 49 | >0.9999 |
| Nf1$^{R1861C/+}$ (Night) | 379.0 ± 12 | 104 | **<0.0001** |
| **Fig. S7D** Kruskal Wallis $H_{(3)}$ = 27.40 **p<0.0001** | | | |
| w$^{1118}$ (Day) | 24.84 ± 0.76 | 35 | |
| Nf1$^{Null/+}$ (Day) | 25.38 ± 0.76 | 49 | >0.9999 |
| Nf1$^{C1045Y/+}$ (Day) | 28.91 ± 0.61 | 49 | **0.0004** |
| Nf1$^{R1861C/+}$ (Day) | 24.87 ± 0.44 | 104 | >0.9999 |
| Nf1$^{C1045Y/+}$ (Day) | 28.91 ± 0.61 | 49 | |
| Nf1$^{R1861C/+}$ (Day) | 24.87 ± 0.44 | 104 | **<0.0001** |
| Nf1$^{Null/+}$ (Day) | 25.38 ± 0.76 | 49 | |
| Nf1$^{C1045Y/+}$ (Day) | 28.91 ± 0.61 | 49 | **0.0014** |
| Nf1$^{R1861C/+}$ (Day) | 24.87 ± 0.44 | 104 | >0.9999 |
| Kruskal Wallis $H_{(3)}$ = 19.00 **p=0.0003** | | | |
| w$^{1118}$ (Night) | 23.51 ± 0.81 | 35 | |
| Nf1$^{Null/+}$ (Night) | 19.30 ± 0.77 | 48 | **0.0006** |
| Nf1$^{C1045Y/+}$ (Night) | 18.72 ± 0.98 | 49 | **0.0008** |
| Nf1$^{R1861C/+}$ (Night) | 19.98 ± 0.56 | 104 | **0.0041** |
| Nf1$^{C1045Y/+}$ (Night) | 18.72 ± 0.98 | 49 | |
| Nf1$^{R1861C/+}$ (Night) | 19.98 ± 0.56 | 104 | >0.9999 |
| Nf1$^{Null/+}$ (Night) | 19.30 ± 0.77 | 48 | |
| Nf1$^{C1045Y/+}$ (Night) | 18.72 ± 0.98 | 49 | >0.9999 |
| Nf1$^{R1861C/+}$ (Night) | 19.98 ± 0.56 | 104 | >0.9999 |
| **Fig. S7E** Kruskal Wallis $H_{(3)}$ = 116.9 **p<0.0001** | | | |
| w$^{1118}$ (Day) | 11.46 ± 0.45 | 35 | |
| Nf1$^{Null/+}$ (Day) | 15.14 ± 0.65 | 49 | **0.0030** |
| Nf1$^{C1045Y/+}$ (Day) | 12.85 ± 0.36 | 49 | 0.9570 |
| Nf1$^{R1861C/+}$ (Day) | 20.58 ± 0.55 | 104 | **<0.0001** |
| Nf1$^{C1045Y/+}$ (Day) | 12.85 ± 0.36 | 49 | |
| Nf1$^{R1861C/+}$ (Day) | 20.58 ± 0.55 | 104 | **<0.0001** |
| Nf1$^{Null/+}$ (Day) | 15.14 ± 0.65 | 49 | |
| Nf1$^{C1045Y/+}$ (Day) | 12.85 ± 0.36 | 49 | 0.1372 |

| | | | |
|---|---|---|---|
| $Nf1^{R1861C/+}$ (Day) | 20.58 ± 0.55 | 104 | **<0.0001** |
| Kruskal Wallis $H_{(3)}$ = 67.70 **p<0.0001** | | | |
| $w^{1118}$ (Night) | 21.57 ± 0.82 | 33 | |
| $Nf1^{Null/+}$ (Night) | 30.94 ± 1.3 | 48 | **0.0029** |
| $Nf1^{C1045Y/+}$ (Night) | 33.22 ± 2.5 | 47 | 0.0139 |
| $Nf1^{R1861C/+}$ (Night) | 21.66 ± 1.6 | 104 | 0.2120 |
| $Nf1^{C1045Y/+}$ (Night) | 33.22 ± 2.5 | 47 | |
| $Nf1^{R1861C/+}$ (Night) | 21.66 ± 1.6 | 104 | **<0.0001** |
| $Nf1^{Null/+}$ (Night) | 30.94 ± 1.3 | 48 | |
| $Nf1^{C1045Y/+}$ (Night) | 33.22 ± 2.5 | 47 | >0.9999 |
| $Nf1^{R1861C/+}$ (Night) | 21.66 ± 1.6 | 104 | **<0.0001** |
| **Fig. S7F** Kruskal Wallis $H_{(3)}$ = 99.16 **p<0.0001** | | | |
| $w^{1118}$ (Day) | 5848 ± 330 | 35 | |
| $Nf1^{Null/+}$ (Day) | 5618 ± 300 | 48 | >0.9999 |
| $Nf1^{C1045Y/+}$ (Day) | 4114 ± 180 | 49 | 0.0126 |
| $Nf1^{R1861C/+}$ (Day) | 2746 ± 140 | 104 | **<0.0001** |
| $Nf1^{C1045Y/+}$ (Day) | 4114 ± 180 | 49 | |
| $Nf1^{R1861C/+}$ (Day) | 2746 ± 140 | 104 | **<0.0001** |
| $Nf1^{Null/+}$ (Day) | 5618 ± 300 | 48 | |
| $Nf1^{C1045Y/+}$ (Day) | 4114 ± 180 | 49 | 0.0256 |
| $Nf1^{R1861C/+}$ (Day) | 2746 ± 140 | 104 | **<0.0001** |
| Kruskal Wallis $H_{(3)}$ =65.48 **p<0.0001** | | | |
| $w^{1118}$ (Night) | 2702 ± 210 | 35 | |
| $Nf1^{Null/+}$ (Night) | 3123 ± 170 | 48 | >0.9999 |
| $Nf1^{C1045Y/+}$ (Night) | 2848 ± 190 | 49 | >0.9999 |
| $Nf1^{R1861C/+}$ (Night) | 5984 ± 330 | 102 | **<0.0001** |
| $Nf1^{C1045Y/+}$ (Night) | 2848 ± 190 | 49 | |
| $Nf1^{R1861C/+}$ (Night) | 5984 ± 330 | 102 | **<0.0001** |
| $Nf1^{Null/+}$ (Night) | 3123 ± 170 | 48 | |
| $Nf1^{C1045Y/+}$ (Night) | 2848 ± 190 | 49 | >0.9999 |
| $Nf1^{R1861C/+}$ (Night) | 5984 ± 330 | 102 | **<0.0001** |
| **Fig. S7G** Kruskal Wallis $H_{(3)}$ =85.45 **p<0.0001** | | | |
| $w^{1118}$ (Day) | 5.004 ± 0.25 | 35 | |
| $Nf1^{Null/+}$ (Day) | 5.203 ± 0.25 | 48 | >0.9999 |
| $Nf1^{C1045Y/+}$ (Day) | 3.799 ± 0.14 | 49 | 0.0166 |

| | | | |
|---|---|---|---|
| $Nf1^{R1861C/+}$ (Day) | 2.838 ± 0.12 | 104 | **<0.0001** |
| $Nf1^{C1045Y/+}$ (Day) | 3.799 ± 0.14 | 49 | |
| $Nf1^{R1861C/+}$ (Day) | 2.838 ± 0.12 | 104 | **0.0006** |
| $Nf1^{Null/+}$ (Day) | 5.203 ± 0.25 | 48 | |
| $Nf1^{C1045Y/+}$ (Day) | 3.799 ± 0.14 | 49 | **0.0035** |
| $Nf1^{R1861C/+}$ (Day) | 2.838 ± 0.12 | 104 | **<0.0001** |
| Kruskal Wallis $H_{(3)}$ =50.36 **p<0.0001** | | | |
| $w^{1118}$ (Night) | 2.881 ± 0.21 | 35 | |
| $Nf1^{Null/+}$ (Night) | 3.514 ± 0.18 | 48 | 0.5981 |
| $Nf1^{C1045Y/+}$ (Night) | 3.145 ± 0.17 | 49 | >0.9999 |
| $Nf1^{R1861C/+}$ (Night) | 5.271 ± 0.26 | 104 | **<0.0001** |
| $Nf1^{C1045Y/+}$ (Night) | 3.145 ± 0.17 | 49 | |
| $Nf1^{R1861C/+}$ (Night) | 5.271 ± 0.26 | 104 | **<0.0001** |
| $Nf1^{Null/+}$ (Night) | 3.514 ± 0.18 | 48 | |
| $Nf1^{C1045Y/+}$ (Night) | 3.145 ± 0.17 | 49 | **0.0005** |
| $Nf1^{R1861C/+}$ (Night) | 5.271 ± 0.26 | 104 | **<0.0001** |
| **Fig. S8A** ANOVA $F_{(2,15)}$ = 35.33 **p<0.0001** | | | |
| $w^{1118}$ | 1.00 ± 0 | 6 | |
| $Nf1^{C1045Y}$ | 1.249 ± 0.021 | 6 | **<0.0001** |
| $Nf1^{R1861C}$ | 1.152 ± 0.030 | 6 | **0.0003** |
| $Nf1^{C1045Y}$ | 1.249 ± 0.021 | 6 | |
| $Nf1^{R1861C}$ | 1.152 ± 0.030 | 6 | **0.0052** |
| **Fig. S8B** Kruskal Wallis $_{(2)}$ = 82.93 **p<0.0001** | | | |
| ElavGal4/+ (Day) | 4625 ± 180 | 92 | |
| UASdNf1; $Nf1^{C1045Y}$(Day) | 2819 ± 95 | 80 | **<0.0001** |
| ElavGal4>UASdNf1; $Nf1^{C1045Y}$ (Day) | 4863 ± 160 | 136 | >0.9999 |
| ElavGal4>UASdNf1; $Nf1^{C1045Y}$ (Day) | 4863 ± 160 | 136 | |
| UASdNf1; $Nf1^{C1045Y}$ (Day) | 2819 ± 95 | 80 | **<0.0001** |
| Kruskal Wallis $_{(2)}$ = 117.6 **p<0.0001** | | | |
| ElavGal4/+ (Night) | 4409 ± 200 | 91 | |

| | | | |
|---|---|---|---|
| UASdNf1; Nf1$^{C1045Y}$ (Night) | 2835 ± 130 | 81 | **<0.0001** |
| ElavGal4>UASdNf1; Nf1$^{C1045Y}$ (Night) | 6471 ± 220 | 134 | **<0.0001** |
| ElavGal4>UASdNf1; Nf1$^{C1045Y}$ (Night) | 6471 ± 220 | 134 | |
| UASdNf1; Nf1$^{C1045Y}$ (Night) | 2835 ± 130 | 81 | **<0.0001** |
| Mann-Whitney (U)=3837 p=0.3305 | | | |
| ElavGal4/+ (Day) | 4625 ± 180 | 92 | |
| ElavGal4/+ (Night) | 4409 ± 200 | 91 | 0.3305 |
| Mann-Whitney (U)=5582 **p<0.0001** | | | |
| ElavGal4>UASdNf1; Nf1$^{C1045Y}$(Day) | 4863 ± 160 | 136 | |
| ElavGal4>UASdNf1; Nf1$^{C1045Y}$ (Night) | 6471 ± 220 | 134 | **<0.0001** |
| **Fig. S8C** Kruskal Wallis $_{(2)}$ = 132.9 **p<0.0001** | | | |
| ChaGal4/+ (Day) | 5359 ± 210 | 95 | |
| UASdNf1; Nf1$^{C1045Y}$ (Day) | 3814 ± 190 | 97 | **<0.0001** |
| ChaGal4>UASdNf1; Nf1$^{C1045Y}$ (Day) | 7172 ± 190 | 148 | **<0.0001** |
| ChaGal4>UASdNf1; Nf1$^{C1045Y}$ (Day) | 7172 ± 190 | 148 | |
| UASdNf1; Nf1$^{C1045Y}$ (Day) | 3814 ± 190 | 97 | **<0.0001** |
| Kruskal Wallis $_{(2)}$ = 57.07 **p<0.0001** | | | |
| ChaGal4/+ (Night) | 3001 ± 170 | 95 | |
| UASdNf1; Nf1$^{C1045Y}$ (Night) | 3275 ± 300 | 97 | >0.9999 |
| ChaGal4>UASdNf1; Nf1$^{C1045Y}$ (Night) | 4601 ± 170 | 148 | **<0.0001** |
| ChaGal4>UASdNf1; Nf1$^{C1045Y}$ (Night) | 4601 ± 170 | 148 | |
| UASdNf1; Nf1$^{C1045Y}$ (Night) | 3275 ± 300 | 97 | **<0.0001** |
| Mann-Whitney (U)= 1653 **p<0.0001** | | | |
| ChaGal4/+ (Day) | 5359 ± 210 | 95 | |
| ChaGal4/+ (Night) | 3001 ± 170 | 95 | **<0.0001** |
| Mann-Whitney (U)= 4293 **p<0.0001** | | | |
| ChaGal4>UASdNf1; Nf1$^{C1045Y}$ (Day) | 7172 ± 190 | 148 | |
| ChaGal4>UASdNf1; Nf1$^{C1045Y}$ (Night) | 4601 ± 170 | 148 | **<0.0001** |
